# Stepwise and lineage-specific divergence of a major immune co-chaperone complex in leptosporangiate ferns

**DOI:** 10.64898/2025.12.17.694876

**Authors:** Hyeon-Min Jeong, Yu Sugihara, Michael W. Webster, Philip Carella

## Abstract

Protein-protein interactions are essential for proper cellular function and are often under strong evolutionary pressures to maintain their stability and specificity. In plants, a broadly distributed chaperone complex comprised of the RAR1 (REQUIRED FOR MLA12 RESISTANCE 1) and SGT1 (SUPRESSOR OF THE G2 ALLELE OF SKP1) co-chaperones alongside the HSP90 (HEAT SHOCK PROTEIN90) core chaperone are important for immunity. Despite its importance in flowering plants, a deeper understanding of how this complex evolved remains limited. Here, we examine the molecular evolutionary history of the RAR1-SGT1 interaction across land plants. We identified a lineage-specific divergence of the RAR1-SGT1 binding interface in vascular non-seed ferns, which renders orthologs unable to interact outside of their lineage. Further investigation of interface diversity uncovered a single amino acid residue in RAR1 and three corresponding residues in SGT1 that collectively dictate binding specificity. Ancestral state reconstruction supported stepwise evolution of specificity in SGT1 in leptopsporangiate ferns, which was initiated by a promiscuous intermediate state that widened its capacity to bind RAR1 before subsequent mutations locked in specificity. Our data highlight the broad conservation of the RAR1-SGT1 interface and the coevolutionary dynamics that shaped interface maintenance during lineage-specific diversification.

## INTRODUCTION

Protein-protein interactions are essential to cellular processes and the interfaces that mediate these interactions are under strict selection pressures. The complex interaction networks within a given cellular compartment requires specificity to ensure that proteins interact with necessary partners. This requirement drives the molecular coevolution of interacting proteins, where it is essential to maintain important protein-protein interactions as organisms diversify (Lovell and Robertson 2010). Indeed, coevolving residues are predominantly located at binding interfaces (Liu et al. 2007; Yeang and Haussler 2007; Skerker et al. 2008). Understanding the coevolutionary dynamics of protein-protein interaction is therefore an important facet of our mechanistic understanding of biological systems.

The robustness of organisms against environmental perturbations is supported by the buffering capacity by chaperones (Jarosz et al. 2010), which are broadly classified into two categories: core chaperones and co-chaperones. Core chaperones assist client proteins to reach and maintain their required conformations by mediating correct folding (Kim et al. 2013). Co-chaperones bind to core chaperones as non-clients and direct their buffering capacity toward specific cellular processes. This occurs through different modes of action, including chaperone-client coupling or by regulating the ATPase cycles of core chaperones (Caplan 2003; Jackson et al. 2004; Shirasu 2009; Taipale et al. 2010).

The co-chaperones RAR1 (REQUIRED FOR MLA12 RESISTANCE 1) and SGT1 (SUPPRESSOR OF THE G2 ALLELE OF SKP1) function alongside the core chaperone HSP90 (HEAT SHOCK PROTEIN 90) and are conserved across eukaryotes (Kitagawa et al. 1999; Azevedo et al. 2002; Shirasu 2009). In flowering plants, these proteins form an immune chaperone complex that maintains the steady-state accumulation of intracellular nucleotide-binding leucine-rich repeat (NLR) immune receptors (Shirasu 2009). Accordingly, *rar1* and *sgt1* mutants are often impaired in immunity, showing enhanced disease susceptibility to downy mildew and bacteria in *Arabidopsis thaliana* (Austin et al. 2002; Muskett et al. 2002; Tör et al. 2002), powdery mildew in barley (Shirasu et al. 1999; Azevedo et al. 2002), and tobacco mosaic virus in tobacco (Liu, Schiff, Marathe, et al. 2002; Liu, Schiff, Serino, et al. 2002).

The RAR1-SGT1-HSP90 immune chaperone complex is formed of two heterotrimers (6 total proteins), where specific domains underpin intermolecular interactions between each member (Zhang et al. 2010). RAR1 contains two CHORD domains (cysteine- and histidine-rich domain; CHORDI and CHORDII) that are separated by a short CCCH motif in plants. The CHORDII domain interacts with the CS (CHORD-containing protein and SGT1) domain of SGT1, which additionally contains TPR (tetratricopeptide repeat) and SGS (SGT1-specific) domains that are conserved across eukaryotes (Shirasu et al. 1999; Azevedo et al. 2002; Shirasu 2009). To complete the complex, the RAR1^CHORDII^ and SGT1^CS^ domains interact with distinct regions of the HSP90 N-terminus (HSP-N). Structural and biochemical investigations have resolved these binding interfaces, identifying conserved residues including RAR1^CHORDII^ isoleucine (I153) and SGT1^CS^ glutamate (E170) and tryptophan (W217) residues that are critical to maintain interactions between RAR1 and SGT1 (Azevedo et al. 2002; Zhang et al. 2010).

The RAR1-SGT1 interaction is conserved in flowering plants, as independent studies confirm protein-protein interactions of homologs encoded in Arabidopsis, rice, soybean, wheat, peanut, and banana (Liu, Schiff, Serino, et al. 2002; Tai 2008; Wang et al. 2008; Fu et al. 2009; Cantu et al. 2013; Yuan et al. 2021; Wang et al. 2024). The robustness of this interaction is exemplified by the use of AtRAR1 and AtSGT1 as benchmark controls for the development of novel *in planta* protein-protein interaction assays like the split-luciferase and split-GAL4 RUBY systems (Chen et al. 2008; Chen et al. 2023). However, we know little about the conservation of RAR1-SGT1 interactions across distantly related non-flowering plant lineages such as bryophytes, lycophytes, monilophytes (ferns), and gymnosperms. This is particularly important given the known role of RAR1-SGT1 in maintaining NLR-mediated immunity in flowering plants and the expanding diversity of these receptor types across non-flowering plant lineages (Andolfo et al. 2019; Chia et al. 2024; Castel et al. 2025).

Here, we uncover the evolutionary history and conservation of RAR1-SGT1 interactions in representative species spanning over 500 million years of plant evolution. We show that interactions are conserved within all tested species, and that the RAR1^CHORDII^-SGT1^CS^ binding interface unexpectedly diverged in a clade of ferns (vascular non-seed plants). Through the interrogation of extant fern diversity alongside ancestral state reconstruction, we reveal the coevolutionary history of RAR1-SGT1 interface diversification that occurred through the emergence of a promiscuous SGT1 intermediate with an expanded capacity to bind RAR1 orthologs. Collectively, our work highlights an evolutionary strategy that facilitated lineage-specific diversification while maintaining interface integrity of the broadly distributed RAR1-SGT1 co-chaperones.

## RESULTS

### RAR1 and SGT1 are conserved in land plants

Interactions between the RAR1 and SGT1 co-chaperones have been documented in several angiosperms (Azevedo et al. 2002; Liu, Schiff, Serino, et al. 2002; Tai 2008; Wang et al. 2008; Fu et al. 2009; Cantu et al. 2013); however, we know little about their evolutionary history and their capacity to interact within non-flowering lineages like bryophytes, lycophytes, monilophytes (ferns), or even extant streptophyte algae, which represent the sister lineage of land plants (Fig. 1A). To address this, we first identified RAR1 and SGT1 homologs from 364 species across diverse land plants and streptophyte algae using BLAST (basic local alignment search tool) and HMM (hidden Markov model) profiling. In total, we identified 294 RAR1 homologs from 294 species (6 algae and 288 land plants) and 395 SGT1 sequences from 328 species (5 algae and 323 land plants), respectively (Table S1). Consistent with the known duplication of SGT1 in flowering plants, we observed higher copy numbers of SGT1 in several species (Table S2), especially in angiosperms, whereas RAR1 exists as a single copy in plants (Table S1). Next, we performed phylogenetic analyses for RAR1 and SGT1, using algal species as outgroups. In both phylogenies, algae-rooted tree topologies generally aligned with known algal and land plant species relationships with some descrepancy observed in liverworts (Fig. 1B and C; Fig. S1; Fig. S2). Notably, the evolutionary trajectories of RAR1 and SGT1 differed in some lineages. For example, gymnosperm RAR1 and SGT1 homologs exhibit notably short branch lengths, which might reflect slower diversification rates in this lineage relative to other land plants. While mostly congruent with species phylogeny, these single gene trees illustrate the complex evolutionary relationships between bryophytes and land plants (Puttick et al. 2018; Sousa et al. 2019; Su et al. 2021; Harris et al. 2022; Chen and De Vries 2025; Dong et al. 2025).

**Figure 1.**
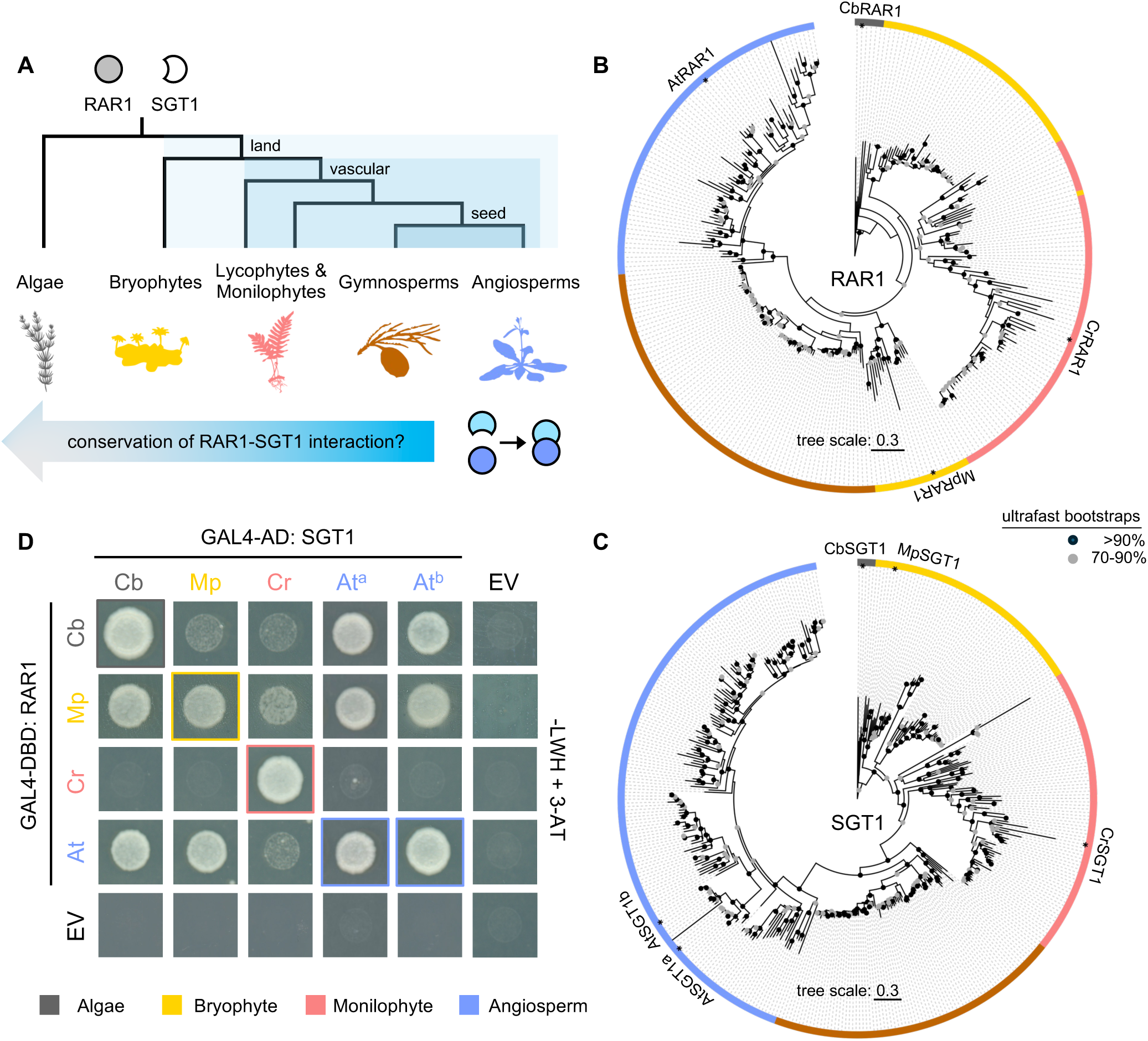
The RAR1-SGT1 interaction is conserved over more than 500 million years of plant evolution. **(A)** Schematic outlining plant evolutionary history with annotation depicting the known RAR1-SGT1 interaction in angiosperms and hypothesized interactions across distantly related lineages. The blue colour gradient indicates relative phylogenetic distance from angiosperms. **(B-C)** Maximum likelihood phylogenetic trees with 1000 ultrafast bootstrap replicates of **(B)** RAR1 and **(C)** SGT1 across land plants. Species used in yeast two-hybrid assays are indicated in each phylogeny. Trees were rooted using streptophyte algal sequences as the outgroup. Tree scale = substitutions per site. The colored ring around each tree indicates plant lineages; streptophyte algae (gray), bryophytes (yellow), lycophytes and monilophyte (pink), gymnosperms (brown), and angiosperms (blue) **(D)** Yeast two-hybrid protein-protein interaction assay with RAR1 orthologs fused to GAL4-DBD (DNA binding domain) and SGT1 orthologs fused to the GAL4-AD (activation domain). Orthologs were sourced from the alga *Chara braunii* (Cb, gray), the liverwort *Marchantia polymorpha* (Mp, yellow), the fern *Ceratopteris richardii* (Cr, pink), and the angiosperm *Arabidopsis thaliana* (At, blue). RAR1-SGT1 interaction testing in each species is highlighted by colour. At^a^ and At^b^ refer to AtSGT1a and AtSGT1b, respectively. Images were taken 3 days after plating and show yeast on SC media without Leu, Trp, and His (-LWH) and supplemented with 15 mM 3-AT (3-amino-1,2,4-triazole). “EV” indicates empty vector controls. This experiment was performed three times with similar results. Extended yeast two-hybrid data related to this figure are included in Data S1.

To assess the extent to which the RAR1-SGT1 interaction is conserved across diverse plants and algae, we performed yeast two-hybrid protein-protein interaction assays within and between homologs of the charophyte alga *Chara braunii,* the liverwort *Marchantia polymorpha*, the fern *Ceratopteris richardii*, and experimentally validated homologs from *Arabidopsis thaliana* (AtSGT1a and AtSGT1b). Yeast two-hybrid assays have previously been successful in identifying RAR1-SGT1 interactions for *A. thaliana* and other flowering plants (Azevedo et al. 2002; Liu, Schiff, Serino, et al. 2002; Wang et al. 2008; Fu et al. 2009; Cantu et al. 2013). Overall, yeast two-hybrid screening demonstrated that RAR1-SGT1 interactions were conserved, as the co-expression of each RAR1 and SGT1 pair from a given species produced clear growth of yeast colonies on interaction-specific media compared to empty vector (EV) controls that failed to interact (Fig. 1D; Data S1). RAR1-SGT1 interactions were also observed between homologs of distantly related species, including mixed RAR1-SGT1 pairs from the alga *C*. *braunii,* liverwort *M. polymorpha,* and angiosperm *A*. *thaliana*. However, CrRAR1 from the fern *C. richardii* interacted only with CrSGT1 and failed to interact with homologs from other lineages. Together, these results suggest that the RAR1-SGT1 interaction is highly conserved across land plants and algae and that their interaction interface has diverged in the fern *C. richardii*.

### A single mutation in the CrRAR1^CHORDII^ domain determines SGT1 specificity

In flowering plants, the RAR1-SGT1 interface is formed between the CHORDII domain of RAR1 and the CS domain of SGT1 (Azevedo et al. 2002; Zhang et al. 2010). We therefore reasoned that CrRAR1 specificity is caused by sequence variation within CHORDII and tested this idea using yeast two-hybrid assays by with CHORDII domain swaps of CrRAR1 and AtRAR1 (Fig. 2A). As anticipated, the wildtype RAR1-SGT1 controls from *C*. *richardii* and *A*. *thaliana* demonstrated strict interaction specificity, such that RAR1-SGT1 interactions were only observed between *A. thaliana* or *C. richardii* pairs. By contrast, CHORDII domain swapping resulted in reciprocal incompatibility, such that the CrRAR1^AtCHORDII^ now interacted only with AtSGT1b and AtRAR1^CrCHORDII^ interacted only with CrSGT1 (Fig. 2B; Data S1). This confirmed that interaction specificity is indeed determined by CHORDII domain identity.

**Figure 2.**
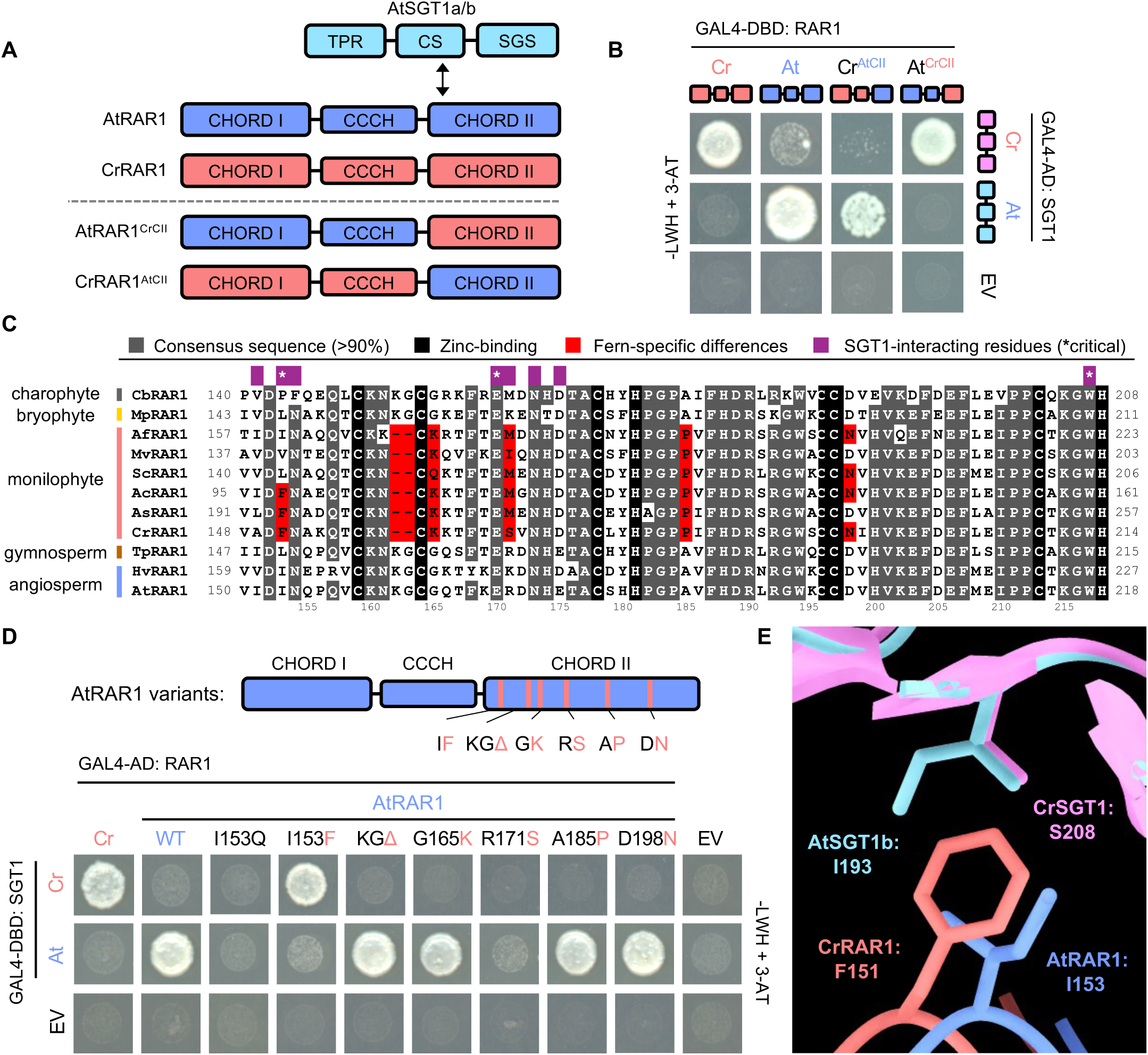
The CHORDII domain of RAR1 determines incompatibility between RAR1-SGT1 homologs in ferns and flowering plants. **(A)** A schematic depicting the known interaction between the AtRAR1-CHORDII and AtSGT1-CS domains is shown alongside wild-type chimeric RAR1 orthologs with swapped CHORDII domains. **(B)** Yeast two-hybrid assay with wild-type *Ceratopteris richardii* and *Arabidopsis thaliana* RAR1 orthologs (Cr and At) tested alongside CHORDII-swapped chimeras (Cr^AtCII^ and At^CrCII^) fused to GAL4-DBD (DNA binding domain) and SGT1 orthologs from *C. richardii* and *A. thaliana* fused to the GAL4-AD (activation domain. Images show yeast plated onto media without Leu, Trp, and His (-LWH) and supplemented with 15 mM 3-AT (3-aminotriazole). “EV” indicates empty vector controls. This experiment was performed three times with similar results. **(C)** Amino acid sequence alignment of the CHORDII domains of representative RAR1 homologs from charophytes (Cb, *Chara braunii*), bryophytes (Mp, *Marchantia polymorpha*), monilophytes (Af, *Azolla filiculoides*; Mv, *Marsilea vestita*; Sc, *Salvinia cucullata*; Ac, *Adiantum capillus-veneris*; As, *Alsophila spinulosa*; Cr, *C. richardii*), gymnosperms (Tp, *Thuja plicata*), and angiosperms (Hv, *Hordeum vulgare*; At, *Arabidopsis thaliana*). Fern-associated differences are colored in red and known SGT1-interacting residues of AtRAR1 are labelled with purple boxes. Asterisks indicate critical residues known to impact AtRAR1-AtSGT1a interactions. **(D)** Yeast two-hybrid assay investigating the impact of fern-specific amino acid variations introduced into the AtRAR1 backbone. RAR1 orthologs and variants are fused to the GAL4-DBD and were tested against *C. richardii* and *A. thaliana* SGT1 orthologs. Fern-specific differences are numbered based on corresponding positions in the AtRAR1 sequence. Images were taken 3 days after plating and show yeast on media without Leu, Trp, and His (-LWH) and supplemented with 15 mM 3-AT (3-amino-1,2,4-triazole). This experiment was performed times with similar results. **(E)** AlphaFold modelling of the RAR1-CHORD II and SGT1-CS binding interfaces from *A. thaliana* and *C. richardii.* The isoleucine (I) and phenylalanine (F) residues of the 153F variation, and the corresponding SGT1 amino acids, isoleucine (I) and serine (S) are depicted. Extended yeast two-hybrid data related to this figure are included in Data S1.

Next, we further analysed CHORDII domain sequences from representative plant and algal species, focusing on amino acid variation specific to ferns. Sequence alignments revealed six high-frequency variants observed within the CHORDII domain of ferns. We defined them as fern-associated differences, and designated as as Ile153Phe (I153F), Lys162Gly163Δ (KGΔ), Gly165Lys (G165K), Arg171Ser (R171S), Ala185Pro (A185P), and Asp198Asn (D198N) based on comparisons between AtRAR1 and CrRAR1 orthologs (Fig. 2C). The I153F and R171S residue variations occur within the known AtSGT1-binding interface (Zhang et al. 2010), while the four remaining fern-specific differences do not. Previous work has also shown that mutating Ile153, Glu170, and Trp217 impacts SGT1 binding capacity (Zhang et al. 2010). Unlike the diversified Ile153 residue, both Glu170 and Trp217 are conserved across distantly related RAR1 orthologs, including in CrRAR1. Amino acid residues associated with HSP90 interactions were conserved between ferns and other land plants (Fig. S3A), aside from a single A-to-P substitution in CHORDII (A185 in AtRAR1) that matches conserved proline at the equivalent site in CHORDI of all land plants (P38 in AtRAR1). This suggests that divergence within the CHORDII interface may be specifically associated with SGT1.

To understand the molecular basis of RAR1-SGT1 interaction specificity, we generated a series of AtRAR1 variants that incorporate each of the single fern-specific differences and tested them against fern and Arabidopsis SGT1. As an additional control, we included the AtRAR1^I153Q^ variant that is known to impair AtRAR1-AtSGT1a interactions (Zhang et al. 2010). As expected, wildtype AtRAR1 interacted with AtSGT1b in yeast whereas AtRAR1^I153Q^ and CrRAR1 failed to interact with AtSGT1b. Individual AtRAR1 variants carrying the fern-specific K162Δ2, G165K, A185P, and D198N all interacted with AtSGT1b and not CrSGT1, indicating that these residue positions do not contribute to specificity. By contrast, the I153F variant resulted in a full reversion of specificity, such that AtRAR1^I153F^ interacted with CrSGT1 instead of AtSGT1b (Fig. 2D; Data S1). To visualize variation at this position, we generated structural predictions of the AtRAR1-AtSGT1b and CrRAR1-CrSGT1 complexes using AlphaFold. Alignment of the two structural predictions shows that isoleucines of AtRAR1 (I153) and AtSGT1b (I193) are central to a hydrophobic interface (Zhang et al. 2010). By contrast, a bulky phenylalanine (CrRAR1:F151) and smaller serine (CrSGT1:S209) occupy the corresponding positions in the CrRAR1-CrSGT1 interface (Fig. 2E). Collectively, our data show that the transition from a isoleucine to a larger phenylalanine residue impacts RAR1-SGT1 specificity.

### The CrSGT1^CS^ domain harbors three fern-specific mutations to accommodate CrRAR1

To determine whether complementory changes in CrSGT1 evolved to accommodate variation in CrRAR1, we compared CS domain sequences of representative SGT1 orthologs and focused on sequence variation commonly observed within ferns. CS domain alignments revealed four fern-associated differences in SGT1 at the Glu191Ile (E191I), Ile193Ser (I193S), Ser195Lys (S195K) and Glu203Asn (E203N) positions (numbering based on AtSGT1b) (Fig. 3A). Among these differences, three residues (E191I, I193S, and S195K) are located within the AtRAR1-AtSGT1a binding interface (Zhang et al. 2010). Notably, previous work has shown that substitution of the Ile193 residue disrupts the RAR1-SGT1 interaction (Botër et al. 2007). An additional critical residue, Gly190, is conserved across land plants including ferns (Fig. 3A). All known HSP90-interacting residues were conserved in the SGT1 orthologs of land plants (Fig. S3B).

**Figure 3.**
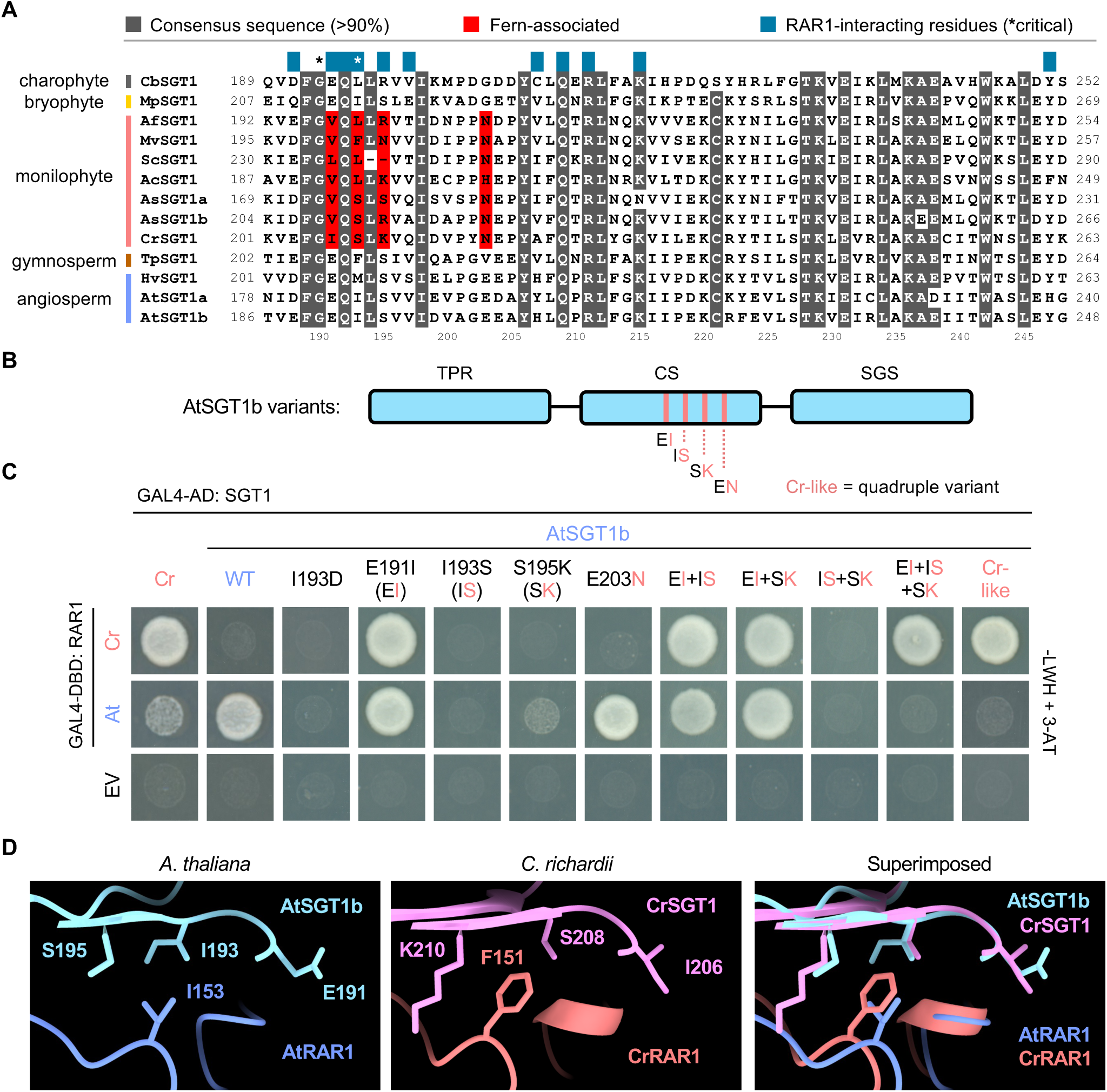
The *Ceratopteris richardii* SGT1-CS domain harbors compensatory mutations that enable interaction with RAR1. **(A)** Amino acid sequence alignment of CS domains from representative SGT1 homologs from charophytes (Cb, *Chara braunii*), bryophytes (Mp, *Marchantia polymorpha*), ferns (Af, *Azolla filiculoides*; Mv, *Marsilea vestita*; Sc, *Salvinia cucullata*; Ac, *Adiantum capillus-veneris*; As, *Alsophila spinulosa*; Cr, *C. richardii*), gymnosperms (Tp, *Thuja plicata*), and angiosperms (Hv, *Hordeum vulgare*; At, *Arabidopsis thaliana*). Fern-specific variations in SGT1 sequence are colored in red and the known RAR1-interacting residues of AtSGT1 are labelled with teal boxes. Asterisks indicate critical residues required for AtRAR1-AtSGT1a interactions that were tested in previous studies. **(B)** Schematic indicating the relative position of fern-specific variants within the CS domain of AtSGT1b. A variant carrying all 4 mutations is referred to as *Cr-like.* **(C)** Yeast-two hybrid assay using RAR1 homologs fused to the GAL4-DBD (DNA binding domain) tested against wildtype *C. richardii* SGT1, *A. thaliana* AtSGT1b, and AtSGT1b variants carrying combinations of fern-specific mutations indicated in (B). Images were taken 3 days after plating and show yeast on media without Leu, Trp, and His (-LWH) and supplemented with 15 mM 3-AT (3-amino-1,2,4-triazole). “EV” indicates empty vector controls. This experiment was performed three times with similar results. The full dilution series and additional interaction assay controls are presented in Data S1. **(D)** AlphaFold modelling of RAR1-SGT1 binding interface in *C*. *richardii* and *A. thaliana*. Models highlight all relevant amino acids as determined by yeast two-hybrid interaction assays.

To assess their contribution to interaction specificity, we generated AtSGT1b variants harbouring each amino acid variant (alone and in combination) (Fig. 3B) and examined their ability to interact with AtRAR1 or CrRAR1 by yeast two-hybrid. As an additional control, we included wild-type CrSGT1 and AtSGT1b, and the AtSGT1b^I193D^ variant that compromises interaction with AtRAR1 (Botër et al. 2007). Consistent with previous results, AtSGT1b and CrSGT1 interacted only with RAR1 orthologs from their own species, and the AtSGT1b^I193D^ negative control failed to interact with AtRAR1 (Fig. 3C; Data S1). We tested a quadruple variant of AtSGT1b carrying all four fern-associated differences that we termed ‘Cr-like’. As expected, the AtSGT1b^Cr-like^ quadruple variant interacted with CrRAR1 and not AtRAR1, demonstrating that interaction specificity is indeed determined by fern-associated sequence variation at these residues.

Single residue mutations were introduced into AtSGT1b to narrow down interaction specificity, however these displayed variable interaction profiles. Two fern-associated variants, I193S and S195K, disrupted interactions with both AtRAR1 and CrRAR1 when introduced into AtSGT1b, while the AtSGT1b^E203N^ variant had no impact on specificity. By contrast, AtSGT1b^E191I^ showed greater binding versatility and interacted with both AtRAR1 and CrRAR1. The most prevalent residue at this position across ferns is valine rather than isoleucine (Fig. S4), therefore we tested and confirmed that AtSGT1b^E191V^ similarly enables interaction with both AtRAR1 and CrRAR1 (Fig. S5; Data S1), reinforcing the idea that aliphatic residue at this position broadens binding capacity.

Since the interaction profile of AtSGT1b^E203N^ was identical to wildtype AtSGT1b, this mutation was excluded from further testing. To determine if the other three fern-associated variants (E191I, I193S, and S195K) could cooperatively reconstitute a fern-specific binding interface, we next evaluated them in double and triple combinations. When I193S or S195K mutations were combined with E191I, the resulting double mutant proteins remained as versatile as E191I, interacting with both AtRAR1 and CrRAR1. By comparison the I193S/S195K double mutant failed to interact with either RAR1 ortholog, consistent with the known impact of mutating I193 (Botër et al. 2007), Strikingly, combining all three mutations (E191I, I193S, and S195K) into a triple mutant protein fully recapitulated the interaction profile of CrSGT1, where it interacted specifically with CrRAR1 and not AtRAR1 (Fig. 3C, Data S1). This complete reversal in binding specificity demonstrates that all three CrSGT1 residues are collectively required for interaction specificity with CrRAR1.

Using AlphaFold, we mapped key residues associated with SGT1-RAR1 binding-specificity for both the *A. thaliana* and *C. richardii* pairs. This expanded view suggests that the amino acid residues impacting SGT1-RAR1 binding-specificity exist within close <u>proximity to</u> one another (Fig. 3D). In RAR1, the substitution of a small aliphatic isoleucine residue (*Arabidopsis*) with a bulky aromatic phenylalanine (*Ceratopteris*) is substantial change in size that impacts the physical space within the interface. Exactly how the three corresponsing residues in SGT1 accommodate this size difference is unknown and may benefit from future structural studies, however we observed substitutions of smaller aliphatic residues in fern SGT1 that likely accommodate the bulkier phenylalanine and enable RAR1-SGT1 interactions in ferns. In any event, our data collectively defines the molecular code underpinning RAR1-SGT1 interaction specificity that evolved in ferns like *C. richardii*.

### Four amino acids dictate RAR1-SGT1 binding specificity

To further support the idea that four fern-associated mutations provide RAR1-SGT1 binding specificity (three in SGT1 and one in RAR1), we performed yeast two-hybrid assays incorporating variants of all fern-associated changes in both RAR1 and SGT1 into Arabidopsis orthologs. CrRAR1 interacted with CrSGT1 and the triple AtSGT1b^EI+IS+SK^ variant but not wildtype AtSGT1b or the empty vector control (Fig. 4A; Data S1). By contrast, AtRAR1 interacted strongly with AtSGT1b and failed to interact with CrSGT1, AtSGT1b^EI+IS+SK^, or the empty vector control. Lastly, AtRAR1^I153F^ interacted with CrSGT1 and the fern-like AtSGT1b^EI+IS+SK^, but failed to interact with AtSGT1b or the empty vector control (Fig. 4A; Data S1).

**Figure 4.**
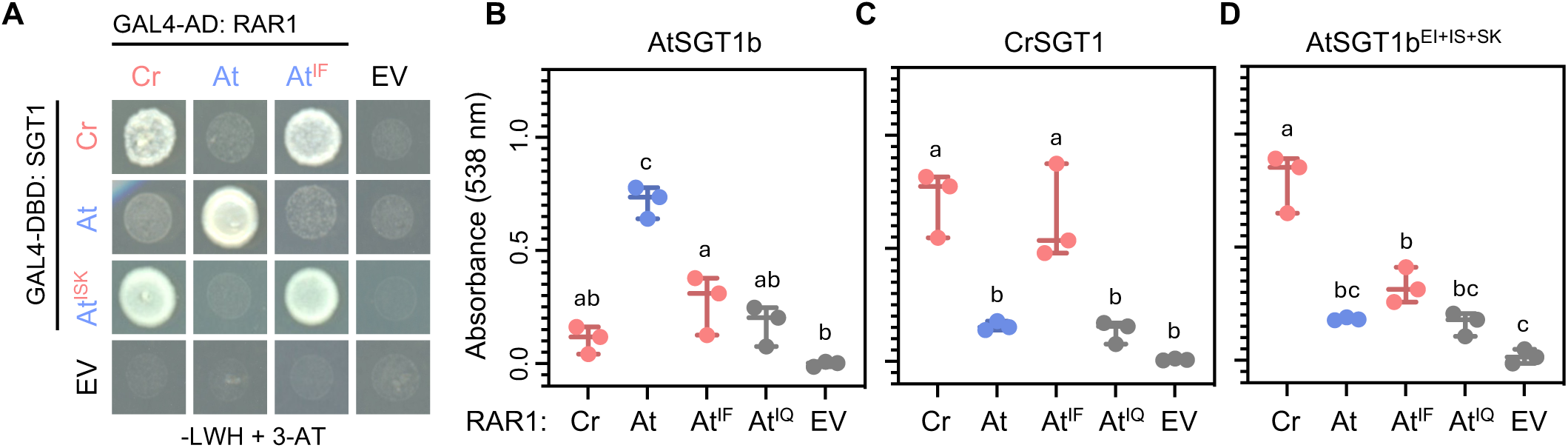
Four amino acids underpin RAR1-SGT1 interface compatibility. **(A)** Yeast-two hybrid assay comparing wildtype *C. richardii* SGT1, *A. thaliana* SGT1b, and the AtSGT1 ^EI+IS+SK^ triple variant (referred to as ‘ISK’ for simplicity) fused to the GAL4-DBD (DNA binding domain) tested against wildtype *C. richardii* RAR1, wildtype *A. thaliana* RAR1, and the AtRAR1^IF^ variant. Images were taken 3 days after plating and show yeast on media without Leu, Trp, and His (-LWH) and supplemented with 15 mM 3-AT (3-amino-1,2,4-triazole). “EV” indicates empty vector controls. This experiment was performed three times with similar results. Extended yeast two-hybrid data related to this figure are included in Data S1. **(B)** *In planta* split GAL4 RUBY interaction assay testing wildtype *A. thaliana* SGT1b against *C. richardii* RAR1, *A. thaliana* RAR1, the *A. thaliana* RAR1^IF^ and RAR1^IQ^ variants, and an empty vector (EV) control. **(C)** *In planta* split GAL4 RUBY interaction assay testing wildtype *C. richardii* SGT1 against *C. richardii* RAR1, *A. thaliana* RAR1, the *A. thaliana* RAR1^IF^ and RAR1^IQ^ variants, and an empty vector (EV) control. **(D)** *In planta* split GAL4 RUBY interaction assay testing the *A. thaliana* SGT1b^EI+IS+SK^ triple variant against *C. richardii* RAR1, *A. thaliana* RAR1, the *A. thaliana* RAR1^IF^ and RAR1^IQ^ variants, and an empty vector (EV) control. For data displayed in B-D, each datapoint represents the mean of single experimental replicate (n = 6 leaves per experiment) to a total of three biological replicates. Horizontal lines denote the mean of the data and error bars represent standard deviation. Different letters indicate statistically significant differences as determined by ANOVA (p < 0.05, Tukey’s HSD).

Next, we performed split *GAL4 RUBY* assays to quantify the RAR1-SGT1 interaction *in planta*. In this assay, proteins of interest are fused to a GAL4 DNA-binding domain (bait: GAL4^DB^) or a VP16 activation domain (prey: VP16^AD^) and transiently co-expressed in *Nicotiana benthamiana* leaves by *Agrobacterium tumefaciens*. When proteins interact, the reconstituted transcription factor activates a *RUBY* betalain biosynthesis reporter cassette leading to red pigmentation that is easily extracted and measured by spectrophotometry. In support of the utility of this approach, AtSGT1b-GAL4^DB^ and VP16^AD^-AtRAR1 produced strong accumulation of betalain pigments indicative of a strong interaction (Chen et al. 2023). Co-expressing AtSGT1b-GAL4^DB^ and VP16^AD^-RAR1 variants led to strong accumulation of pigments with AtRAR1 but not fusions with CrRAR1, AtRAR1^I153F^, or the AtRAR1^I153Q^ negative control (Fig. 4B). As anticipated, CrSGT1-GAL4^DB^ showed strong pigmentation when co-expressed with CrRAR1 and AtRAR1^I153F^ but not wildtype AtRAR1 or AtRAR1^I153Q^ (Fig. 4C). Lastly, we interrogated binding specificity using the AtSGT1b^EI+IS+SK^ variant, which showed strong pigmentation only with CrRAR1 and a weaker increase with AtRAR1^I153F^, though this was not statistically significant. Importantly, AtSGT1b^EI+IS+SK^ failed to interact with wild-type AtRAR1 or the AtRAR1^I153Q^ control (Fig. 4D). Altogether, the data support the hypothesis that four key amino acid residues dictate RAR1-SGT1 binding specificity in plants.

### Stepwise divergence in the SGT1 interface precedes diversification of RAR1 in leptosporangiate ferns

To better understand how RAR1 and SGT1 diversified within ferns, we surveyed critical RAR1 (AtRAR1:Ile193) and SGT1 (AtSGT1b:Glu191, Ile193, Ser195) interface residues in extant species representing the full diversity of eusporangiate and leptosporangiate ferns (Fig 5). This revealed that the RAR1 Ile-to-Phe substitution occurs only in Cyatheales and Polypodiales, the latter of which includes the model fern *C. richardii*. Indeed, ferns were the only lineage to encode bulky aromatic amino acids (phenylalanine or tyrosine) at this position when the survey was expanded to include other land plant lineages (Fig. S6). Across all land plants, the most prevalent amino acid at the AtRAR1:Ile153 position was leucine, which is encoded by six possible codons (TTA, TTG, and CTN). Eusporangiate ferns, a taxonomic group that diverged earlier than leptosporangiate ferns encode TTG codons that produce leucine at the Ile153 position. By contrast, later-diverging leptosporangiate ferns belong to Cyatheales and Polypodiales encode phenylalanine predominantly with the codon TTC and occasionally through the TTT codon (Fig. S6). Collectively, these data suggest that a single G-to-C nucleotide polymorphism occurred as the ancestor of Cyatheales and Polypodiales diverged from other ferns, resulting in an isoleucine-to-phenylalanine substitution that imparts RAR1-SGT1 binding specificity.

**Figure 5.**
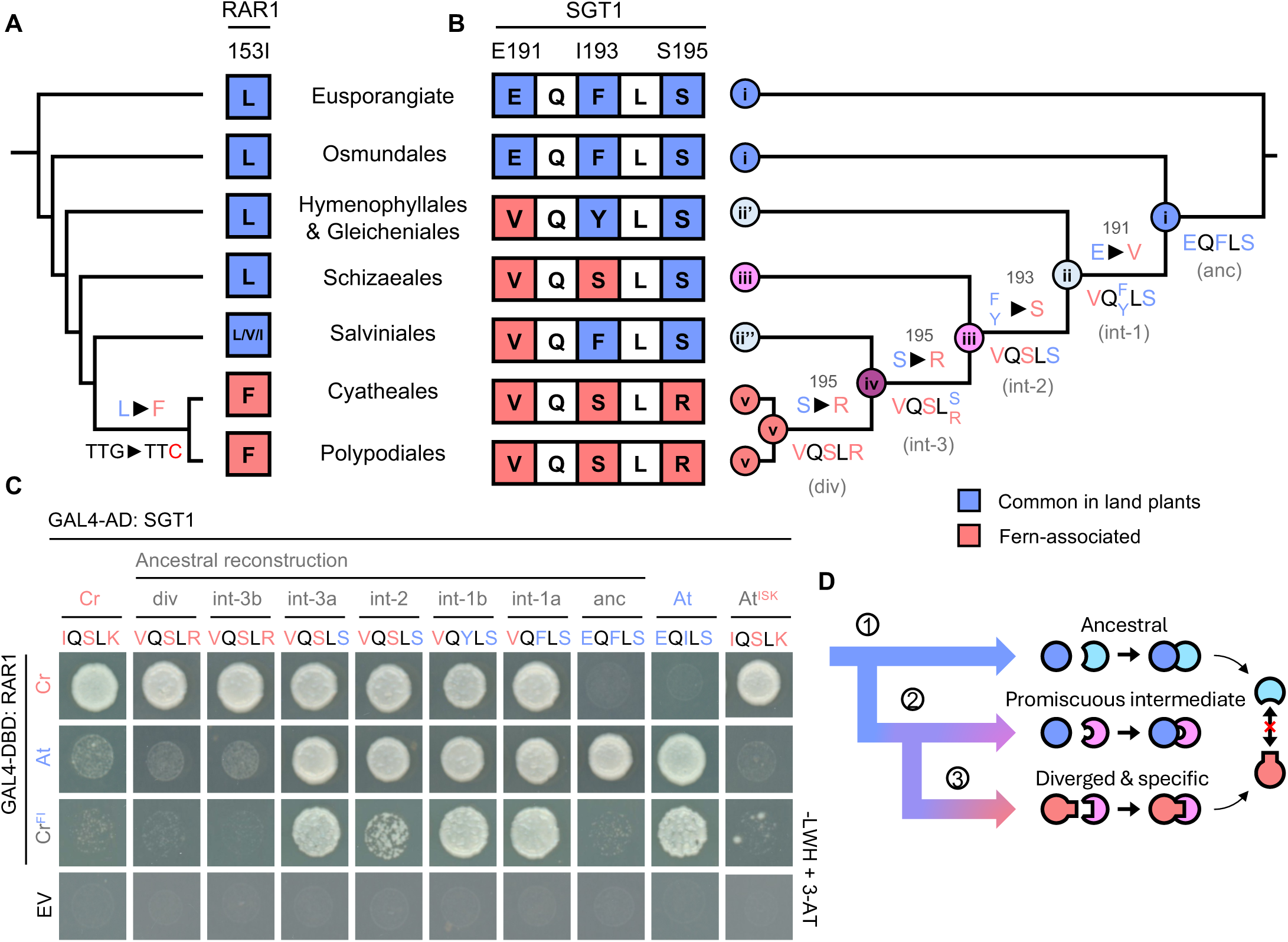
Retracing RAR1-SGT1 interface divergence in leptosporangiate ferns. **(A)** Phylogeny of representative fern lineages indicating the consensus amino acid at position Ile153 in the RAR1 CHORDII domain. Clear codon change observed across Cyatheales and Polypodiales related to other ferns (leucine to phenylalanine substitution) is denoted on the tree. Amino acids are labelled as common across diverse lineage or are specific to ferns. **(B)** Ancestral state reconstruction of fern SGT1 displaying the sequential accumulation of fern-specific amino acid substitutions in the CS domain. Ancestral states are noted on corresponding branches and were assigned categories based on their identity (i, ancestral/common state; ii-iv, intermediate states; v, divergent state). **(C)** Yeast-two hybrid assay using ancestrally reconstructed fern SGT1 states and AtRAR1, CrRAR1, and the CrRAR1^FI^ mutant. RAR1 homologs fused to the GAL4-DBD (DNA binding domain) were tested against SGT1 homologs/variants (At, AtSGT1b; Cr, CrSGT1b; At^ISK^, AtSGT1b^ISK^) and ancestral states (corresponding to B and Table S5) to Gal4-AD (activation domain). Key SGT1 interface residues are depicted. Images were taken 3 days after plating and show yeast on media without Leu, Trp, and His (-LWH) and supplemented with 15 mM 3-AT (3-amino-1,2,4-triazole). EV indicates empty vector controls. This experiment was performed three times with similar results for the Cr, At, and EV baits; and two times with similar results for the Cr^FI^ bait. Extended yeast two-hybrid data related to this figure are included in Data S1. **(D)** Simple schematic outlining the sequential states of the RAR1-SGT1 interface throughout fern evolution. (1) an ancestral and commonly observed interface state is observed in early eusporangiate ferns, which (2) diversifies through a promiscuous intermediate SGT1 variant with an expanded RAR1 binding capacity, and then finally (3) RAR1-SGT1 specificity is restored by subsequent mutations that render the interface significantly different from its original state.

Three fern-specific mutations in the SGT1^CS^ domain appeared as the Osmundales diverged from other leptosporangiate ferns. Interestingly, the E191 ‘versatility’ substitution (hydrophobic residues enabling interactions with AtRAR1) emerged first and was followed by substitutions in the remaining two positions, which are together required to restrict interface specificity by disrupting the ancestral interaction with AtRAR1 (Fig. 5B; Fig. S4). However, the order by which the I193S and S195K mutations evolved could not be determined by analysing extant species alone due to apparent reversions at the Ile193 and Ser195 positions in the Salviniales. To overcome this limitation and retrace the evolutionary trajectory of fern SGT1, we performed ancestral state reconstruction (Brayer et al. 2011; Hochberg and Thornton 2017; Laursen et al. 2021) using protein and nucleotide sequence of 60 *SGT1* homolog sequences from 56 extant fern species (Table S1) and the liverwort Mp*SGT1* as an outgroup (Fig. 5B; Fig. S7 and S8). This clearly predicted the sequential acquisition of the three fern-specific mutations in SGT1, which began with a glutamate to valine substitution at position 191 (E191) after divergence from the Osmundales (from ancestral state i to ii) as expected from patterns observed in extant ferns (Fig. 5B; Table S5). Interestingly, phenylalanine at position 193 was replaced by tyrosine, a similar bulky aromatic amino acid, in the common ancestor of Hymenophyllales and Gleicheniales (ancestral state ii’). This was followed by a phenylalanine/tyrosine to serine substitution at position 193 (ancestral state iii) occurring as Schizaeales diverged from Hymenophyllales. Lastly, the binding interface experienced a serine to arginine substitution at position 195 that began in the common ancestor of the Salviniales, Cyatheales and Polypodiales (ancestral state iv) and is fixed in the common ancestor of the Cyatheales and Polypodiales (ancestral state v). Ancestral state reconstruction also suggests that a reversion from serine to the bulky amino acid phenylalanine at position 193 likely occurred in the common ancestor of Salviniales (ancestral state ii’’). These analyses predict that the binding interface diverged first within the SGT1^CS^ domain, accumulating a sequence of modifications that enabled subsequent diversification in RAR1^CHORDII^. Given that the first SGT1 residue to diversify was the E191 residue that expands SGT1 binding capacity in yeast two-hybrid assays (E191I), our data support the idea that divergence in the RAR1-SGT1 interface was facilitated by the emergence of a ‘promiscuous’ SGT1 intermediate that facilitated subsequent diversification in RAR1.

We first explored whether shifts in RAR1-SGT1 binding specificity occur as the fern lineage diversified by performing yeast two-hybrid analyses using orthologs sourced from extant fern species. This included RAR1-SGT1 pairs from *C. richardii* (Pol; Polypodiales), *Azolla filiculoides* (Sal; Salviniales), *Hymenophyllum dentatum* (Hym; Hymenophyllales), and *Osmunda sp.* (Osm; Osmundales) that were tested alongside Arabidopsis as a control (Ang; Angiosperm) (Fig. S9A). Similar to previous results, within species pairs of RAR1-SGT1 clearly interacted and showed clear signs of yeast growth, consistent with the idea that RAR1-SGT1 interactions are conserved across diverse plants, and the *C. richardii* and *A. thaliana* RAR1-SGT1 pairs failed to interact with one another (Fig. S9B; Data S1). By comparison, yeast-two hybrid screening showed a gradual shift in the interaction compatibility between RAR1 and SGT1 homologs across ferns. An exception to this was OspRAR1 (Osm) that unexpectedly interacted with CrSGT1 (Pol) despite being more similar to the Arabidopsis-type interface identity, although OspSGT1 interacted with AtRAR1 and representatives from intermediate Hym and Osm fern lineages and not CrRAR1 or AfRAR1 (Pol or Sal) (Figure S9B; Data S1). HdSGT1 (Hym) and AfSGT1 (Sal), which represent homologs encoding the predicted promiscuous transition state, were more versatile than homologs from later-diverging leptosporangiate ferns (Pol), earlier-diverging ferns (Osm), or angiosperms (Ang). These data hint toward further diversification within RAR1 and SGT1 present across extant ferns. Given this complexity, we propose that an alternative approach utilising ancestral state reconstruction is best suited to directly assess how interface specificity emerged and evolved in ferns.

Ancestral state reconstruction suggested the stepwise divergence towards specificity in the SGT1^CS^ domain of leptosporangiate ferns that concludes with a compensatory substitution within the RAR1^CHORDII^ domain. To better support this predicted shift in specificity throughout fern evolution, we tested interaction compatibility of the major ancestral states of fern SGT1 against *Arabidopsis* and *Ceratopteris* RAR1. Our analyses included seven predicted ancestral versions of SGT1 that ranged from the predicted ancestor (“anc” - ancestral state i), two predicted versions of first intermediate (int-1a/b – ancestral state ii), second intermediate (int-2 – ancestral state iii), two predicted versions of the third intermediate (int-3a/b – ancestral state iv), and the diverged state (div – ancestral state v). We also included wildtype CrSGT1, AtSGT1b, and the triple AtSGT1b^ISK^ variant as controls. Yeast growth indicative of protein interaction was observed for CrRAR1 paired with CrSGT1 and AtSGT1b^ISK^ but not wildtype AtSGT1b, whereas AtRAR1 interacted with AtSGT1b but not CrSGT1 or AtSGT1b^ISK^ (Fig. 5C; Data S1). As anticipated, CrRAR1 failed to interact with the ancestral fern SGT1 that carries an Arabidopsis-like interface (anc) but clearly interacted with all other ancestral states of fern SGT (int-1a/b, int-2, int-3ab, and div). Each of these predicted variants notably encode the valine ‘191’ residue compared to the glutamate encoded in ancestral or *Arabidopsis* SGT1. By comparison, AtRAR1 interacted with the ancestral fern SGT1 (anc) and intermediate variants carrying two of the three fern-associated substitutions (int-1a/b, int-2, int-3a), but failed to interact with the diverged (div) state or the predicted third intermediate carrying all three substitutions (int-3b) (Fig. 5C; Data S1).

Lastly, we generated a CrRAR1 variant substituting the diversified phenylalanine to an aliphatic isoleucine (CrRAR1^FI^) and tested whether this was sufficient to alter it’s binding profile. Strikingly, CrRAR1^FI^ failed to interact with CrSGT1, the AtSGT1b^EI+IS+SK^ triple variant, or diverged ancestral states of fern SGT1 carrying all three amino acid substitutions in the RAR1 binding interface (div, int-3b) (Fig. 5C, Data S1). Instead, this variant interacted with wild-type AtSGT1b and intermediate ancestral states of fern SGT1 carrying one or two substitutions in the three key RAR1-interacting residues (int-1a/b, int-2, int-3a). Unexpectedly, CrRAR1^FI^ also failed to interact with the ancestral (anc) fern SGT1 that carries a binding interface comparable to AtSGT1b and interacted with AtRAR1 (Fig. 5C; Data S1). Considered alongside our interaction data for extant fern RAR1-SGT1 pairs, this suggests that additional factors likely influence interface specificity. In any event, our data suggests that the RAR1^I153F^ substitution is likely a compensatory mutation that adapts to structural changes in the RAR1-SGT1 interface imposed by divergence in SGT1. Based on our functional interaction assays and evolutionary analyses, we conclude that diversification within the RAR1-SGT1 interface occurred first in SGT1, where an initial substitution from the ancestral state common amongst land plants towards a more promiscuous state expanded SGT1 interaction capcities. This enabled subsequent diversification that ultimately facilitated further divergence in SGT1 and RAR1 that locked in binding specificity (Fig. 5D).

## DISCUSSION

In this study, we discovered that the RAR1-SGT1 binding interface is widely conserved in land plants but underwent lineage-specific divergence through a promiscuous SGT1 intermediate in leptosporangiate ferns. The incompatibility between CrRAR1 and SGT1 homologs of other plants demonstrates that the RAR1-SGT1 binding interface significantly diverged in *C*. *richardii*. By leveraging RAR1 and SGT1 sequence diversity in extant land plants, we identified the minimal set of amino acid substitutions required to shift RAR1-SGT1 interaction compatibilities between *Arabidopsis* and *Ceratopteris* orthologs. Re-tracing the evolutionary history of these residues, we discovered a stepwise divergence that occurred first in SGT1 through the emergence of a promiscuous intermediate state and culminated in compensatory substitutions in RAR1 to maintain interface integrity.

RAR1 and SGT1 are broadly conserved across land plants and in some streptophyte algae. Strikingly, the level of conservation is such that mismatched RAR1 and SGT1 orthologs from algae and land plants retain the capacity to interact despite separating from one another >500 million years ago (Morris et al. 2018; Puttick et al. 2018). This pattern is similar to other conserved macromolecular complexes in plants, including those associated with post transcriptional silencing (Jia et al. 2021), transcription co-factor assembly (Ji et al. 2023), and auxin-related protein-DNA interactions (Rienstra et al. 2025). In plants, RAR1 and SGT1 complex with HSP90 to sustain intracellular NLR immune receptors that detect and respond to pathogen activity (Shirasu 2009; Jones et al. 2024). Consequently, *rar1* and *sgt1* mutants are more susceptible to pathogen attack (Azevedo et al. 2002; Tör et al. 2002; Tornero et al. 2002; Azevedo et al. 2006). Whether this conserved immune complex conditions NLRs from algae-to-angiosperms across the plant kingdom remains to be determined, however diverse plants and streptophytes like *C. braunii* encode NLRs with predicted roles in immunity (Gao et al. 2018; Chia et al. 2024; Feng et al. 2024). Future functional interrogations of NLRs and disease resistance across non-flowering plants and algae may therefore be facilitated by interrogating mutants of the RAR1-SGT1-HSP90 complex.

RAR1-SGT1 interaction divergence parallels protein evolution across diverse domains of life. The type II bacterial toxin-antitoxin system composed of ParE (toxin) and ParD (antitoxin) is underpinned by highly specific interactions and has similarly shown that interface diversification is preceded by the emergence of more promiscuous intermediates (Aakre et al. 2015; Ding et al. 2022). This was also described for insect Dscam1 (Down syndrome adhesion molecule 1) adhesion proteins, where gene expansion resulted in promiscuous heterophilic interactions between paralogs that progress towards highly specific homophilic interactions (Wiseglass and Rubinstein 2024). Interaction specificity switching in the related ParB (Partitioning protein B) and Noc (Nucleoid occlusion factor) DNA-binding proteins identified intermediate variants capable of dual binding, however these variants showed reduced binding affinities compared to original ParB or Noc proteins (Jalal et al. 2020). Such mutational trajectories reflect the critical nature of these molecular interactions, where non-functional ancestral isoforms (i.e. complete loss of binding) are selected against due to their impact on organismal fitness. Our work on RAR1-SGT1 echoes this, and highlights the importance of the these co-chaperones in maintaining plant immunity.

Why has the RAR1-SGT1 interface diversified in leptosporangiate ferns? There are two possible scenarios to explain lineage-specific divergence that may not be mutually exclusive. First, direct selection pressure may have been applied by fern-infecting pathogens. In angiosperms, several pathogen-secreted effector proteins are reported to target SGT1 and disrupt RAR1-SGT1 interactions, resulting in immune suppression (Yu et al. 2020; Nakano et al. 2021; Wang et al. 2022). Targeting of SGT1 by pathogens has also been demonstrated in humans, where the *Salmonella enterica* effector protein SspH2 interferes with a human SGT1 homolog (Bhavsar et al. 2013). It is therefore conceivable that a host-pathogen arms race occurred between the ancestral fern complex and pathogen effector proteins targeting RAR1 and/or SGT1, although this would also be expected in other plant lineages impacted by pathogens. Newly established fern-pathogen interaction systems (Grenz et al. 2024; Castel et al. 2025) could facilitate future comparative studies on immunity and pathogen effector-mediated suppression in diverse plants.

Alternatively, RAR1-SGT1 divergence may have initiated through random structural transitions in a “hole-and-knob” manner. Here, a neutral promiscuity mutation is randomly introduced into the ancestral fern SGT1, which maintains its existing interaction with RAR1 but carries the potential to interact with a wider spectrum of potential RAR1 variants. Our work identifies the initial SGT1 E-to-V susbstitution (ancestral state ii) as such a mutation, and the fact that this intermediate promiscuity mutation is still detected in extant fern lineages (from Hymenophyllales to Polypodiales) demonstrates its long-term evolutionary stability. This substitution to a small aliphatic amino acid may open up the binding pocket to enable a greater range of interactions, thereby creating a structural alteration (a “hole”) that could facilitate a complementary mutation (a “knob”) in RAR1. This mechanism of an initial random variation followed by structural lock-in mirrors the macroevolutionary trajectories documented in previous ancestral reconstruction studies of other complex molecular machines, such as the V-ATPase proton pump and vertebrate hemoglobin (Finnigan et al. 2012; Pillai et al. 2020; Chisholm et al. 2024).

In summary, although RAR1–SGT1 binding has remained conserved over ∼500 myrs, our work revealed an unexpected lineage specific pattern: RAR1–SGT1 coevolution in ferns followed an evolutionary trajectory enabled by promiscuous intermediates in SGT1, which buffered interaction compatibility until a compensatory mutation in RAR1 restored binding specificity. These findings not only provide a molecular explanation for divergence in fern RAR1–SGT1 interactions but also reinforce a general evolutionary principle: promiscuous protein-protein interactions often act as an intermediate that permits interface divergence without disrupting essential cellular processes that depend on protein complex formation.

## MATERIALS AND METHODS

### Plant and microbial growth

*Nicotiana benthamiana* were grown in soil with a constant temperature of 22 °C and a long-day photoperiod (16 h light; 160 to 200 μE, fluorescent—Sylvania F58W/GRO) under controlled growth conditions. *Agrobacterium tumefaciens* GV3101 (pMP90) was grown in LB media supplemented with appropriate antibiotics at 28 °C. *Saccharomyces cerevisiae* MaV203 (Invitrogen) was grown in 2x YPAD or selective Synthetic Complete media as appropriate for transformation selection or yeast two-hybrid assays as described below.

### Identification of RAR1 and SGT1 homologs

RAR1 and SGT1 sequences used in this study were primarily obtained from SymDB (https://www.polebio.lrsv.ups-tlse.fr/symdb/). Transcriptomes or genomes of each species were searched using the hidden Markov model-based software HMMER v3.4 (http://hmmer.org/). RAR1 and SGT1 homologs of species not included in SymDB were identified using BLASTp directly in corresponding online databases or by local BLASTp using NCBI BLAST+ (Camacho et al. 2009) version 2.13.0 with e-value threshold < 10^-50^ (Table S1 and S2). Positive hits from this search were examined by reciprocal BLAST using the well annotated plant genomes of *Arabidopsis thaliana* (flowering plant, Araport11 at Phytozome (https://phytozome-next.jgi.doe.gov/)) and *Marchantia polymorpha* (bryophyte, https://marchantia.info/). For BLASTp analyses, the following five sequences were used as queries: MpRAR1 (Mp4g22990.1), AtRAR1 (AT5G51700.1), MpSGT1 (Mp5g04260.1), AtSGT1a (AT4G23570.1), and AtSGT1b (AT4G11260.1). Integrity of RAR1 and SGT1 sequences was examined by presence of key domains described by previous studies(Shirasu et al. 1999; Azevedo et al. 2002; Zhang et al. 2010) after multiple sequence alignment by MUSCLE algorithm (Edgar 2004) version 3.8.1551 using default setting of SnapGene (https://www.snapgene.com/).

### Identification of *CrRAR1* and *CrSGT1* from *C*. *richardii* cDNA library

We validated predicted *CrRAR1* and *CrSGT1* gene models using reverse-transcription PCR. In brief, total RNA was extracted from flash-frozen *C*. *richardii* gametophore tissue by the PureLink Plant RNA Reagent (Invitrogen, 12322012). DNA was then removed by TURBO DNA-free kit (Invitrogen, AM1907). cDNA was synthesized from 2 μg of total RNA using SuperScript II Reverse Transcriptase (Invitrogen, 18064014). All assays were performed following the manufacturer’s protocols.

To detect Cr*SGT1* paralogs, Cr*SGT1* annotated from a *de novo* transcriptome (Geng et al. 2021) the Cr*SGT1N* (Ceric.25g062300.1), and Cr*SGT1C* (Ceric.1Z100300.1) annotated from the *C. richardii* genome database(Marchant et al. 2022) were amplified together with Cr*RAR1* (Ceric.17G016600.1) and a fragment of Cr*ACT1* (Ceric.08g028400.1) (Fig. S10). The amplified Cr*RAR1* and Cr*SGT1* products were purified using QIAquick Gel Extraction Kit (QIAGEN, 28704). The RAR1 and SGT1 fragments were then cloned into pDONR221 using Gateway BP Clonase II Enzyme mix (Invitrogen, 11789020) and subsequently validated by diagnostic PCR and Sanger sequencing. All primers are listed in Table S3.

### Phylogenetic analyses

RAR1 and SGT1 sequences were aligned using MAAFT (Katoh and Standley 2013) v7.505 using the options “--maxiterate 1000 --localpair”. Maximum likelihood phylogenetic analyses were then performed using IQ-TREE2 (Nguyen et al. 2015; Minh et al. 2020) multicore version 2.3.2. ModelFinder (Kalyaanamoorthy et al. 2017) was performed to identify the best-fit model according to the Bayesian Information Criterion, which selected “JTTDCMut+I+G4” for RAR1 and “Q.plant+I+G4” for SGT1. Each maximum likelihood phylogeny was built using the abovementioned best-fit model with 1000 ultrafast bootstrap replicates (Hoang et al. 2018). The resulting trees were visualized using TVBOT (https://www.chiplot.online/tvbot.html) (Xie et al. 2023).

### Yeast two-hybrid cloning and interaction assays

Coding sequences of *RAR1* and *SGT1* homologs from *Chara braunii* and *Azolla filiculoides*, Cb*RAR1* (CBR_g32001), Cb*SGT1* (CBR_g30800), Af*RAR1* (Azfi_s0014.g013658), and Af*SGT1* (Azfi_s0197.g057330) were synthesized based on their genome annotations. Coding sequences of *RAR1* and *SGT1* homologs of *Hymenophyllum dentatum* and *Osmunda* sp., Hd*RAR1*, Hd*SGT1*, Osp*RAR1*, and Osp*SGT1*, were identified from transcriptome data (Tables S1 and S2) and codon-optimized for yeast and synthesized (Table S4). All gene synthesis was performed by TWIST Bioscience (https://www.twistbioscience.com/) and were prepared as Gateway entry clones. MpRAR1, MpSGT1, and AtRAR1 were cloned into the pDONR221 DONR vector using primers listed in Table S3 by BP clonase II reactions (Inivtrogen). All resulting ENTRY clones, synthesized by TWIST or cloned in-house, were subjected to LR reactions into the pDEST22 and/or pDEST32 yeast two-hybrid vectors (Invitrogen). Two *A*. *thaliana* SGT1 prey plasmids, pDEST-AD-AtSGT1a (PDEST-AD008B04) and pDEST-AD-AtSGT1b (PDEST-AD013B12) were obtained from the Arabidopsis Biological Resource Center (ABRC; https://abrc.osu.edu/).

At*RAR1* and At*SGT1b* variants carrying single amino acid polymorphisms were synthesized following the same procedures described above. BsaI domestication of At*SGT1b* and point mutations were designed according to the codon usage of *A*. *thaliana*. At*SGT1b* variants carrying multiple amino acid polymorphisms (At*SGT1b*^EI+IS^, At*SGT1b*^EI+SK^, and At*SGT1b*^EI+IS+SK^) and CrRAR1 variant carrying single amino acid polymorphism (CrRAR1^FI^) were generated by site-directed mutagenesis. The resulting variants, At*SGT1b*^EI+IS^, At*SGT1b*^EI+SK^, At*SGT1b*^EI+IS+SK^, and Cr^FI^, were subcloned into pDONR221 and verified by Sanger sequencing before further cloning into pDEST22/pDEST32 and subsequent use in yeast-two hybrid assays. All primers used for cloning are provided in Table S3.

Expression clones in Gateway-compatible pDEST32 (GAL4 DNA Binding Domain, GAL4^DB^) and pDEST22 (GAL4 DNA Activation Domain, GAL4^AD^) were generated using the Gateway recombination system (Invitrogen). Vectors were transformed into the budding yeast (*Saccharomyces cerevisiae*) MaV203 yeast two-hybrid strain was using the Frozen EZ-Yeast Transformation II Kit (Zymogen). For yeast growth, transformations, and interaction assays, Synthetic Complete (SC) medium was prepared from yeast nitrogen base without amino acids (Formedium, CYN0405), agar (Formedium, AGA03), glucose (Fisher, G/0500/53), and SC amino acid drop-out mix (-Leu -Trp, Formedium DSCK172; -Leu -Trp -His, Formedium DSCK242). For stringent His auxotrophic selection, 15 mM of 3-amino-1,2,4-triazole (Sigma-Aldrich, A8056) was supplemented to SC -Leu -Trp -His plates. A slightly higher concetration of 50 mM 3-AT was used for Gal4^DB^:HdSGT1 constructs to avoid mild autoactivation (Data S1). For URA3 counter-selection, SC -Leu -Trp medium was supplemented with 1 mg/ml 5-fluoroorotic acid (Fisher, R0811) with 1 mg/ml final concentration. Growth of yeasts on selective and counter-selective plates was performed at 30 °C and were assessed at 3 days after inoculation. All raw and extended yeast two-hybrid analyses are displaye in Data S1.

### Protein modelling and visualization

Protein interaction models were performed using AlphaFold3 (Jumper et al. 2021; Abramson et al. 2024) of the *C. richardii* SGT1-CS (amino acids 172-263) and RAR1-CHORDII (amino acids 147-214) domains (pTM = 0.8; ipTM = 0.8). This model was contrasted against an existing crystal structure of the Arabidopsis RAR1-SGT1 interface (PDB: 2XCM) (Zhang et al. 2010). Protein structures (known and predicted) were visualized in Chimera X (Pettersen et al. 2021) to highlight areas of interest.

### Split GAL4 RUBY assay

RAR1 and SGT1 orthologs and their associated variants were amplified by PCR using primers containing BsaI cloning sites for subsequent cloning into level 2 Golden Gate-compatible acceptors for the split GAL4 Ruby assay (Zenodo, doi: 10.5281/zenodo.15873182): pICSL22026 and pICSL22027 (TSL SynBio). Purified *SGT1* fragments or a GUS control (Addgene #50327), were assembled with one of two promoters, pICH85281 (mas promoter + TMVΩ, Addgene #50272) or pICH87633 (Nos promoter+ TMVΩ, Addgene #50271), alongside the GAL4 DNA-binding domain (pICSL50065; GAL4DBD, TSL SynBio), and the 35S terminator (pICH41414, AddGene #50337) into the acceptor plasmid pICSL22026 (TSL SynBio). Purified *RAR1* fragments or the GUS control (Addgene #50327), were assembled with one of two promoters, pICSL13004 (mas promoter + TMVΩ 5’ UTR, TSL Synbio) or pICSL13010 (Nos promoter + TMVΩ, TSL Synbio), alongside the VP16 activation domain (pICSL30055, TSL SynBio), and pICH41414 into the acceptor plasmid pICSL22027. All final constructs were verified by Sanger sequencing.

Vectors were electroporated into *A*grobacterium *tumefaciens* GV3101. For infiltration, equal amounts of three independent *A*. *tumefaciens* liquid overnight cultures carrying the RAR1 construct, the SGT1 construct, and a strain carrying the P19 silencing suppressor were mixed at 1:1:1 ratio to the desired final OD_600_. Final OD_600_ was set to 0.1 for mas promoter contructs and 0.5 for Nos promoter constructs. All assays were initially performed using mas constrcuts, however weak pigmentation patterns for the CrSGT1-CrRAR1 control interaction prompted us to perform subsequent assays for CrSGT1 using Nos promoter combinations. Before pressure infiltrating expanded leaves of 4-week-old *N. benthamiana* leaves using 1 mL needleless syringes, the mixed *A*. *tumefaciens* cultures were incubated in induction buffer (10 mM MES pH 5.6, 10 mM MgCl_2_, and 150 µM acetosyringone) for 1 h in the dark.

Quantification of betalain accumulation was performed as previously described (Chen et al. 2023) with minor modifications. *Agrobacterium*-infiltrated *N*. *benthamiana* leaves were harvested and cleared of chlorophyll overnight in 100% ethanol. For each *RAR1*–*SGT1* combination, 8 mm diameter leaf discs were collected from the centre of each infiltration site across six independent leaves and pooled. The pooled discs were extracted overnight in 0.6 mL of water. Water-soluble betalain absorbance of the clean supernatant was measured at 538 nm wavelength using a spectrophotometer (CLARIOstar Plus, BMG Labtech) with three technical replicates.

### Ancestral state reconstruction

Protein and nucleotide sequences of 60 fern SGT1 homologs from 56 fern species and the MpSGT1 (outgroup) were used in ancestral state reconstruction (Supplementary Fig. 1). The protein seqences were aligned using MAFFT (Katoh and Standley 2013) and a phylogenetic tree was created using IQ-TREE (Nguyen et al. 2015; Minh et al. 2020) multicore version 2.3.2. The “MG+F3X4+R5” model was chosen by ModelFinder (Kalyaanamoorthy et al. 2017) as the best-fit model and a maximum likelihood was built using 1000 ultrafast bootstrap replicates (Hoang et al. 2018). To generate a codon-based nucleotide sequence alignment input, nucleotide sequences for each gene were threaded onto the amino acid alignment using phykit (Steenwyk et al. 2021) v2.0.1, and the resulting codon-based alignment was verified by back-translation of each sequence. Ancestral state reconstruction was conducted using the CODON model of the “ancseq” pipeline (https://github.com/YuSugihara/ancseq) (Sugihara et al. 2025). Briefly, this pipeline integrates IQ-TREE’s empirical Bayes method with a Jukes-Cantor type binary model to reconstruct both ancestral sequences and indels. The results were visualized in iTOL (Letunic and Bork 2024) v6 with annotated node numbers. Statistical values associated with ancestral state reconstruction at critical interaction residues are provided in Table S5.

### Statistical Analyses

Details regarding statistical analyses are indicated in appropriate figure legends. The statistical tests employed, sample size (i.e., n = number of samples per condition) and dispersion/precision measurements are provided (error bars, p value cutoffs, etc.). Statistical analysis of betalaine accumulation was performed using R software with the two-way ANOVA ‘aov’ function on raw data that passed Shapiro-Wilk normality testing. Differences between interaction groups were determined by Tukey’s HSD post-hoc testing (p < 0.05) and significance grouping between treatments was defined using the multcompView package (Graves et al. 2024).

## ACKNOWLEDGEMENTS

We thank TSL SynBio for access to Golden Gate cloning vectors and we thank all current and past members of the Carella group at the John Innes Centre for technical assistance and insightful discussions on RAR1-SGT1 coevolution. We thank Sophien Kamoun (TSL) for constructive discussions on protein complex evolution and for commenting on an earlier draft of the manuscript.

## AUTHOR CONTRIBUTIONS

H.J. and P.C. designed the research; H.J., Y.S., and P.C. performed the research; H.J., Y.S., M.W.W, and P.C. analyzed the data; H.J. and P.C. prepared figures and final datasets; H.J. and P.C. wrote the manuscript with contributions from all authors.

## SUPPLEMENTARY DATA

Figure S1. Annotated RAR1 phylogeny Figure S2. Annotated SGT1 phylogeny

Figure S3. HSP90-interacting residues in RAR1 and SGT1 are conserved across land plants

Figure S4. Extended analysis of SGT1 evolutionary diversity at the RAR1-interacting residues

Figuer S5. A fern-associated valine amino acid variant broadens binding capacity in AtSGT1b

Figure S6. Extended analysis of RAR1 evolutionary diversity at the Ile 153 position

Figure S7. Annotated monilophyte SGT1 phylogeny

Figure S8. Summary of ancestral state reconstruction for key SGT1 interface residues in the fern lineage

Figure S9. Assessing diverse RAR1-SGT1 interactions in extant ferns

Figure S10. Cloning of *CrRAR1* and *CrSGT1* from *Ceratopteris richardii*

Table S1. List of species used in this study

Table S2. List of SGT1 paralogs in land plants

Table S3. Primers used in this study

Table S4. Synthesized DNA constructs used in this study

Table S5. SGT1 ancestral state reconstruction data

Data S1. Raw data and extended yeast two-hybrid analyses

## DATA AVAILABILITY

Relevant gene identifiers are provided in the manuscript and in the supporting information online.

---

**Figure S1.**
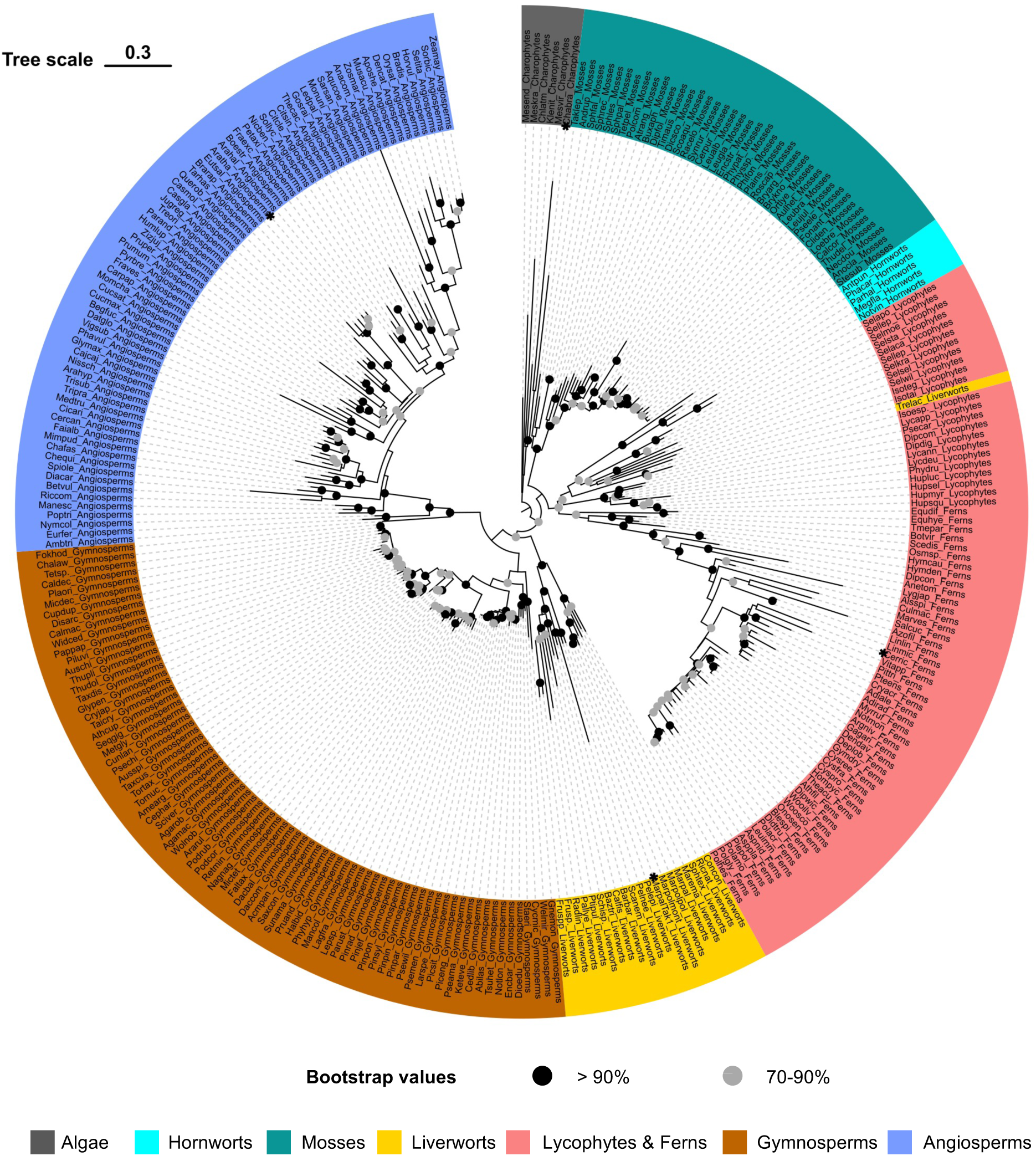
Annotated RAR1 phylogeny. Maximum likelihood phylogeny of RAR1 proteins with 1000 ultrafast bootstrap replicates across diverse land plants and algae. Model = “JTTDCMut+I+G4”. Tree scale = substitutions per site. Related to Figure 1B. Orthologs are colored based on their clade across streptophyte lineages: charophytes(gray), hornworts (cyan), liverworts (yellow), mosses (teal), lycophytes and ferns (salmon pink), gymnosperms (brown), and angiosperms (blue). This phylogenetic tree is rooted using streptophyte algal sequences as the outgroup. Bootstrap values are indicated as circles: black (> 90%) and light gray (70-90%). Asterisks indicate RAR1 homologs tested in Figure 1d.

**Figure S2.**
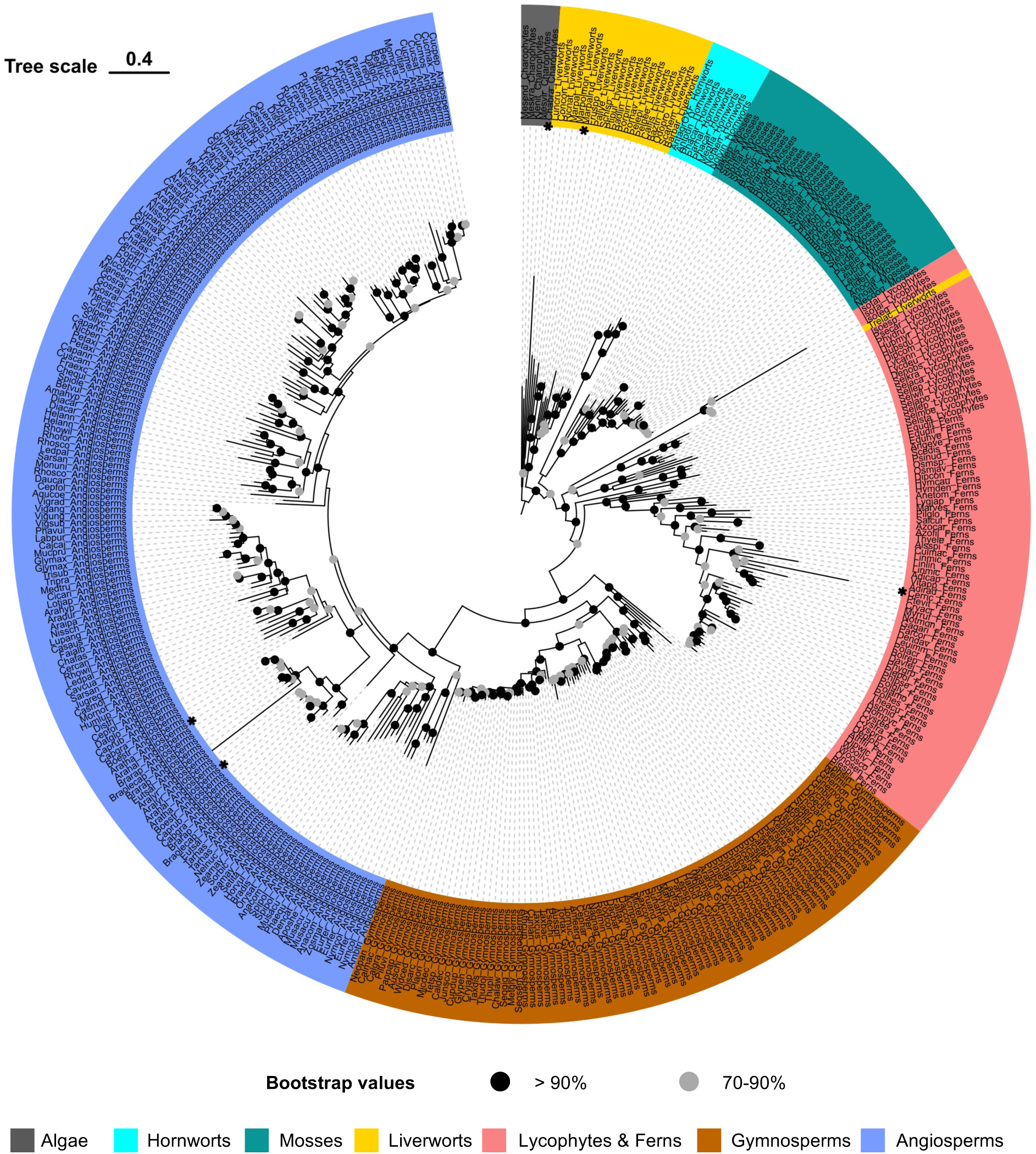
Annotated SGT1 phylogeny. Maximum likelihood phylogeny of SGT1 proteins with 1000 ultrafast bootstrap replicates across diverse land plants and algae. Model = “Q.plant+I+G4” . Tree scale = substitutions per site. Related to Figure 1C. Orthologs are colored based on their clade across streptophyte lineages: charophytes(gray), hornworts (cyan), liverworts (yellow), mosses (teal), lycophytes and ferns (salmon pink), gymnosperms (brown), and angiosperms (blue). This phylogenetic tree is rooted using streptophyte algal sequences as the outgroup. Bootstrap values are indicated as circles: black (> 90%) and light gray (70-90%). Asterisks indicate SGT1 homologs tested in Figure 1d.

**Figure S3.**
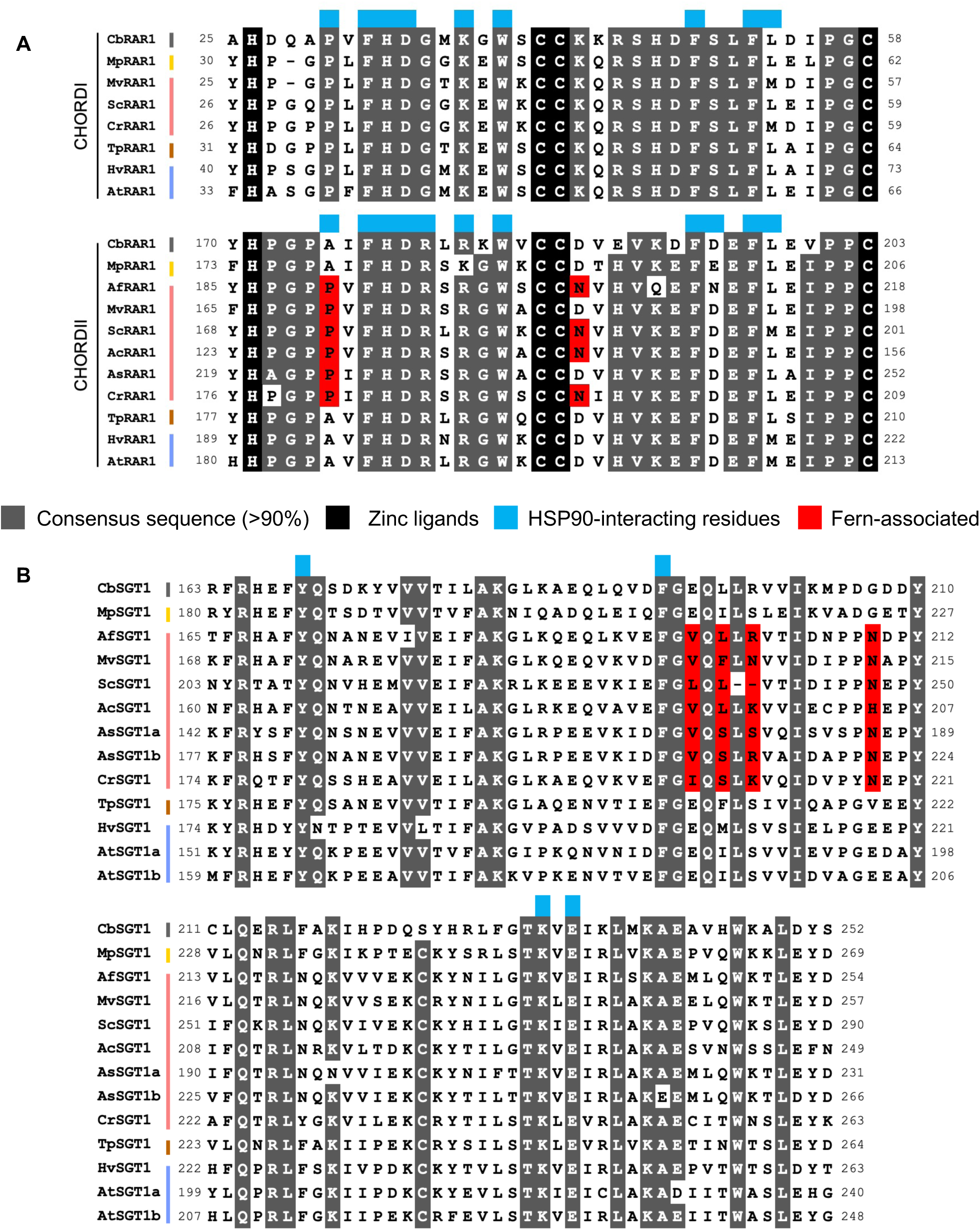
HSP90-interacting residues in RAR1 and SGT1 are conserved across land plants. The known HSP90-interacting residues of the CHORD domains of AtRAR1 (A) and the CS domain of SGT1 (B) are labelled with blue boxes on the top of the multiple sequence alignments. The species set from Fig. 2 was used. Orthologs are colored with the same scheme as Fig. 2: charophyte (grey), bryophyte (yellow), monilophyte (salmon pink), gymnosperm (brown), and angiosperm (blue).

**Figure S4.**
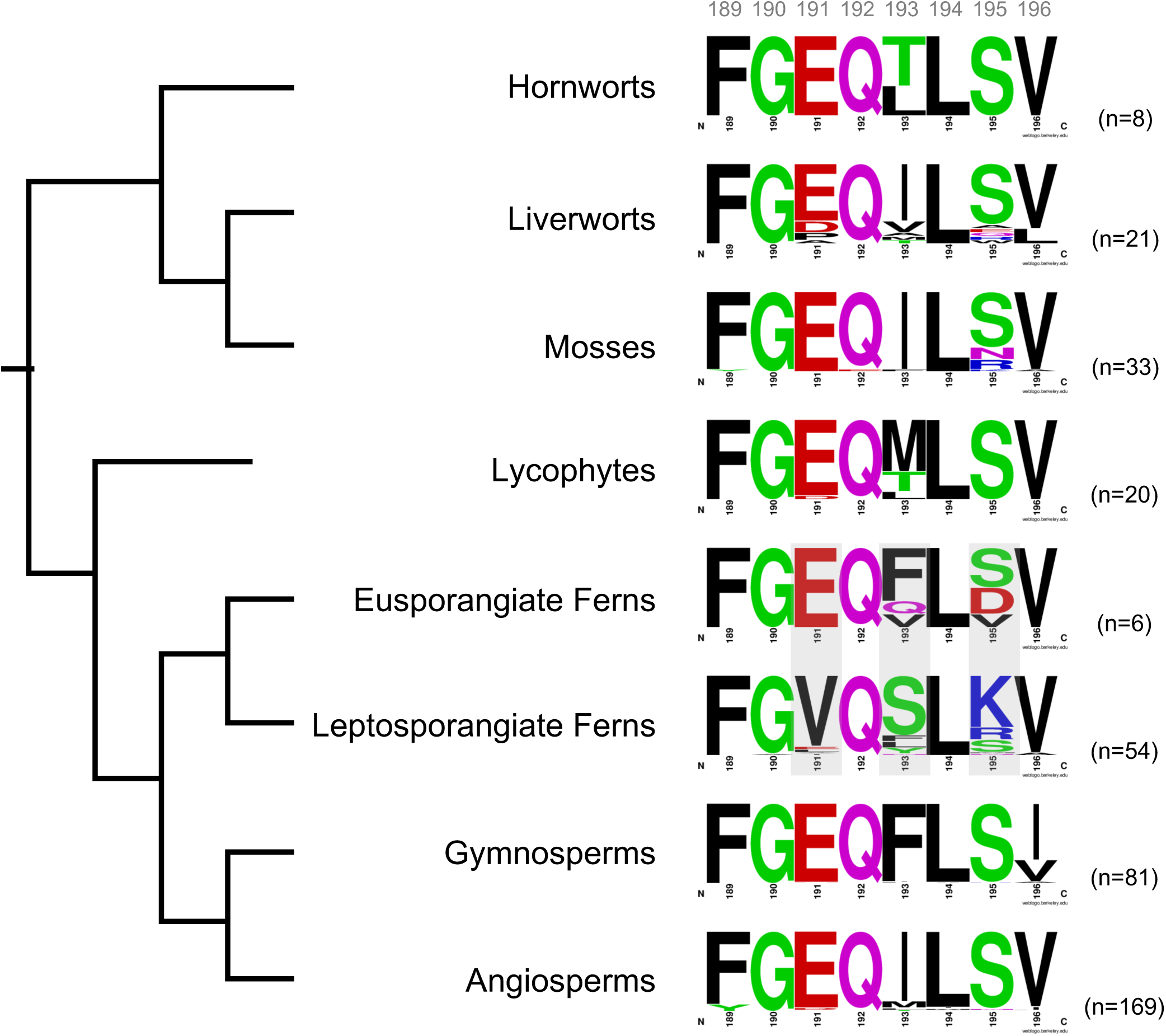
Extended analysis of SGT1 evolutionary diversity at the RAR1-interacting residues. A schematic phylogeny illustrates evolutionary relationships between lineages. Consensus amino acid identity between the position 189 and 196 is shown for each clade. Amino acid frequency plot at this polymorphic region is illustrated by sequence logo (WebLogo: https://weblogo.berkeley.edu/logo.cgi). The total number of SGT1 homologs used for comparative analyses (n) are indicated next to frequency plots. Gray shade indicates three critical positions identified in Figure 3: Glu191, Ile193, and Ser195.

**Figure S5.**
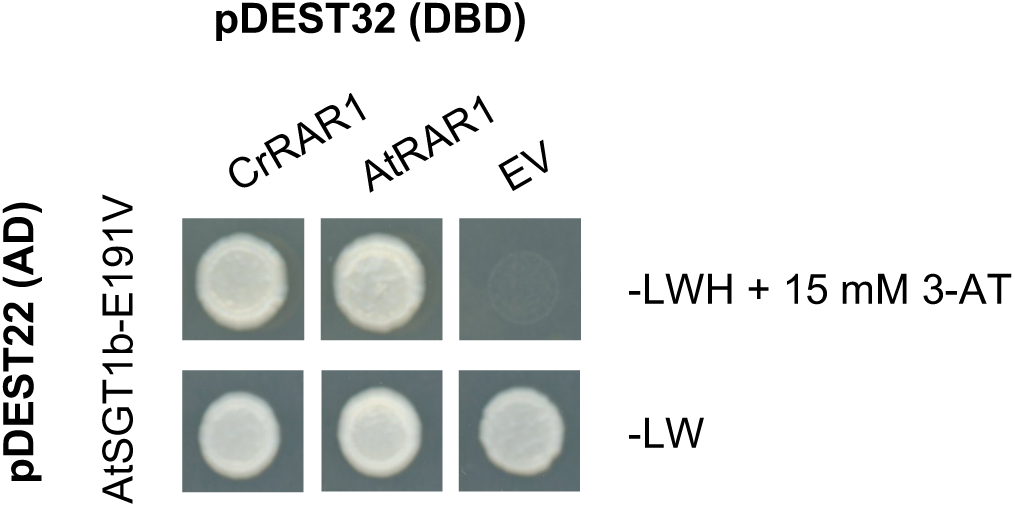
A fern-associated valine amino acid variant broadens binding capacity in AtSGT1b. Yeast two-hybrid of an E191V (glutamate-to-vaine) amino acid substitution introduced into AtSGT1b (prey vector pDEST22; AD, activation domain) enables interaction with both CrRAR1 and AtRAR1 (bait vector; DBD, DNA binding domain). Images show yeast plated onto SC (synthetic complete) media without Leu, Trp (-LW), or onto SC media without Leu, Trp, and His (-LWH) and supplemented with 15 mM 3-AT (3-aminotriazole). This experiment was performed at least three times with similar results. Full interaction assays results are shown in Data S1.

**Figure S6.**
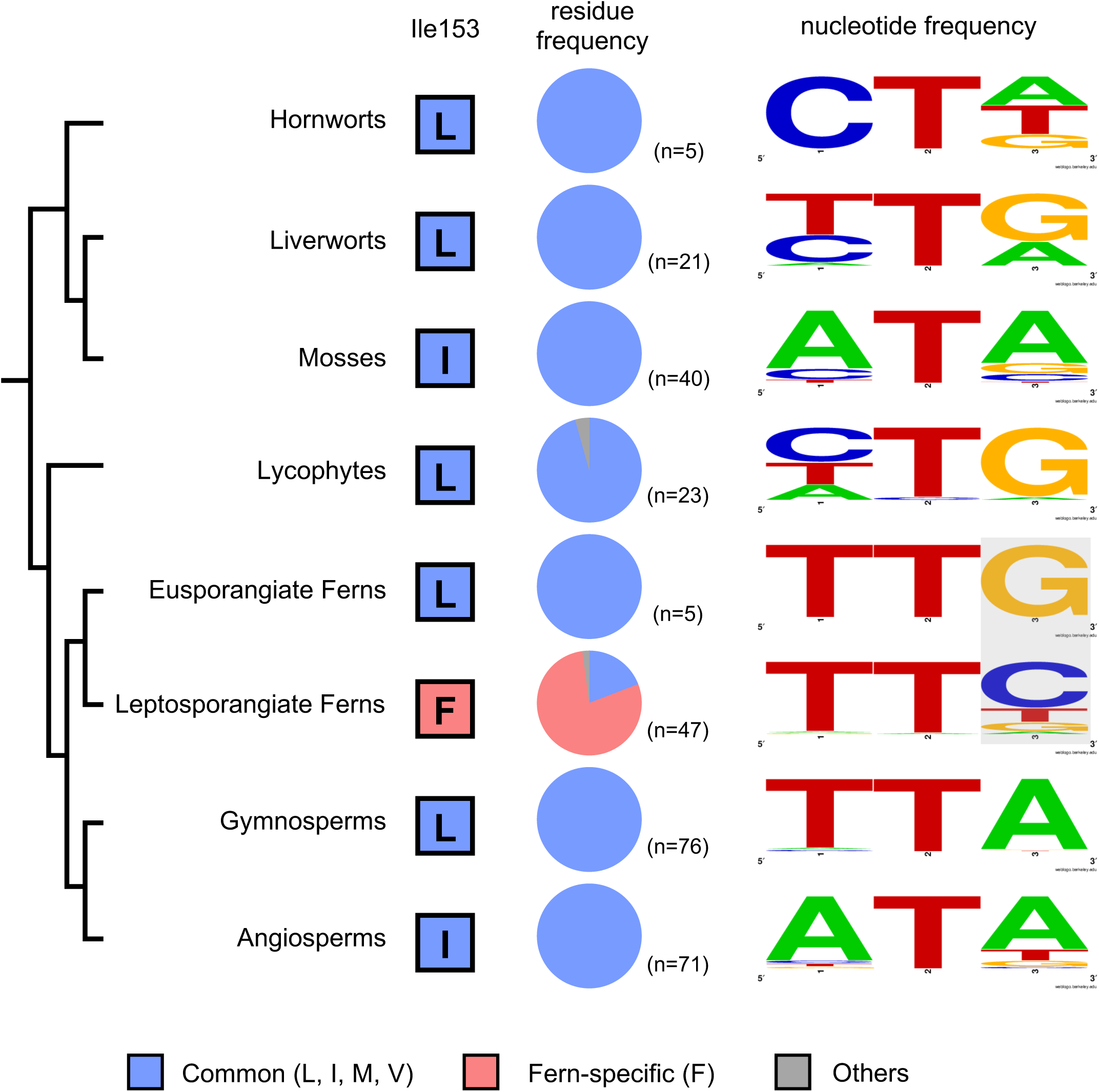
Extended analysis of RAR1 evolutionary diversity at the Ile 153 position. A schematic phylogeny illustrates evolutionary relationships between lineages. Consensus amino acid identity at the I153 position is shown for each clade, with frequences of common (hydrophobic amino acids such as Leu, Ile, Met, and Val) compared to fern-specific phenylalanine (Phe) or other unique amino acids (others) not commonly observed (less than 1% frequency) in land plants. The total number of RAR1 homologs used for comparative analyses (n) are indicated next to frequency pie charts. Nucleotide (codon) frequency plot at this position is also illustrated by sequence logo (WebLogo: https://weblogo.berkeley.edu/logo.cgi). Gray shade indicates G-to-C and G-to-T single nucleotide polymorphism resulting Ile-to-Phe mutation in ferns. Related to Figure 5A.

**Figure S7.**
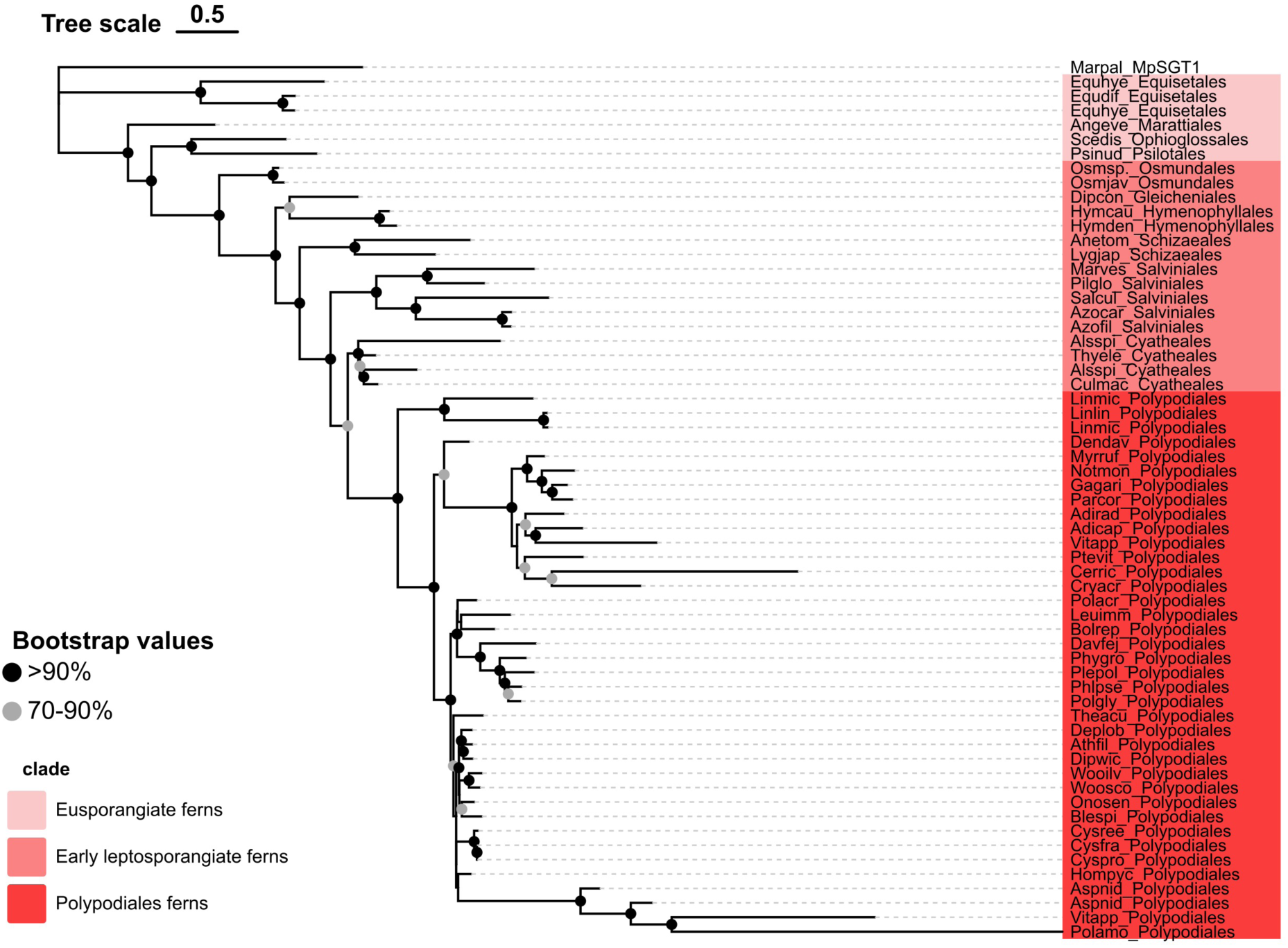
Annotated Monilophyte SGT1 phylogeny. Diverse SGT1 orthologs sourced from ferns and the outrgroup MpSGT1 control. Maximum likelihood phylogeny (model = “MG+F3X4+R5”) with bootstrap values indicated. Tree scale = substitutions per site. Related to Figure 5B.

**Figure S8.**
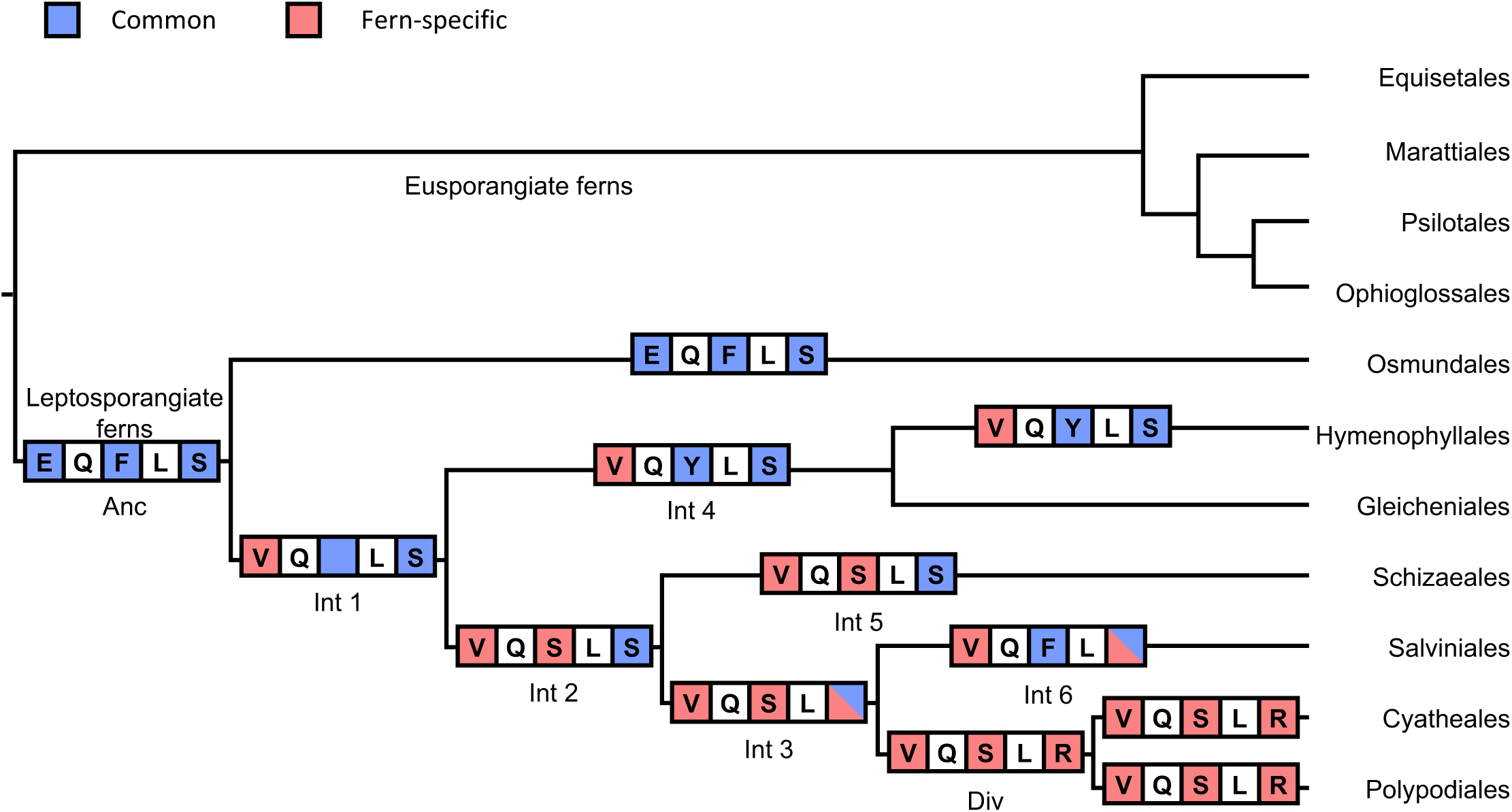
Summary of ancestral state reconstruction for key SGT1 interface residues in the fern lineage. Squares without amino acid abbreviation indicates ambiguous residues, where each possible amino acid has more than 40% of probability. Predicted codons and amino acid identities for each ancestral states (Ancestral, Anc; Intermediate, Int 1-6; Diverged, Div) are described in Supplementary Table 5. Related to Figure 5B.

**Figure S9.**
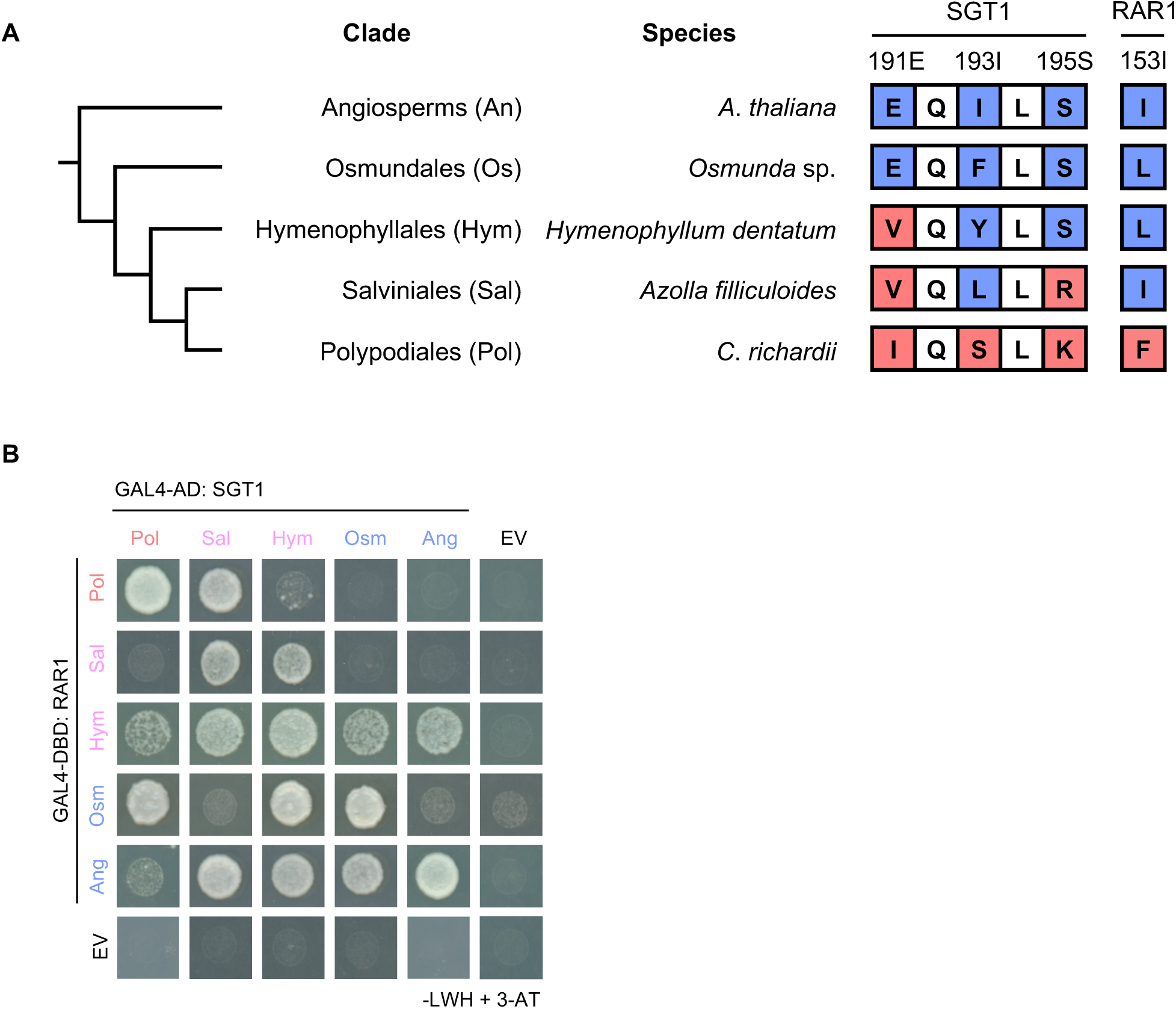
Assessing diverse RAR1-SGT1 interactions in extant ferns. **(A)** Diverse fern species and the identity of key amino acid residues within their RAR1-SGT1 binding interfaces. Related to Figure 5C. **(B)** Yeast two-hybrid testing of diverse fern RAR1-SGT1 orthologs. -LWH indicates SC media without Leu, Trp, or His and supplemented with 15 mM 3-aminotriazole (AT). Images show yeast growth 3 days post plating. “EV” indicates empty vector controls. This experiment was performed at least twice with similar results.

**Figure S10.**
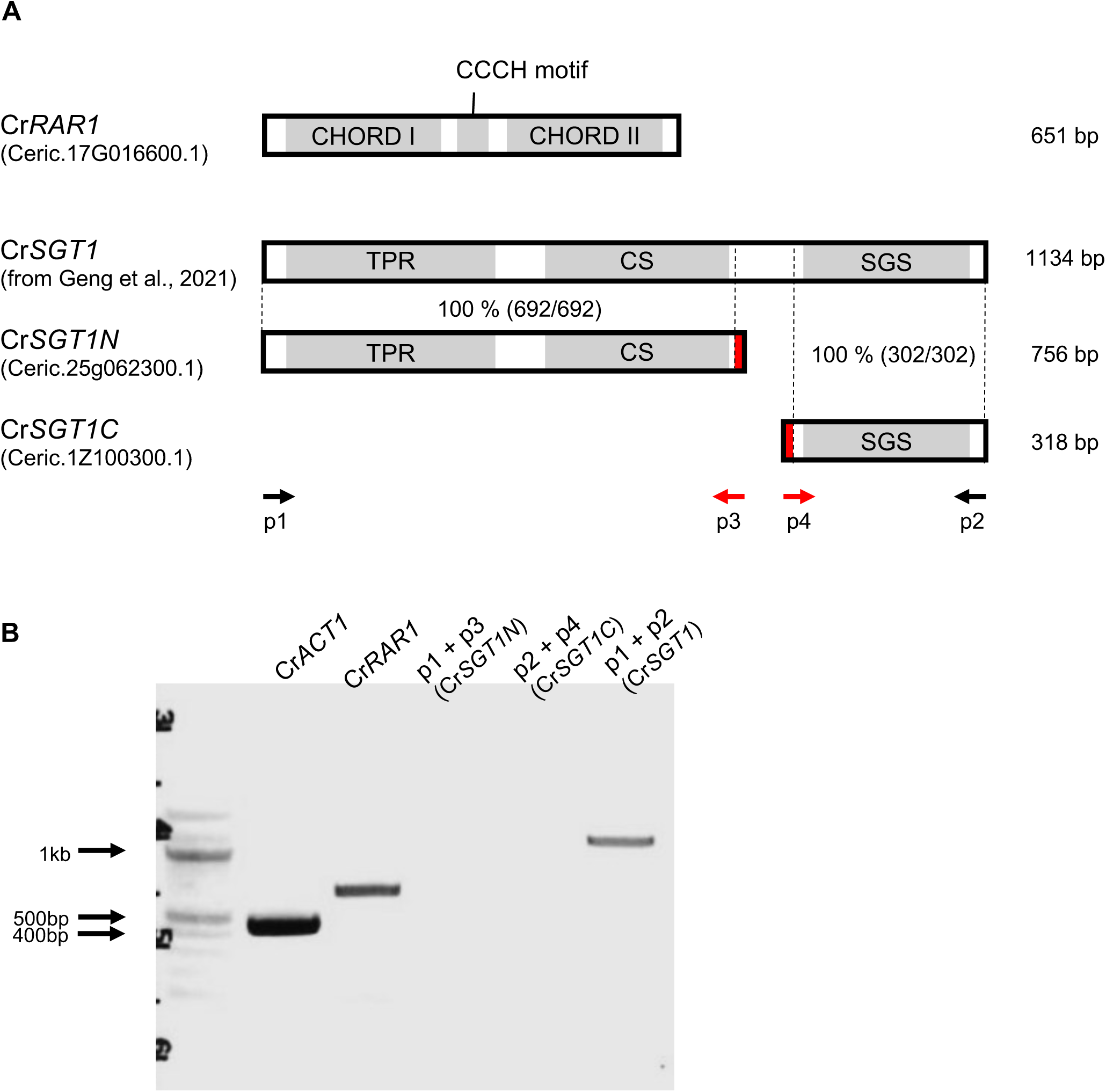
Cloning of Cr*RAR1* and Cr*SGT1* from *C*. *richardii* cDNA. **(A)** Schematic illustrating *Cr*RAR1 and *Cr*SGT1 CDS gene models from public datasets. Red patches indicate unique sequence on Cr*SGT1N* and Cr*SGT1C*. Primers (p1 - p4) that were designed on common (p1 and p2) or unique sequences (p3 and p4) of these models. **(B)** RT-PCR analyses of CrACTIN1, CrRAR1, and common and specific CrSGT1 primers using *C*. *richardii* gametophyte cDNA.

