## Supplementary material for "Stepwise and lineage-specific divergence of a major immune co-chaperone complex in leptosporangiate ferns": Data S1

#### Data S1 – Raw data and extended yeast two-hybrid analyses

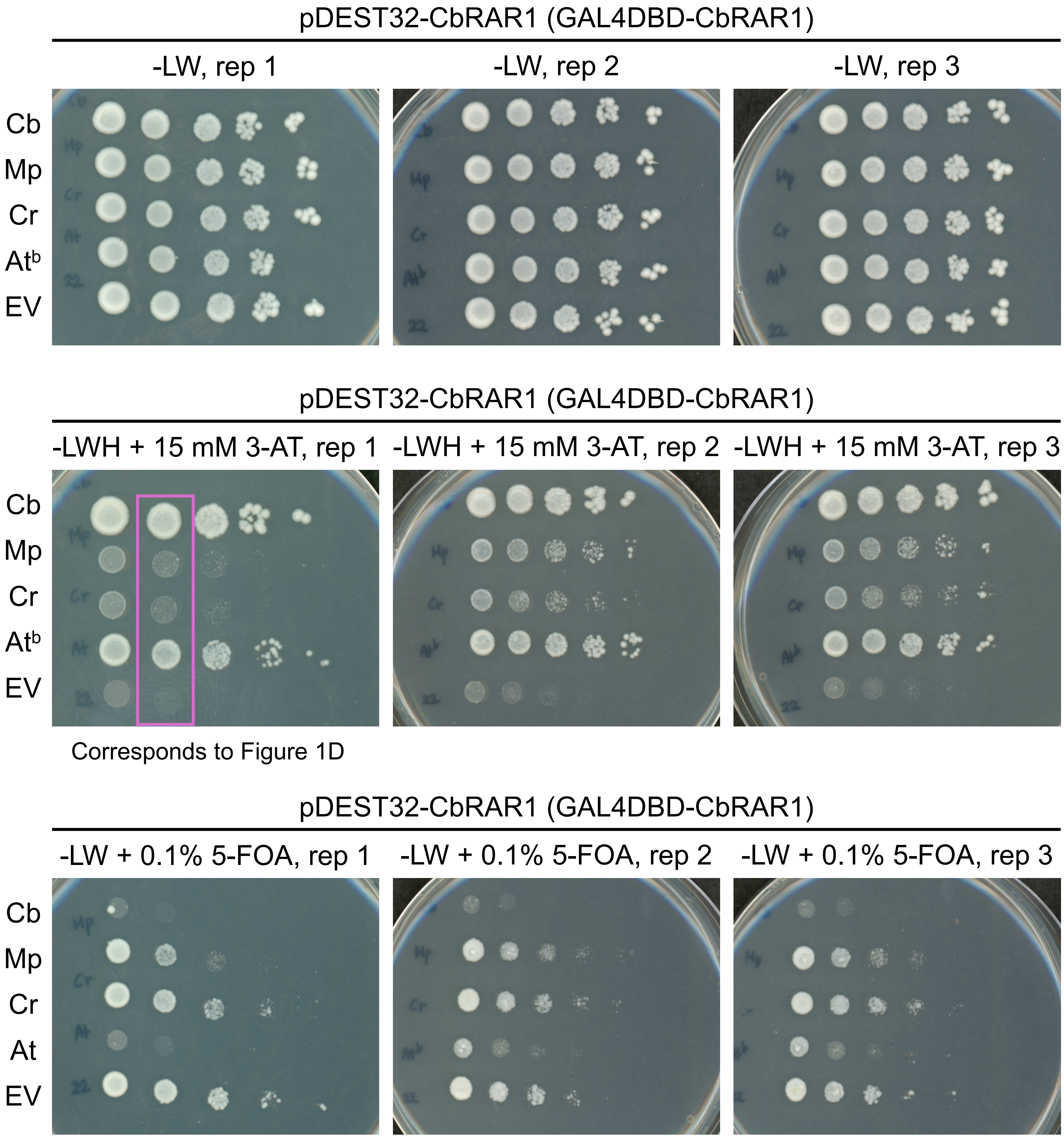


Cb; CbSGT1-pDEST22
Mp; MpSGT1-pDEST22
Cr; CrSGT1-pDEST22
At; pDEST-AD-SGT1b
EV; empty pDEST22

(serial dilution was started to from $OD_{600}$ = 1.0 and finished at $OD_{600}$ = 1.0 × $10^{-4}$)

(every plate was scanned three days after inoculation)

### Data S1 – Raw data and extended yeast two-hybrid analyses

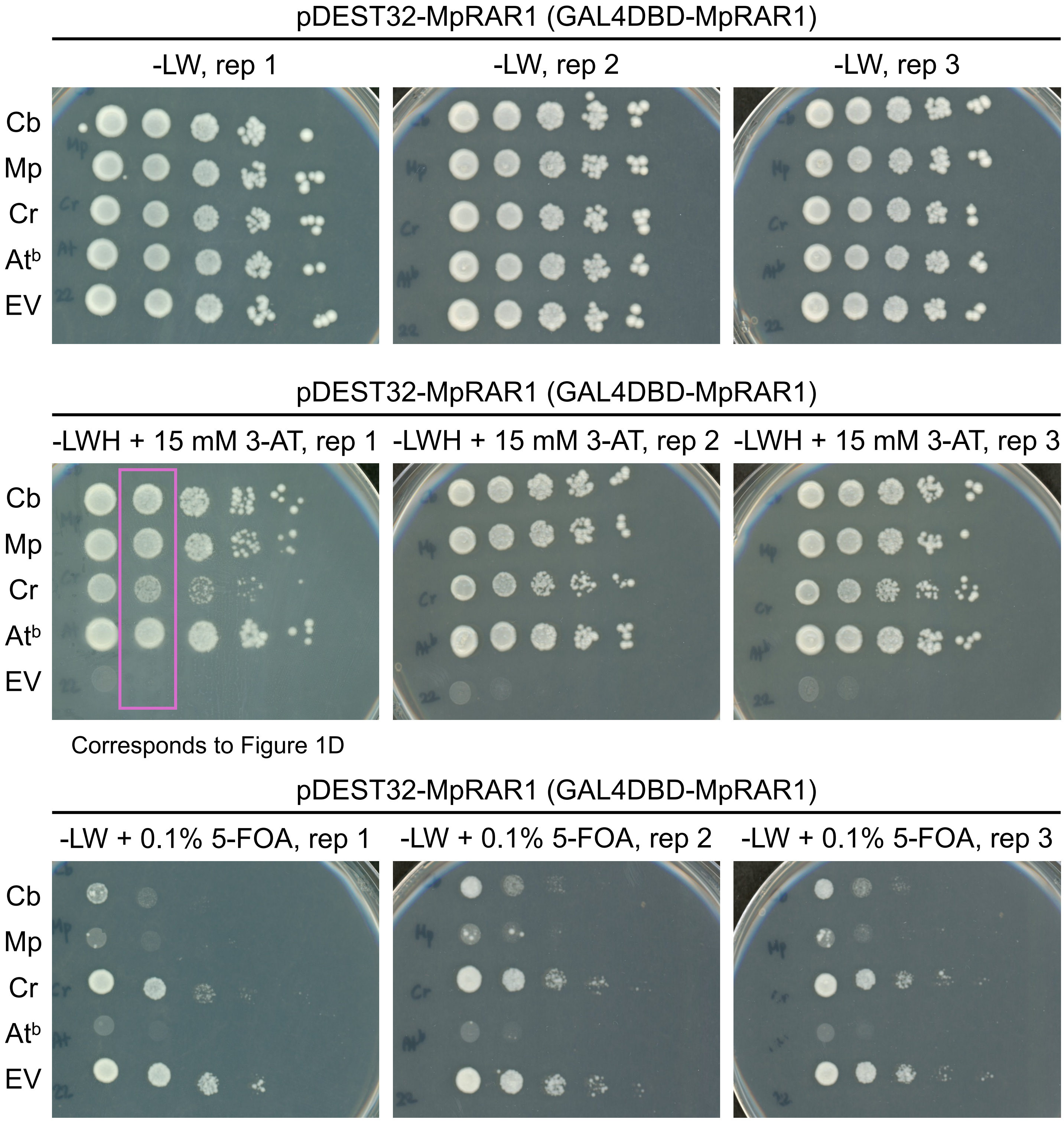


Cb; CbSGT1-pDEST22
Mp; MpSGT1-pDEST22
Cr; CrSGT1-pDEST22
At$^{b}$; pDEST-AD-SGT1b
EV; empty pDEST22

(serial dilution was started to from $OD_{600}$ = 1.0 and finished at $OD_{600}$ = 1.0 × $10^{-4}$)
(every plate was scanned three days after inoculation)

#### Data S1 – Raw data and extended yeast two-hybrid analyses

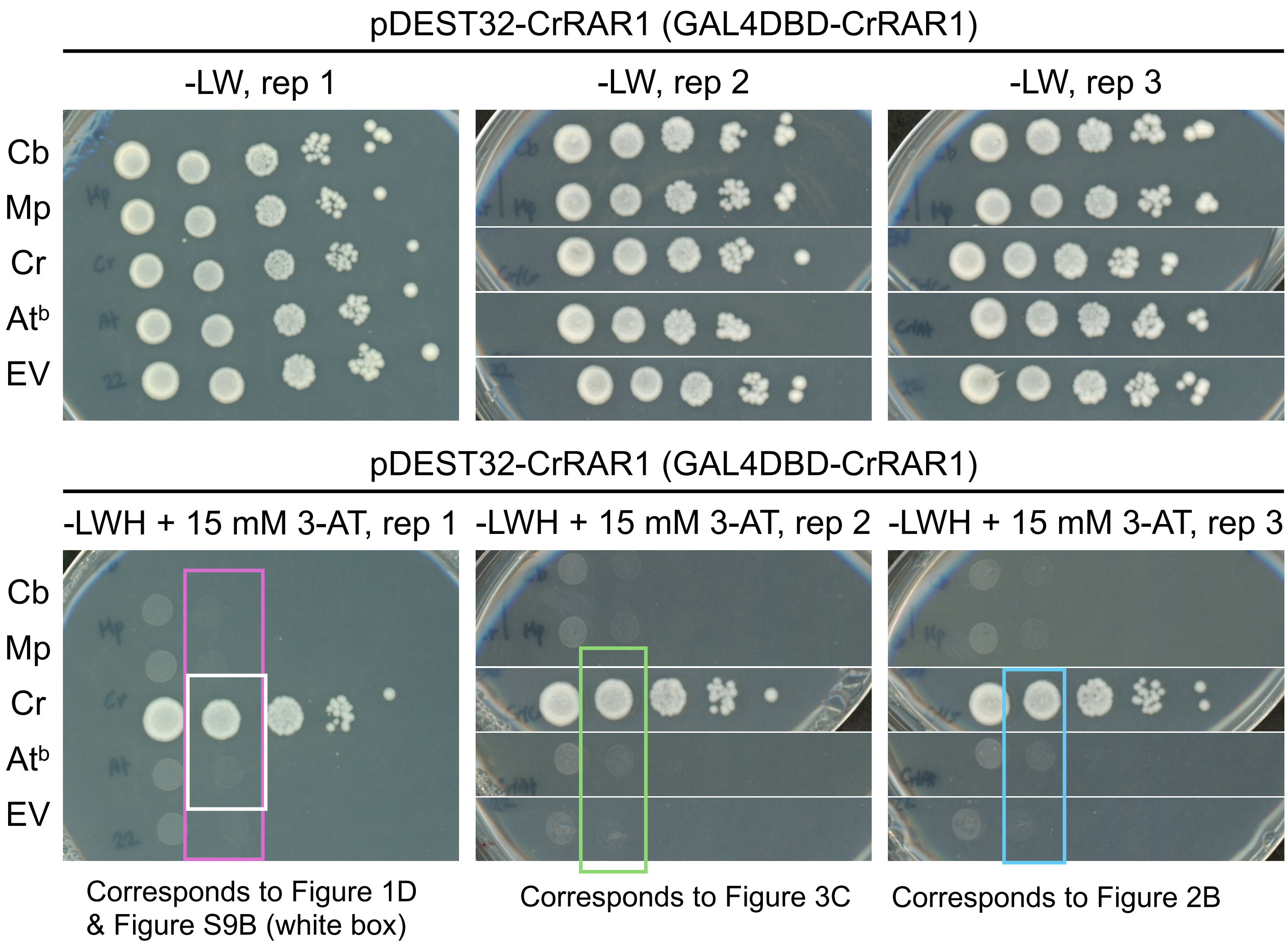


Cb; CbSGT1-pDEST22
Mp; MpSGT1-pDEST22
Cr; CrSGT1-pDEST22
At$^{b}$; pDEST-AD-SGT1b
EV; empty pDEST22

(serial dilution was started to from $OD_{600}$ = 1.0 and finished at $OD_{600}$ = 1.0 × $10^{-4}$)

(every plate was scanned three days after inoculation)

#### Data S1 – Raw data and extended yeast two-hybrid analyses

pDEST32-AtRAR1 (GAL4DBD-AtRAR1)

| -LW, rep 1 | -LW, rep 2 | -LW, rep 3 |
|---|---|---|

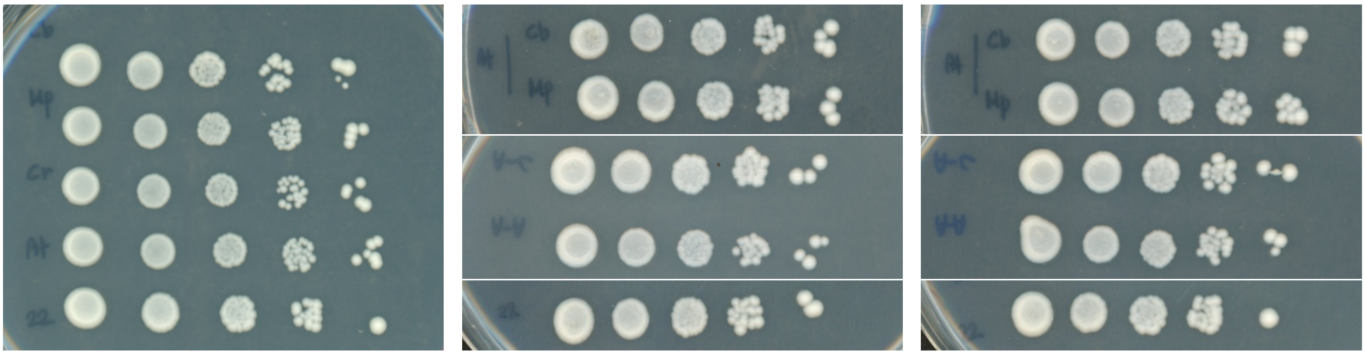


pDEST32-AtRAR1 (GAL4DBD-AtRAR1)

| -LWH + 15 mM 3-AT, rep 1 | -LWH + 15 mM 3-AT, rep 2 | -LWH + 15 mM 3-AT, rep 3 |
|---|---|---|

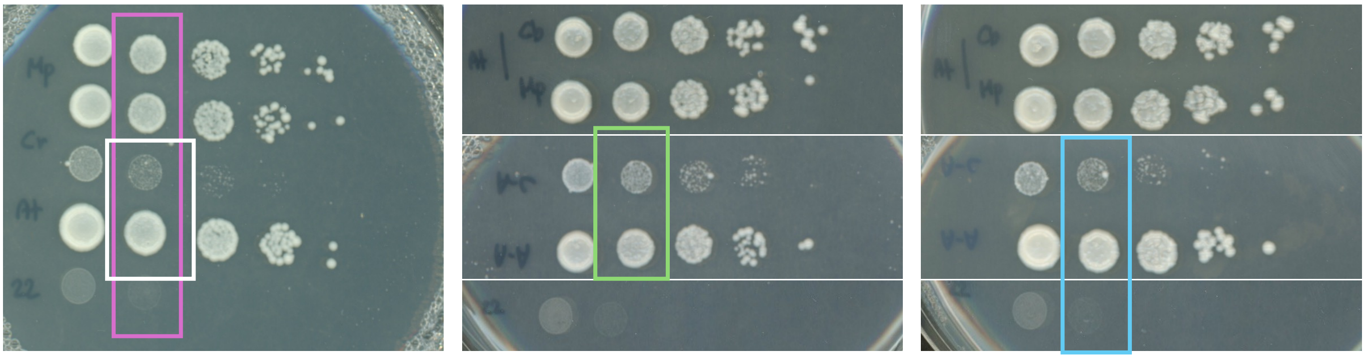


| Corresponds to Figure 1D & Figure S9B (white box) | Corresponds to Figure 3C | Corresponds to Figure 2B |
|---|---|---|

pDEST32-AtRAR1 (GAL4DBD-AtRAR1)

| -LW + 0.1% 5-FOA, rep 1 | -LW + 0.1% 5-FOA, rep 2 | -LW + 0.1% 5-FOA, rep 3 |
|---|---|---|

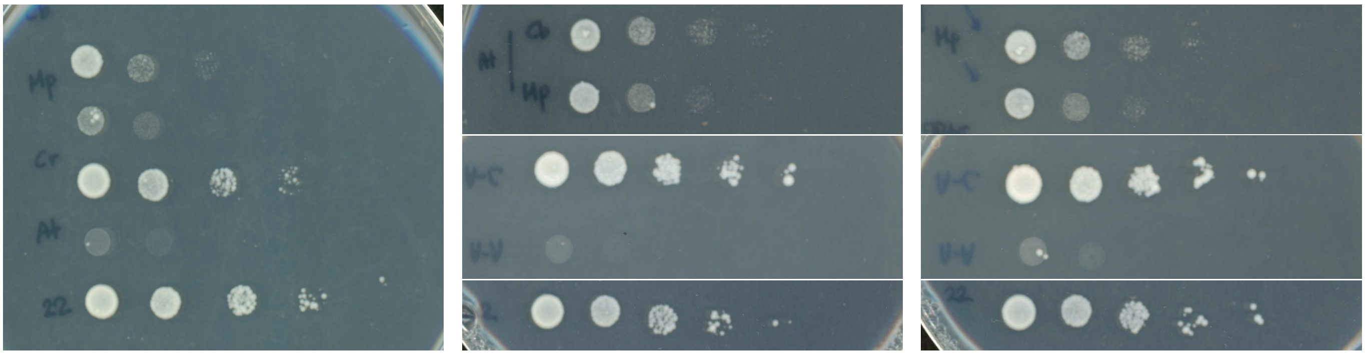


Cb; CbSGT1-pDEST22
Mp; MpSGT1-pDEST22
Cr; CrSGT1-pDEST22
At$^{b}$; pDEST-AD-SGT1b
EV; empty pDEST22

(serial dilution was started to from $OD_{600}$ = 1.0 and finished at $OD_{600}$ = 1.0 × $10^{-4}$)

(every plate was scanned three days after inoculation)

### Data S1 – Raw data and extended yeast two-hybrid analyses

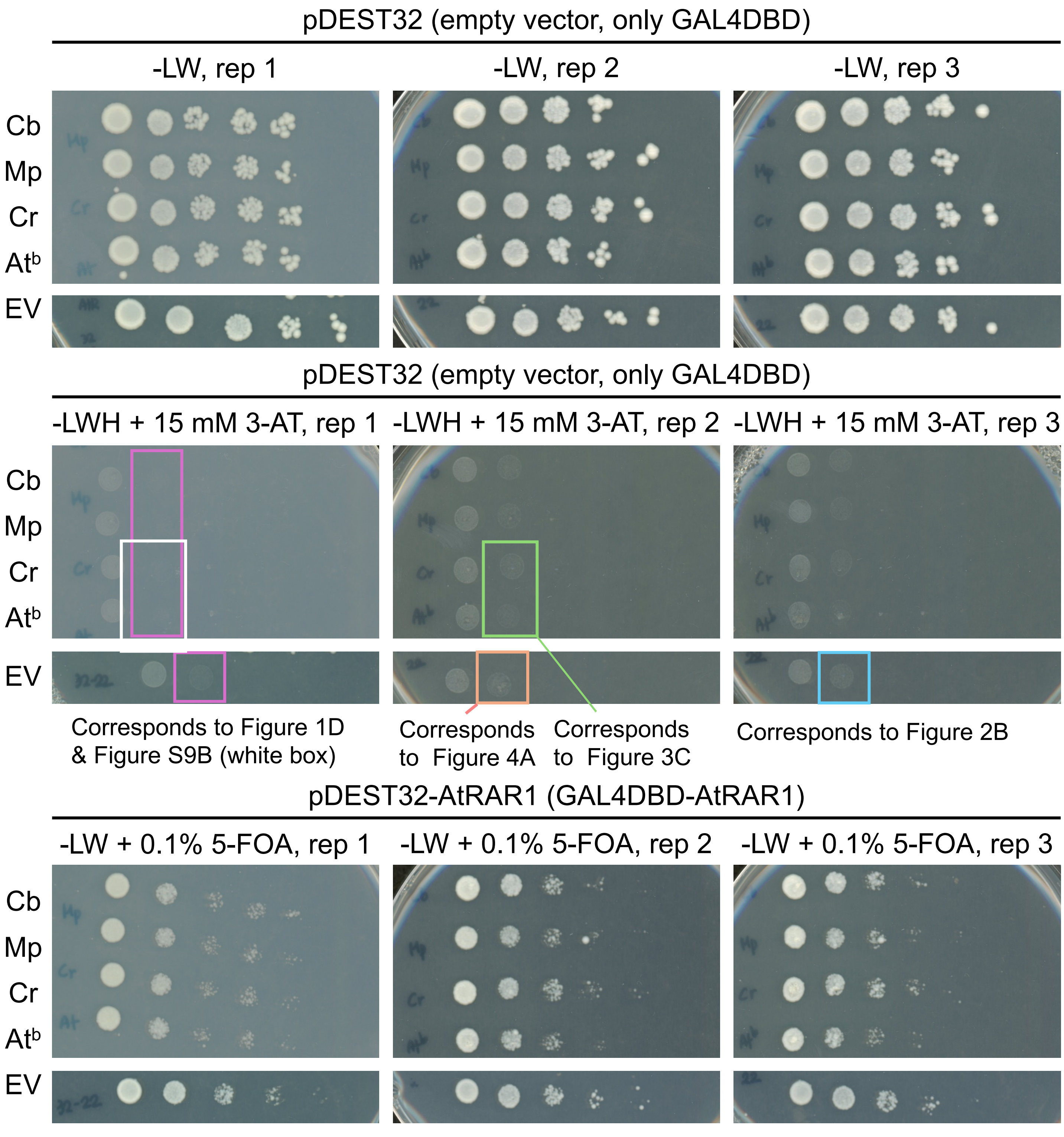


Cb; CbSGT1-pDEST22
Mp; MpSGT1-pDEST22
Cr; CrSGT1-pDEST22
$At^b$; pDEST-AD-SGT1b
EV; empty pDEST22

(serial dilution was started to from $OD_{600}$ = 1.0 and finished at $OD_{600}$ = 1.0 × $10^{-4}$)

(every plate was scanned three days after inoculation)

#### Data S1 – Raw data and extended yeast two-hybrid analyses

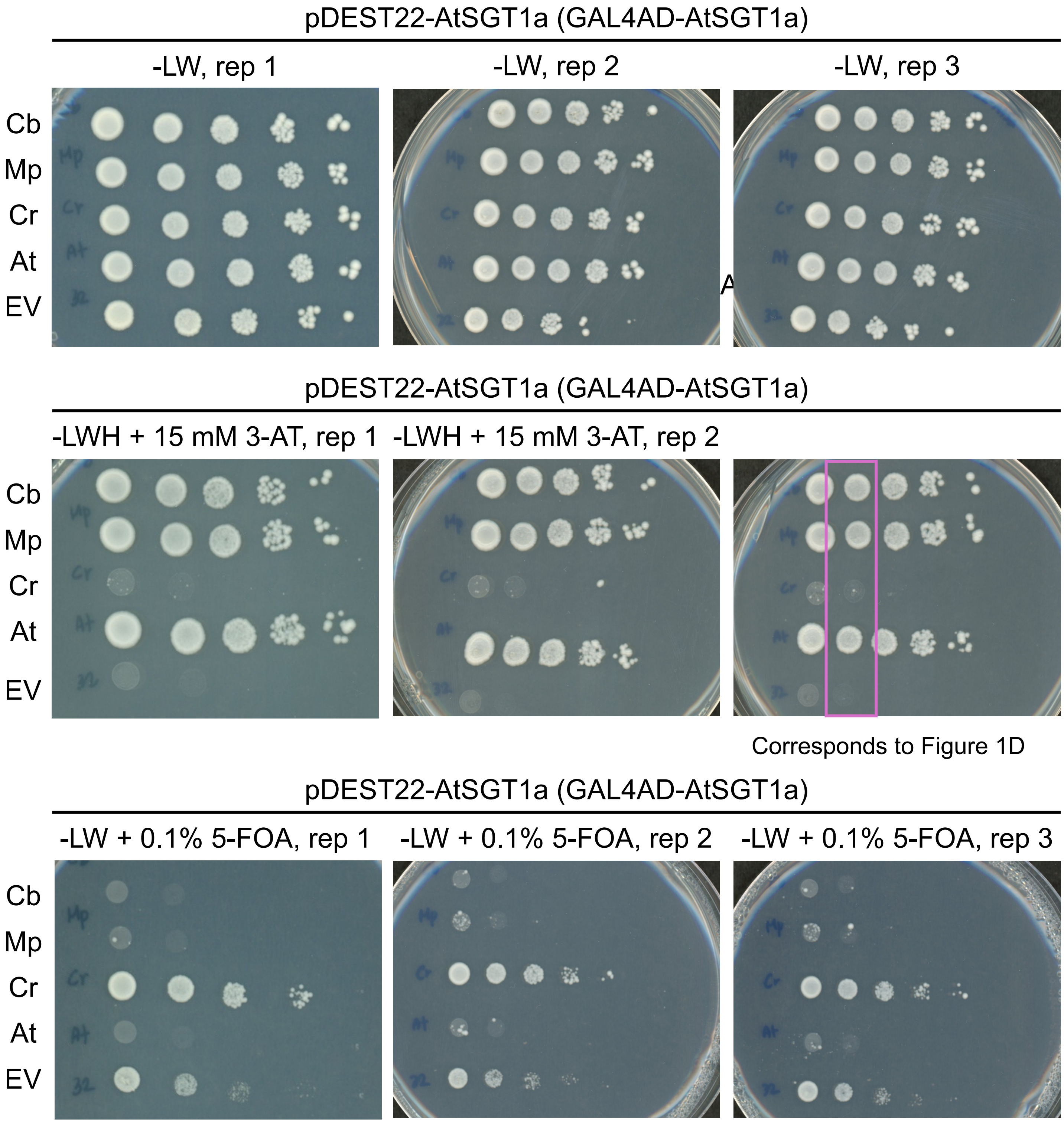


Cb; CbRAR1-pDEST32
Mp; MpRAR1-pDEST32
Cr; CrRAR1-pDEST32
At; AtRAR1-pDEST32
EV; empty pDEST32

(every RAR1 homolog was cloned into pDEST32 (GAL4DBD))

(serial dilution was started to from $OD_{600}$ = 1.0 and finished at $OD_{600}$ = $1.0 \times 10^{-4}$)

(every plate was scanned three days after inoculation)

#### Data S1 – Raw data and extended yeast two-hybrid analyses

pDEST32-CrRAR1$^{AtCHORDII}$ (GAL4DBD-CrRAR1$^{AtCHORDII}$)

-LW, rep 1 | -LW, rep 2 | -LW, rep 3

Cr
$At^b$
EV

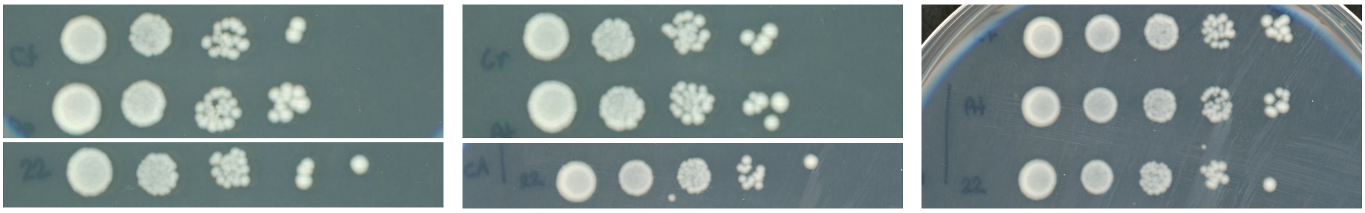

pDEST32-CrRAR1$^{AtCHORDII}$ (GAL4DBD-CrRAR1$^{AtCHORDII}$)

-LWH + 15 mM 3-AT, rep 1 | -LWH + 15 mM 3-AT, rep 2 | -LWH + 15 mM 3-AT, rep 3

Cr
$At^b$
EV

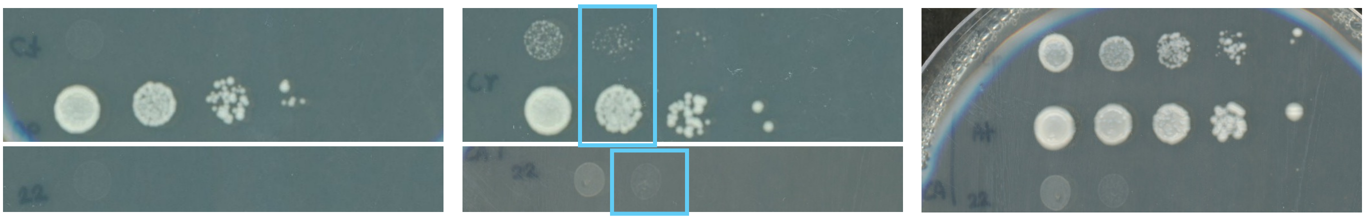

Corresponds to Figure 2B

pDEST32-CrRAR1$^{AtCHORDII}$ (GAL4DBD-CrRAR1$^{AtCHORDII}$)

-LW + 0.1% 5-FOA, rep 1 | -LW + 0.1% 5-FOA, rep 2 | -LW + 0.1% 5-FOA, rep 3

Cr
$At^b$
EV

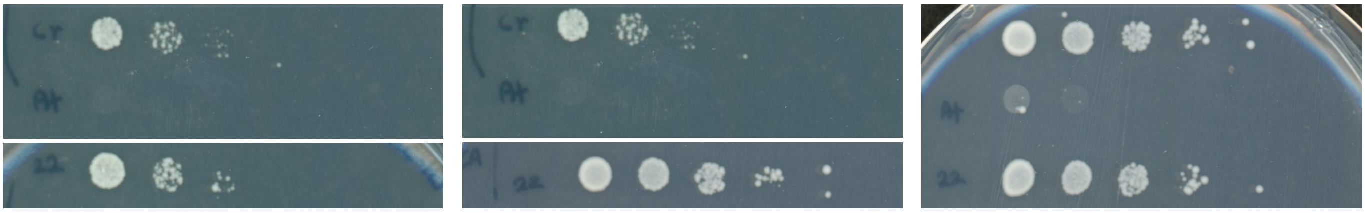

Cr; CrSGT1-pDEST22
$At^b$; pDEST-AD-AtSGT1b
EV; empty pDEST22

(serial dilution was started to from $OD_{600}$ = 1.0 and finished at $OD_{600} = 1.0 \times 10^{-4}$)

(every plate was scanned three days after inoculation)

#### Data S1 – Raw data and extended yeast two-hybrid analyses

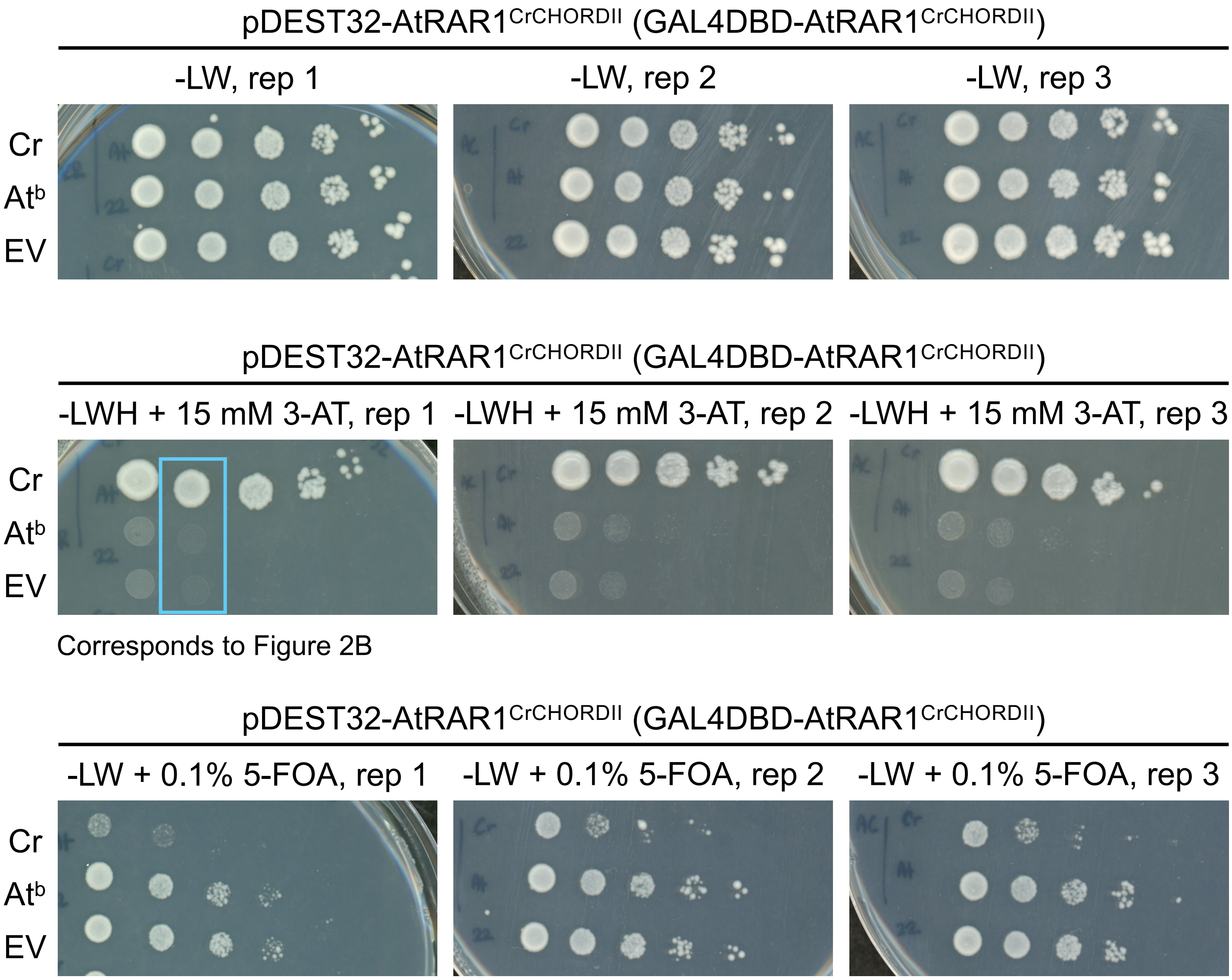


Cr; CrSGT1-pDEST22
At$^{b}$; pDEST-AD-AtSGT1b
EV; empty pDEST22

(every SGT1 homolog was cloned into pDEST22 (GAL4AD), except AtSGT1b (pDEST-AD-AtSGT1b))

(serial dilution was started to from $OD_{600}$ = 1.0 and finished at $OD_{600}$ = 1.0 × $10^{-4}$)

(every plate was scanned three days after inoculation)

#### Data S1 – Raw data and extended yeast two-hybrid analyses

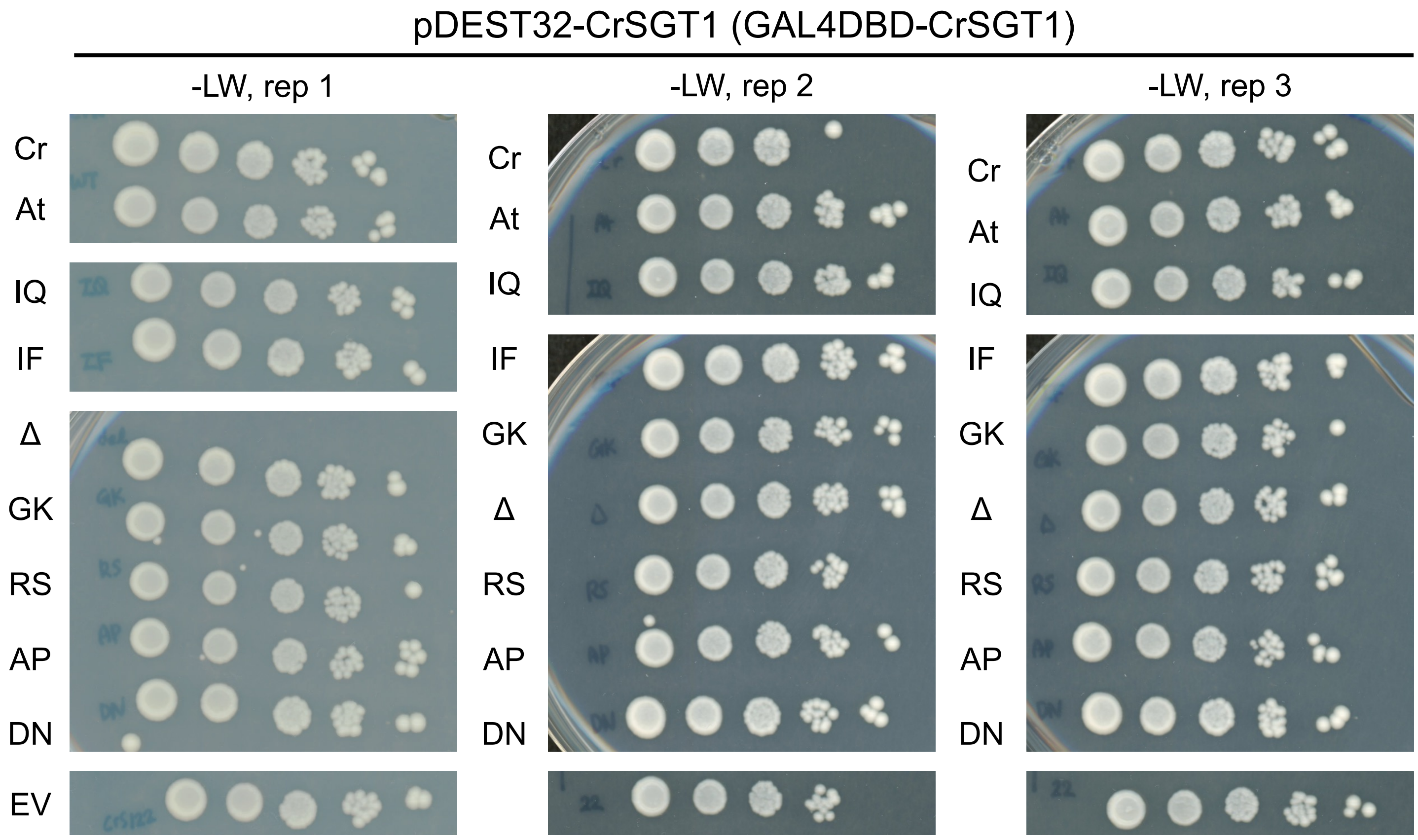


Cr; CrRAR1-pDEST22
At; AtRAR1-pDEST22
IQ; AtRAR1$^{I153Q}$-pDEST22
IF; AtRAR1$^{I153F}$-pDEST22
Δ; AtRAR1$^{\Delta K162G163}$-pDEST22
GK; AtRAR1$^{G165K}$-pDEST22
RS; AtRAR1$^{R171S}$-pDEST22
AP; AtRAR1$^{A185P}$-pDEST22
DN; AtRAR1$^{D198N}$-pDEST22
EV; empty pDEST22

(every RAR1 homolog and variant was cloned into pDEST22 (GAL4AD))

(serial dilution was started to from $OD_{600}$ = 1.0 and finished at $OD_{600}$ = 1.0 × $10^{-4}$)

(every plate was scanned three days after inoculation)

#### Data S1 – Raw data and extended yeast two-hybrid analyses

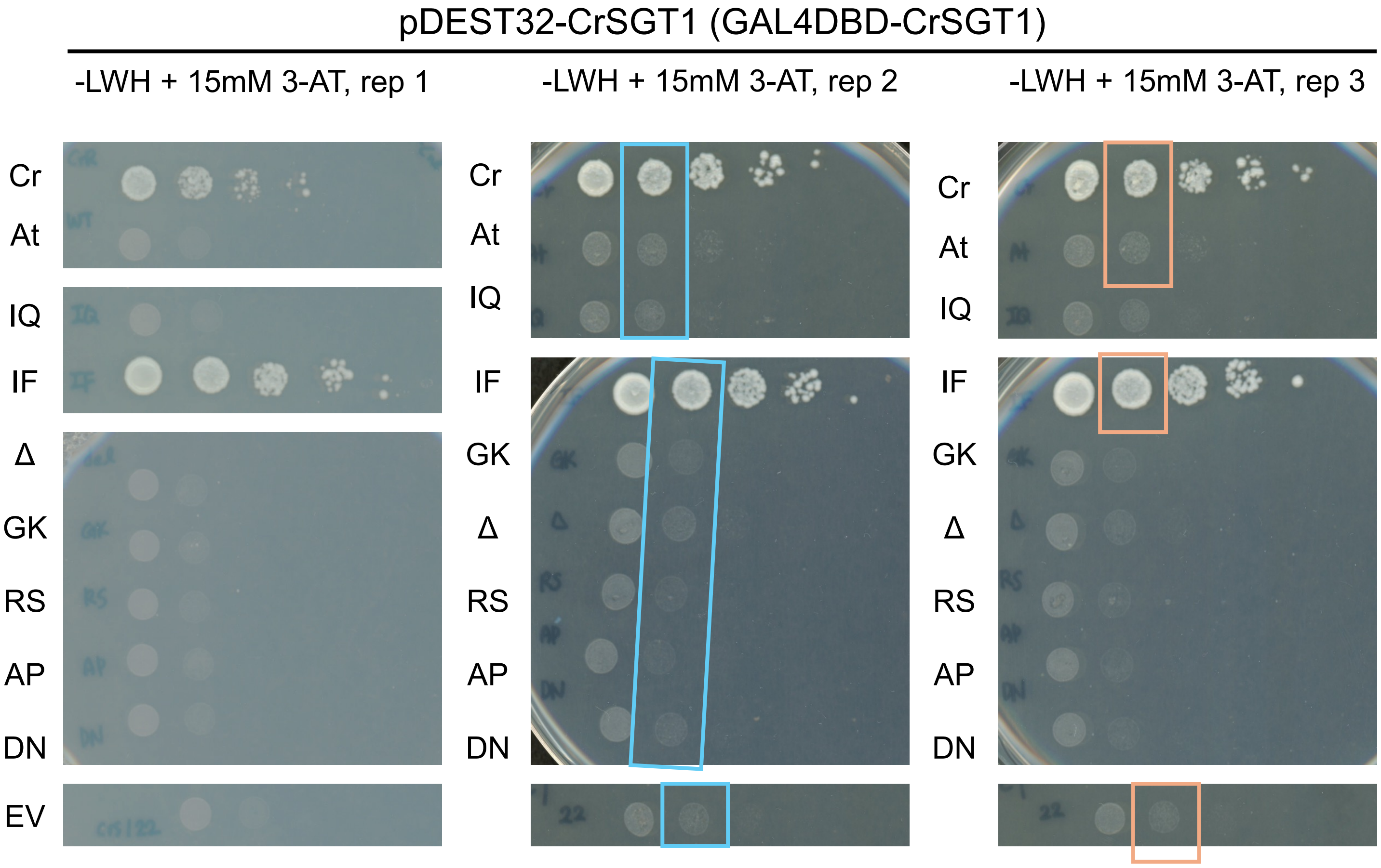


Corresponds to Figure 2D

Corresponds to Figure 4A

Cr; CrRAR1-pDEST22
At; AtRAR1-pDEST22
IQ; AtRAR1$^{I153Q}$-pDEST22
IF; AtRAR1$^{I153F}$-pDEST22
Δ; AtRAR1$^{\Delta K162G163}$-pDEST22
GK; AtRAR1$^{G165K}$-pDEST22
RS; AtRAR1$^{R171S}$-pDEST22
AP; AtRAR1$^{A185P}$-pDEST22
DN; AtRAR1$^{D198N}$-pDEST22
EV; empty pDEST22

(every RAR1 homolog and variant was cloned into pDEST22 (GAL4AD))

(serial dilution was started to from $OD_{600}$ = 1.0 and finished at $OD_{600}$ = 1.0 × $10^{-4}$)

(every plate was scanned three days after inoculation)

### Data S1 – Raw data and extended yeast two-hybrid analyses

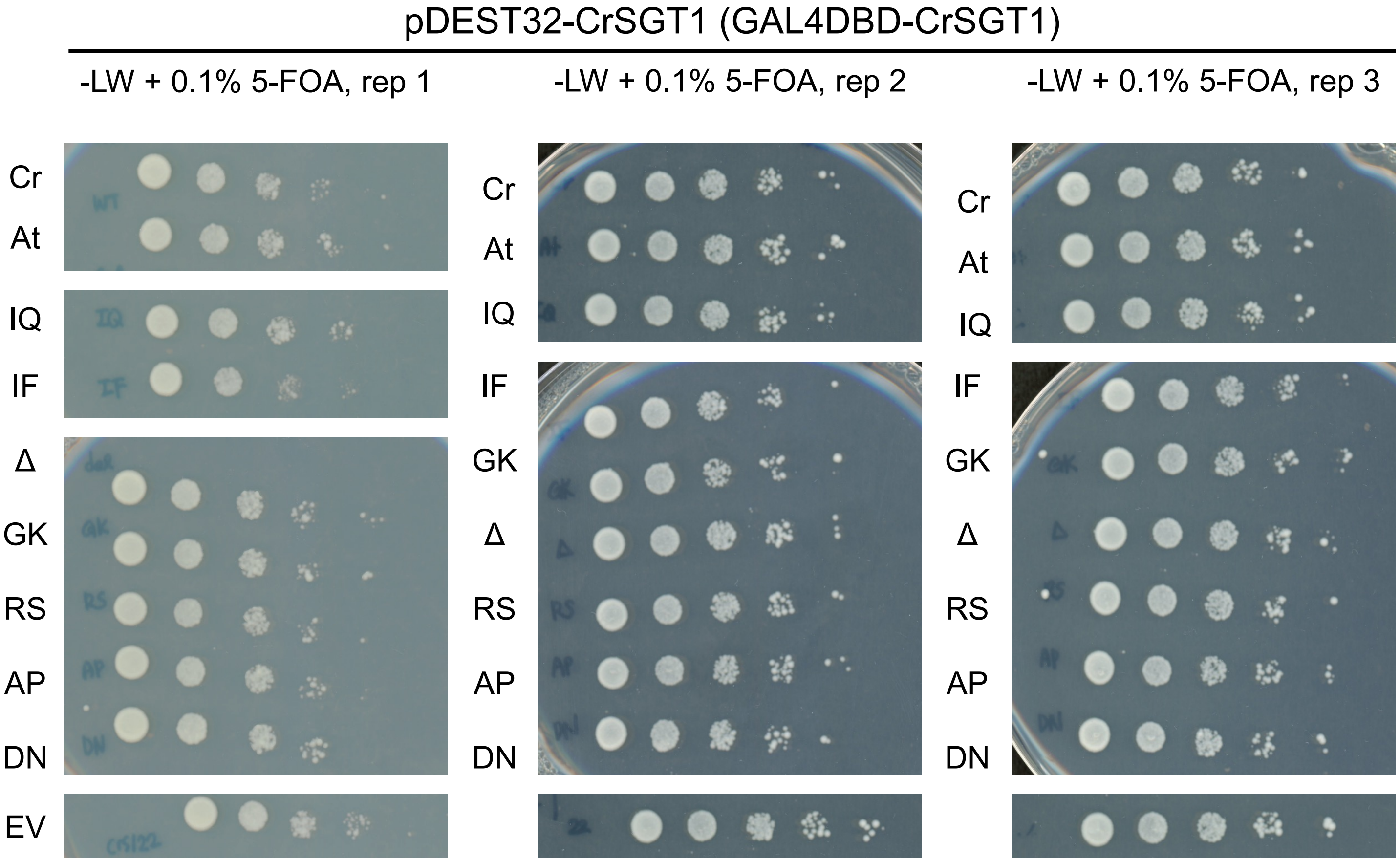


Cr; CrRAR1-pDEST22
At; AtRAR1-pDEST22
IQ; AtRAR1$^{I153Q}$-pDEST22
IF; AtRAR1$^{I153F}$-pDEST22
Δ; AtRAR1$^{\Delta K162G163}$-pDEST22
GK; AtRAR1$^{G165K}$-pDEST22
RS; AtRAR1$^{R171S}$-pDEST22
AP; AtRAR1$^{A185P}$-pDEST22
DN; AtRAR1$^{D198N}$-pDEST22
EV; empty pDEST22

(every RAR1 homolog and variant was cloned into pDEST22 (GAL4AD))

(serial dilution was started to from $OD_{600}$ = 1.0 and finished at $OD_{600}$ = 1.0 × $10^{-4}$)

(every plate was scanned three days after inoculation)

### Data S1 – Raw data and extended yeast two-hybrid analyses

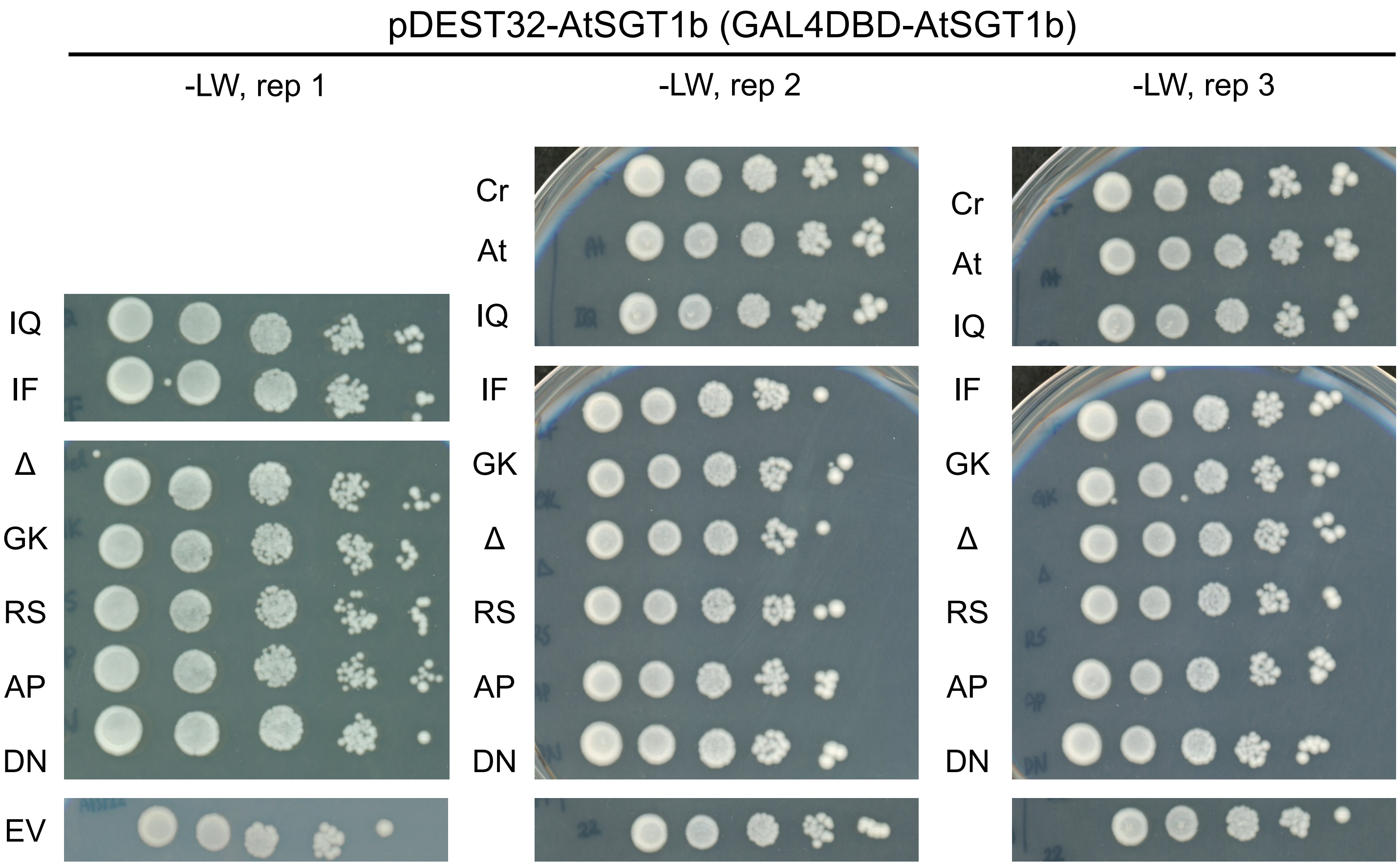


Cr; CrRAR1-pDEST22
At; AtRAR1-pDEST22
IQ; AtRAR1$^{I153Q}$-pDEST22
IF; AtRAR1$^{I153F}$-pDEST22
Δ; AtRAR1$^{\Delta K162G163}$-pDEST22
GK; AtRAR1$^{G165K}$-pDEST22
RS; AtRAR1$^{R171S}$-pDEST22
AP; AtRAR1$^{A185P}$-pDEST22
DN; AtRAR1$^{D198N}$-pDEST22
EV; empty pDEST22

(every RAR1 homolog and variant was cloned into pDEST22 (GAL4AD))

(serial dilution was started to from $OD_{600}$ = 1.0 and finished at $OD_{600}$ = 1.0 × $10^{-4}$)

(every plate was scanned three days after inoculation)

### Data S1 – Raw data and extended yeast two-hybrid analyses

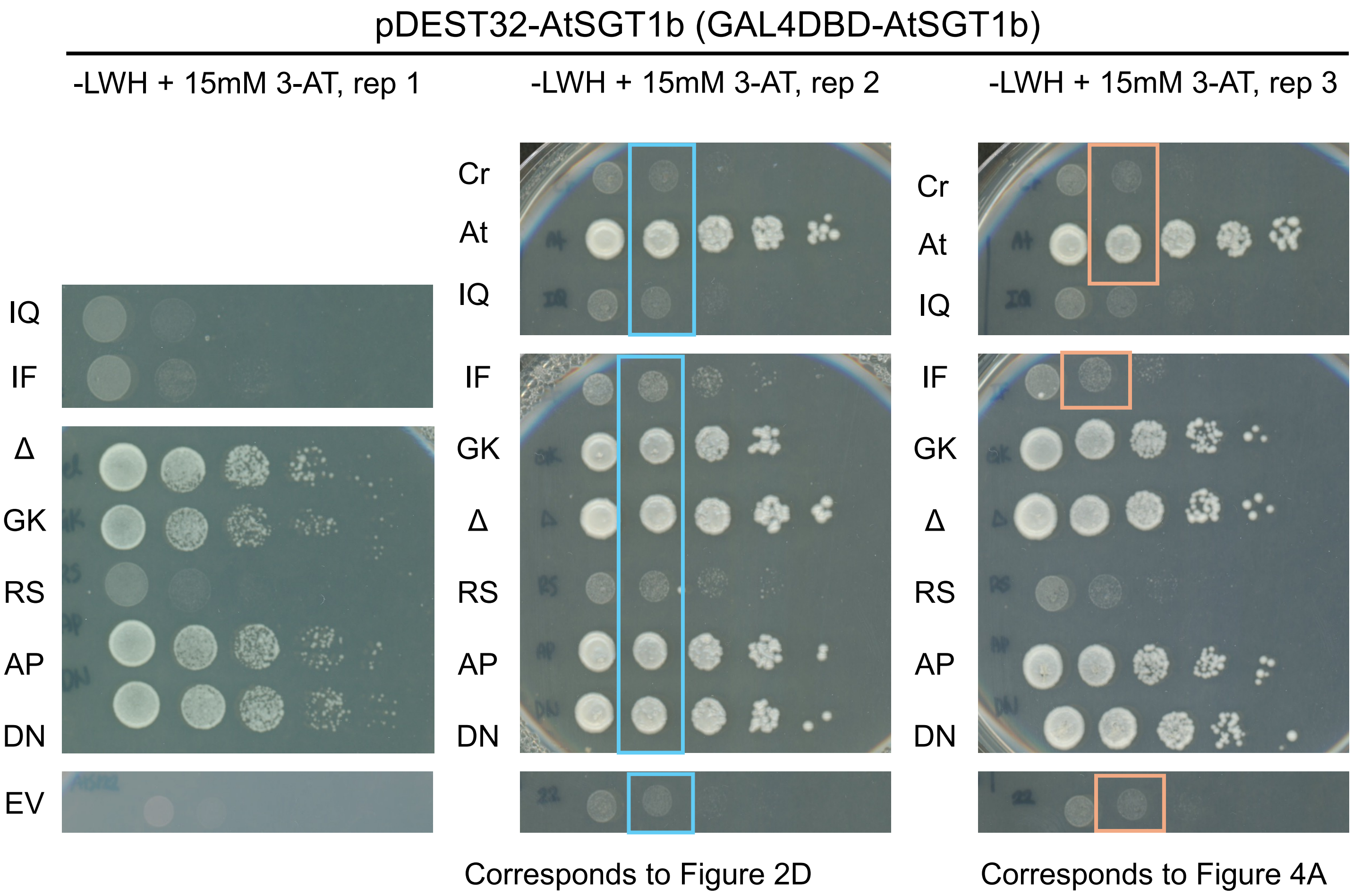


Cr; CrRAR1-pDEST22
At; AtRAR1-pDEST22
IQ; AtRAR1$^{I153Q}$-pDEST22
IF; AtRAR1$^{I153F}$-pDEST22
Δ; AtRAR1$^{\Delta K162G163}$-pDEST22
GK; AtRAR1$^{G165K}$-pDEST22
RS; AtRAR1$^{R171S}$-pDEST22
AP; AtRAR1$^{A185P}$-pDEST22
DN; AtRAR1$^{D198N}$-pDEST22
EV; empty pDEST22

(every RAR1 homolog and variant was cloned into pDEST22 (GAL4AD))

(serial dilution was started to from $OD_{600}$ = 1.0 and finished at $OD_{600}$ = 1.0 × $10^{-4}$)

(every plate was scanned three days after inoculation)

#### Data S1 – Raw data and extended yeast two-hybrid analyses

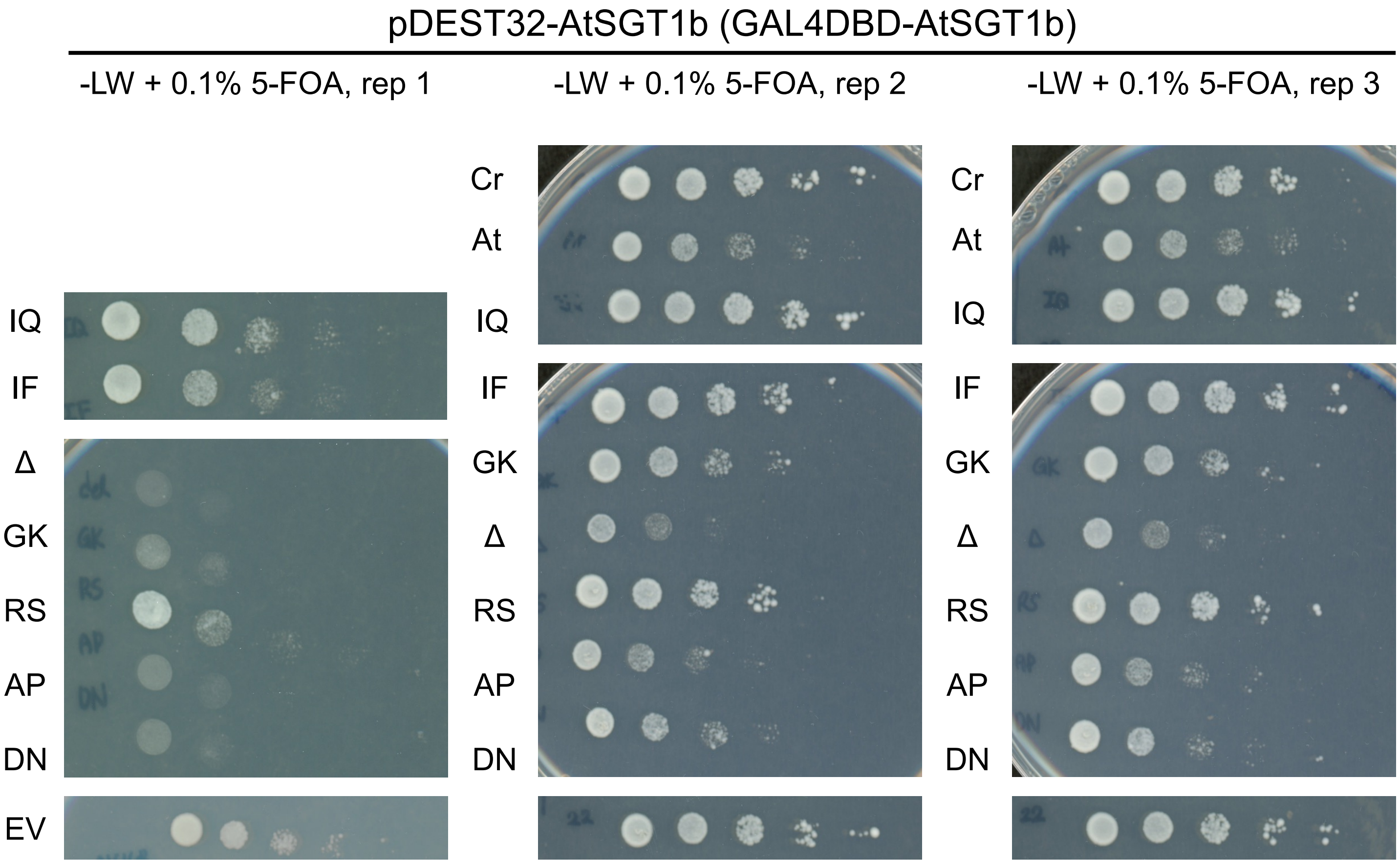


Cr; CrRAR1-pDEST22
At; AtRAR1-pDEST22
IQ; AtRAR1$^{I153Q}$-pDEST22
IF; AtRAR1$^{I153F}$-pDEST22
Δ; AtRAR1$^{\Delta K162G163}$-pDEST22
GK; AtRAR1$^{G165K}$-pDEST22
RS; AtRAR1$^{R171S}$-pDEST22
AP; AtRAR1$^{A185P}$-pDEST22
DN; AtRAR1$^{D198N}$-pDEST22
EV; empty pDEST22

(every RAR1 homolog and variant was cloned into pDEST22 (GAL4AD))

(serial dilution was started to from $OD_{600}$ = 1.0 and finished at $OD_{600}$ = 1.0 × $10^{-4}$)

(every plate was scanned three days after inoculation)

#### Data S1 – Raw data and extended yeast two-hybrid analyses

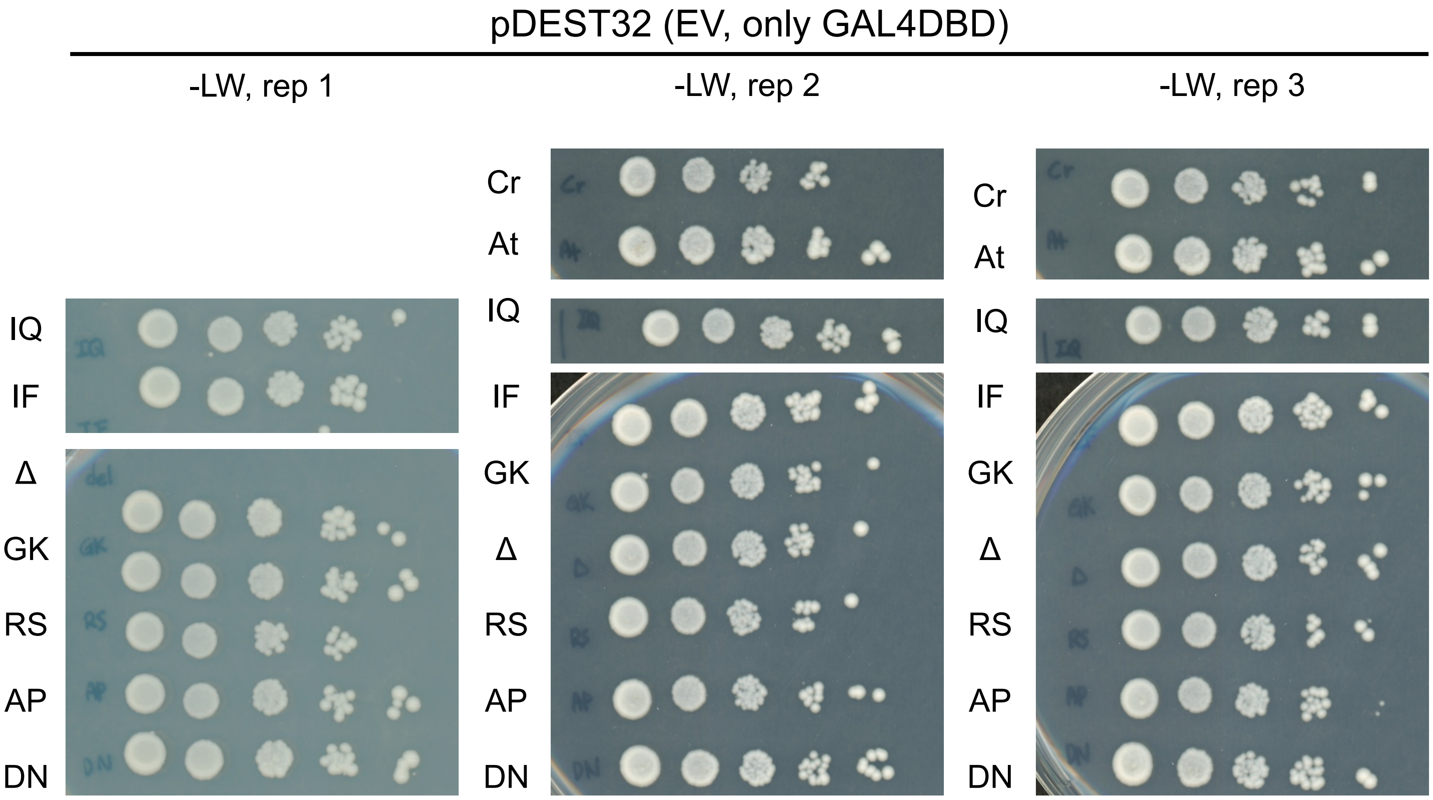


Cr; CrRAR1-pDEST22
At; AtRAR1-pDEST22
IQ; AtRAR1$^{I153Q}$-pDEST22
IF; AtRAR1$^{I153F}$-pDEST22
Δ; AtRAR1$^{\Delta K162G163}$-pDEST22
GK; AtRAR1$^{G165K}$-pDEST22
RS; AtRAR1$^{R171S}$-pDEST22
AP; AtRAR1$^{A185P}$-pDEST22
DN; AtRAR1$^{D198N}$-pDEST22
EV; empty pDEST22

(every RAR1 homolog and variant was cloned into pDEST22 (GAL4AD))

(serial dilution was started to from $OD_{600}$ = 1.0 and finished at $OD_{600}$ = 1.0 × $10^{-4}$)

(every plate was scanned three days after inoculation)

#### Data S1 – Raw data and extended yeast two-hybrid analyses

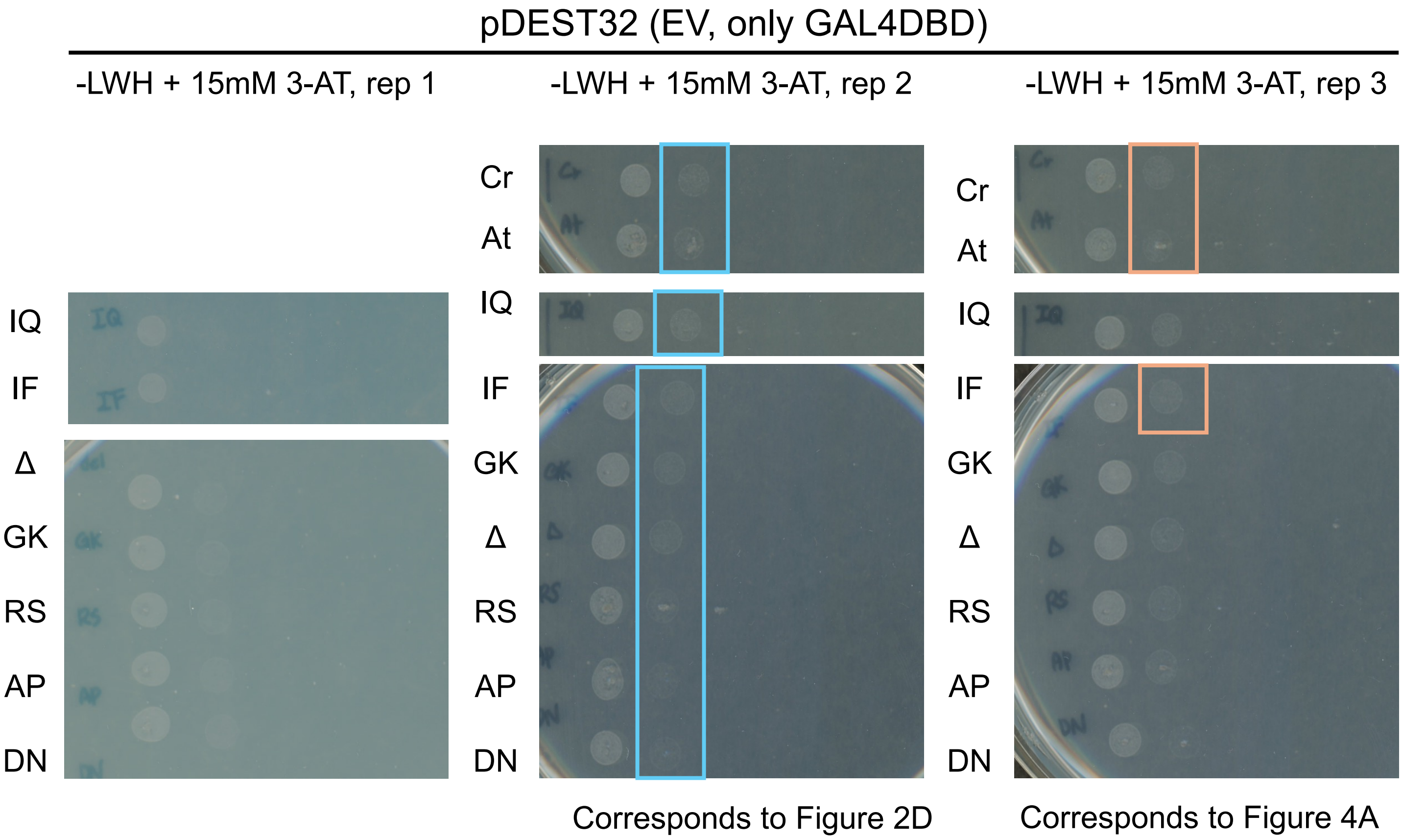


Cr; CrRAR1-pDEST22
At; AtRAR1-pDEST22
IQ; AtRAR1$^{I153Q}$-pDEST22
IF; AtRAR1$^{I153F}$-pDEST22
Δ; AtRAR1$^{\Delta K162G163}$-pDEST22
GK; AtRAR1$^{G165K}$-pDEST22
RS; AtRAR1$^{R171S}$-pDEST22
AP; AtRAR1$^{A185P}$-pDEST22
DN; AtRAR1$^{D198N}$-pDEST22
EV; empty pDEST22

(every RAR1 homolog and variant was cloned into pDEST22 (GAL4AD))

(serial dilution was started to from $OD_{600}$ = 1.0 and finished at $OD_{600}$ = 1.0 × $10^{-4}$)

(every plate was scanned three days after inoculation)

### Data S1 – Raw data and extended yeast two-hybrid analyses

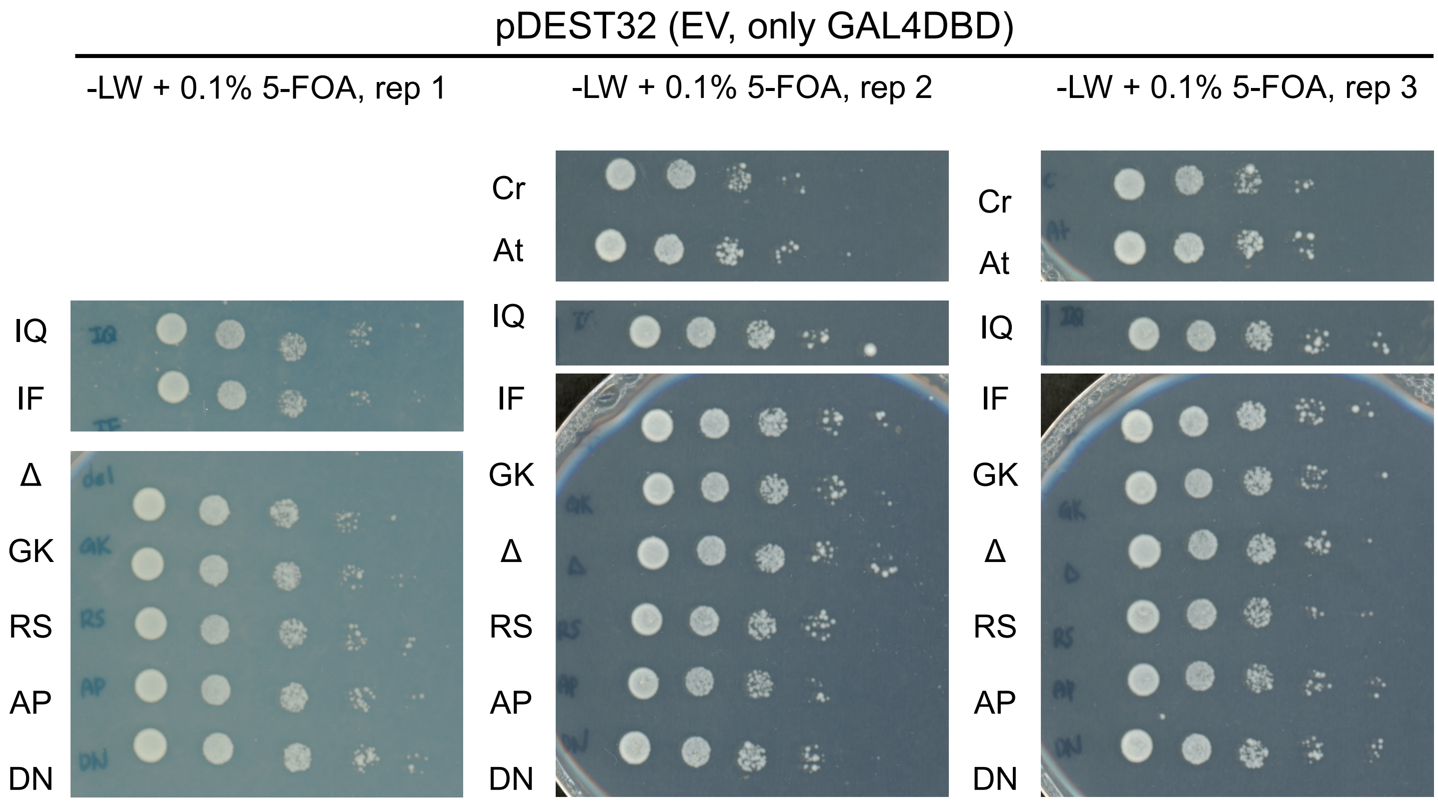


Cr; CrRAR1-pDEST22
At; AtRAR1-pDEST22
IQ; AtRAR1$^{I153Q}$ -pDEST22
IF; AtRAR1$^{I153F}$-pDEST22
Δ; AtRAR1$^{\Delta\ K162G163}$-pDEST22
GK; AtRAR1$^{G165K}$-pDEST22
RS; AtRAR1$^{R171S}$-pDEST22
AP; AtRAR1$^{A185P}$-pDEST22
DN; AtRAR1$^{D198N}$-pDEST22
EV; empty pDEST22

(every RAR1 homolog and variant was cloned into pDEST22 (GAL4AD))

(serial dilution was started to from $OD_{600}$ = 1.0 and finished at $OD_{600}$ = 1.0 × $10^{-4}$)

(every plate was scanned three days after inoculation)

### Data S1 – Raw data and extended yeast two-hybrid analyses

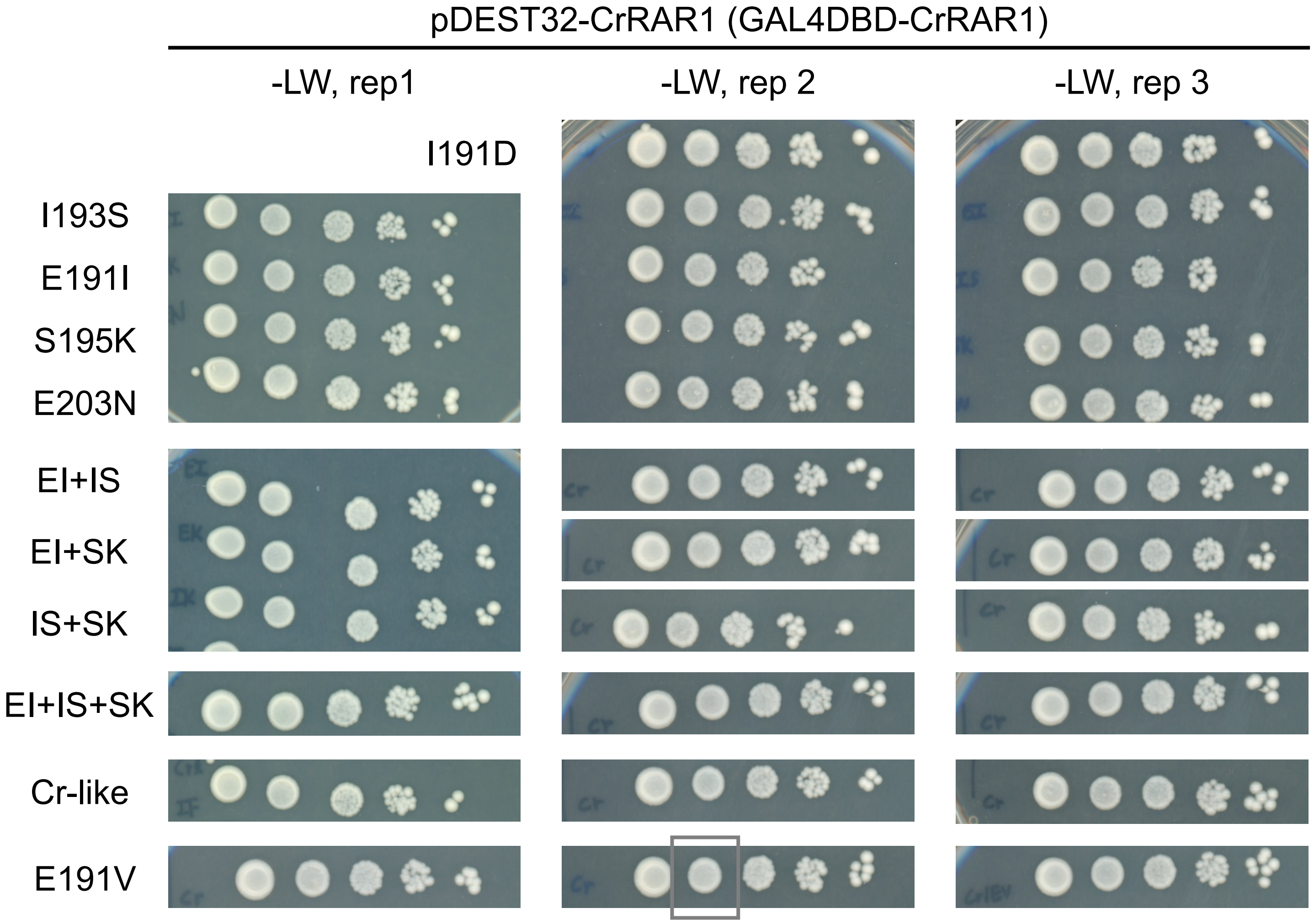


Corresponds to Figure S5

I191D; AtSGT1b$^{ID}$-pDEST22
I193S; AtSGT1b$^{IS}$-pDEST22
E191I; AtSGT1b$^{EI}$ -pDEST22
S195K; AtSGT1b$^{SK}$-pDEST22
E203N; AtSGT1b$^{EN}$-pDEST22
EI+IS; AtSGT1b$^{EI+IS}$-pDEST22
EI+SK; AtSGT1b$^{EI+SK}$-pDEST22
IS+SK; AtSGT1b$^{IS+SK}$-pDEST22
EI+IS+SK; AtSGT1b$^{EI+IS+SK}$-pDEST22
Cr-like; AtSGT1b$^{Cr\text{-}like}$-pDEST22
E191V; AtSGT1b$^{EV}$-pDEST22

(every SGT1 homolog and variant was cloned into pDEST22 (GAL4AD))

(serial dilution was started to from $OD_{600}$ = 1.0 and finished at $OD_{600}$ = $1.0 \times 10^{-4}$)

(every plate was scanned three days after inoculation)

#### Data S1 – Raw data and extended yeast two-hybrid analyses

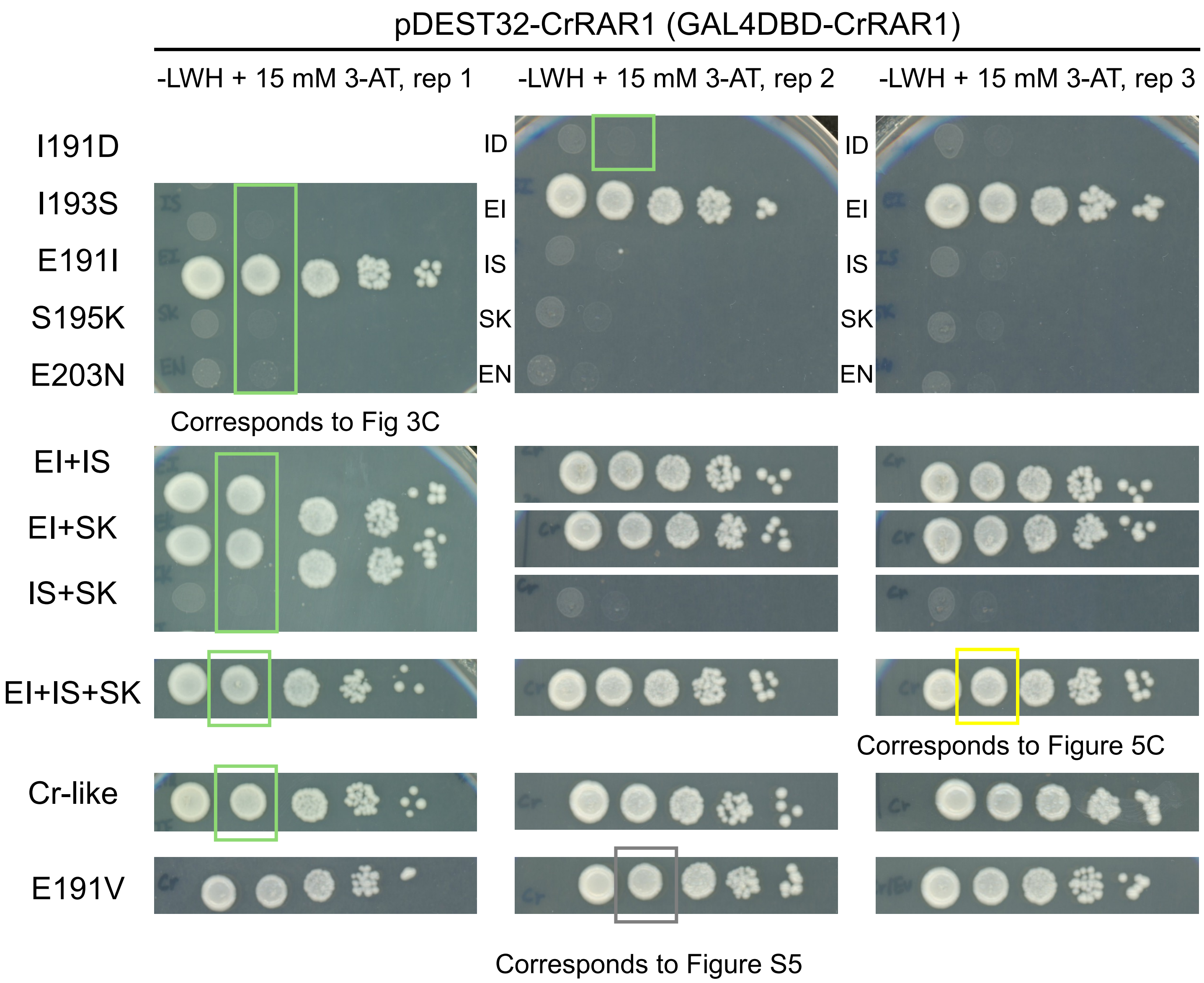


I191D; AtSGT1b$^{ID}$-pDEST22
I193S; AtSGT1b$^{IS}$-pDEST22
E191I; AtSGT1b$^{EI}$ -pDEST22
S195K; AtSGT1b$^{SK}$-pDEST22
E203N; AtSGT1b$^{EN}$-pDEST22
EI+IS; AtSGT1b$^{EI+IS}$-pDEST22
EI+SK; AtSGT1b$^{EI+SK}$-pDEST22
IS+SK; AtSGT1b$^{IS+SK}$-pDEST22
EI+IS+SK; AtSGT1b$^{EI+IS+SK}$-pDEST22
Cr-like; AtSGT1b$^{Cr\text{-}like}$-pDEST22
E191V; AtSGT1b$^{EV}$-pDEST22

(every SGT1 homolog and variant was cloned into pDEST22 (GAL4AD))

(serial dilution was started to from $OD_{600}$ = 1.0 and finished at $OD_{600}$ = 1.0 × $10^{-4}$)

(every plate was scanned three days after inoculation)

#### Data S1 – Raw data and extended yeast two-hybrid analyses

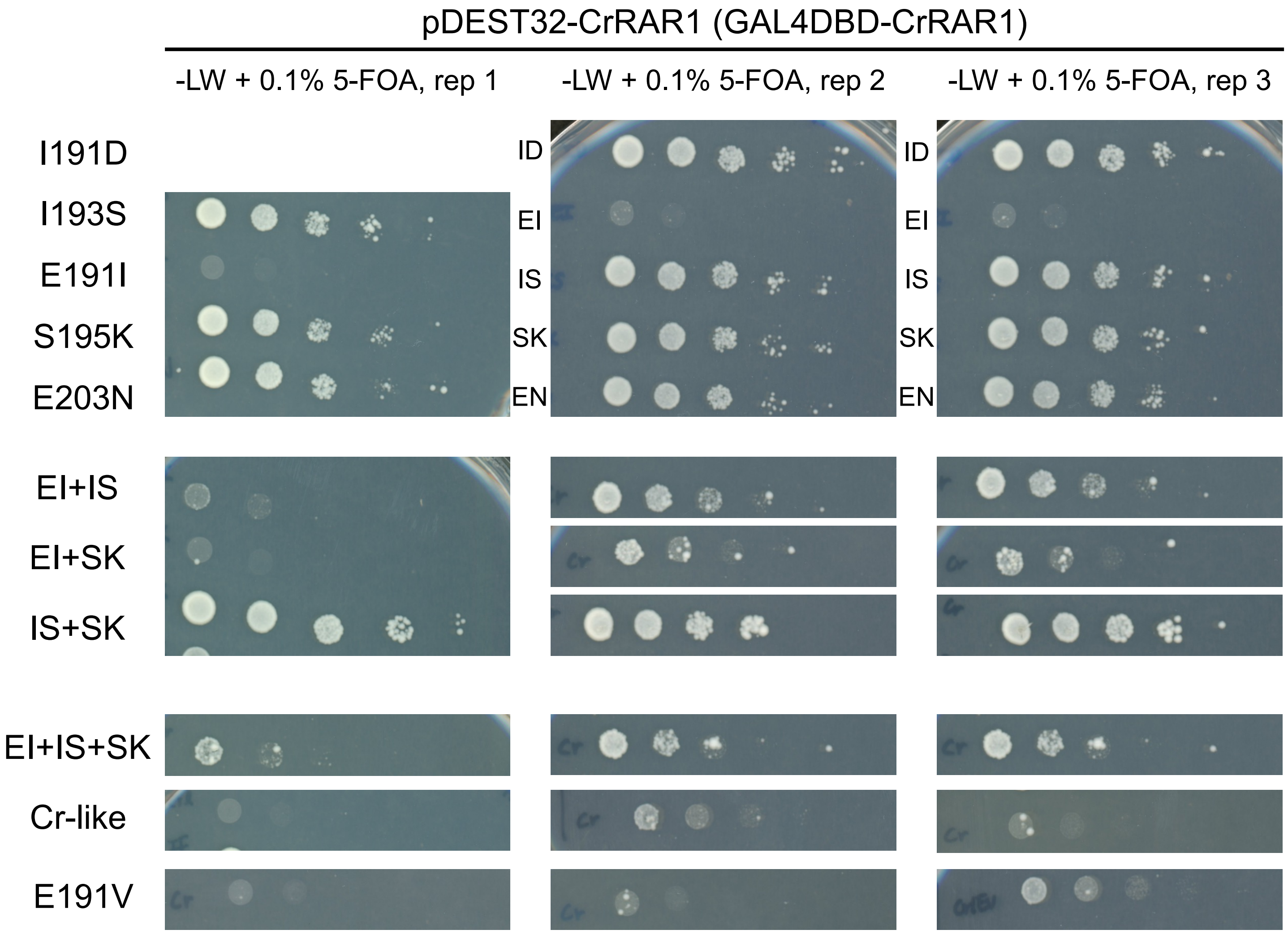


I191D; AtSGT1b$^{ID}$-pDEST22
I193S; AtSGT1b$^{IS}$-pDEST22
E191I; AtSGT1b$^{EI}$ -pDEST22
S195K; AtSGT1b$^{SK}$-pDEST22
E203N; AtSGT1b$^{EN}$-pDEST22
EI+IS; AtSGT1b$^{EI+IS}$-pDEST22
EI+SK; AtSGT1b$^{EI+SK}$-pDEST22
IS+SK; AtSGT1b$^{IS+SK}$-pDEST22
EI+IS+SK; AtSGT1b$^{EI+IS+SK}$-pDEST22
Cr-like; AtSGT1b$^{Cr\text{-}like}$-pDEST22
E191V; AtSGT1b$^{EV}$-pDEST22

(every SGT1 homolog and variant was cloned into pDEST22 (GAL4AD))

(serial dilution was started to from $OD_{600}$ = 1.0 and finished at $OD_{600}$ = 1.0 × $10^{-4}$)

(every plate was scanned three days after inoculation)

### Data S1 – Raw data and extended yeast two-hybrid analyses

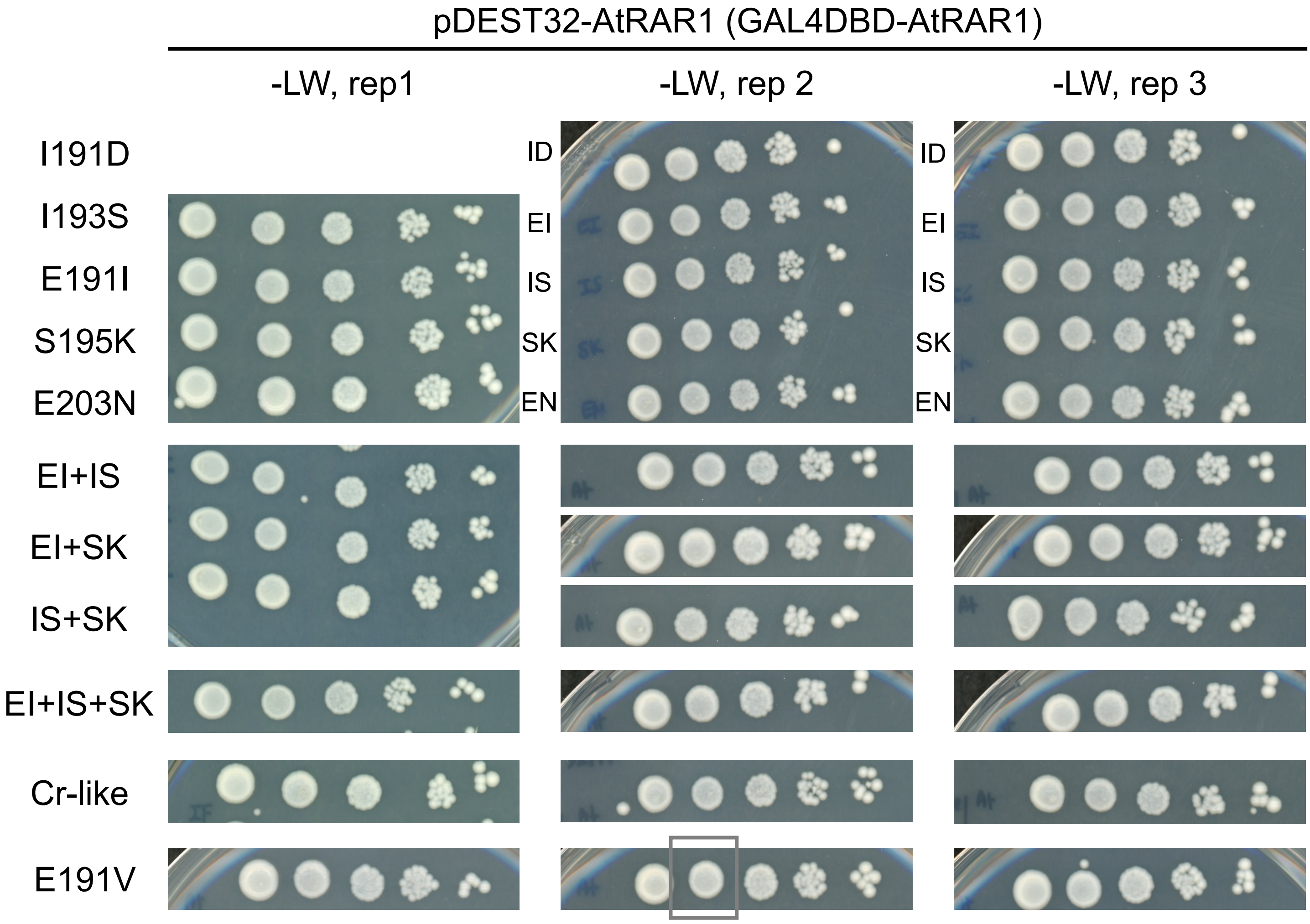


Corresponds to Figure S5

I191D; AtSGT1b$^{ID}$-pDEST22
I193S; AtSGT1b$^{IS}$-pDEST22
E191I; AtSGT1b$^{EI}$ -pDEST22
S195K; AtSGT1b$^{SK}$-pDEST22
E203N; AtSGT1b$^{EN}$-pDEST22
EI+IS; AtSGT1b$^{EI+IS}$-pDEST22
EI+SK; AtSGT1b$^{EI+SK}$-pDEST22
IS+SK; AtSGT1b$^{IS+SK}$-pDEST22
EI+IS+SK; AtSGT1b$^{EI+IS+SK}$-pDEST22
Cr-like; AtSGT1b$^{Cr\text{-}like}$-pDEST22
E191V; AtSGT1b$^{EV}$-pDEST22

(every SGT1 homolog and variant was cloned into pDEST22 (GAL4AD))

(serial dilution was started to from $OD_{600}$ = 1.0 and finished at $OD_{600}$ = 1.0 × $10^{-4}$)

(every plate was scanned three days after inoculation)

#### Data S1 – Raw data and extended yeast two-hybrid analyses

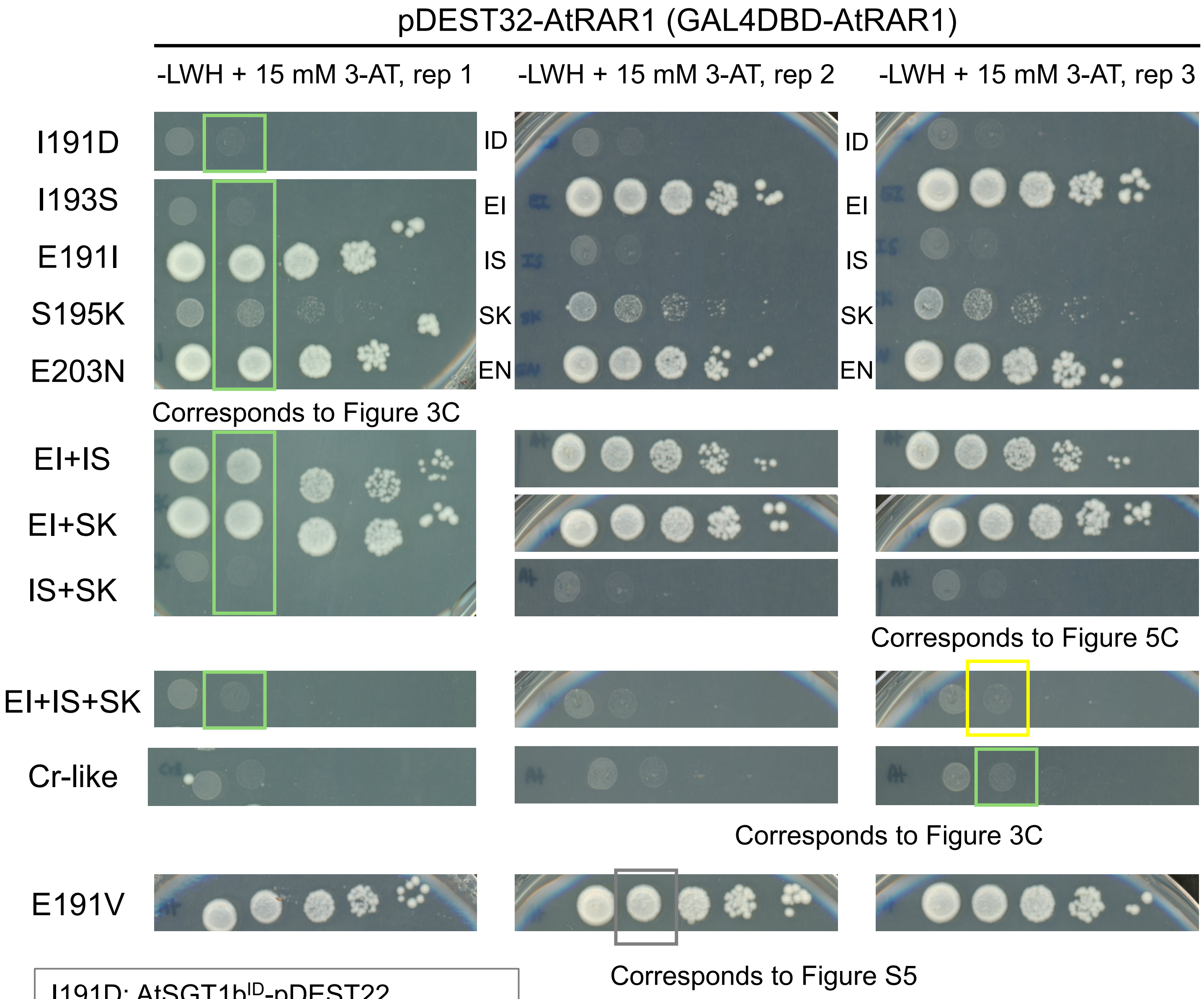


I191D; AtSGT1b$^{ID}$-pDEST22
I193S; AtSGT1b$^{IS}$-pDEST22
E191I; AtSGT1b$^{EI}$ -pDEST22
S195K; AtSGT1b$^{SK}$-pDEST22
E203N; AtSGT1b$^{EN}$-pDEST22
EI+IS; AtSGT1b$^{EI+IS}$-pDEST22
EI+SK; AtSGT1b$^{EI+SK}$-pDEST22
IS+SK; AtSGT1b$^{IS+SK}$-pDEST22
EI+IS+SK; AtSGT1b$^{EI+IS+SK}$-pDEST22
Cr-like; AtSGT1b$^{Cr\text{-}like}$-pDEST22
E191V; AtSGT1b$^{EV}$-pDEST22

(every SGT1 homolog and variant was cloned into pDEST22 (GAL4AD))

(serial dilution was started to from $OD_{600}$ = 1.0 and finished at $OD_{600}$ = 1.0 × $10^{-4}$)

(every plate was scanned three days after inoculation)

#### Data S1 – Raw data and extended yeast two-hybrid analyses

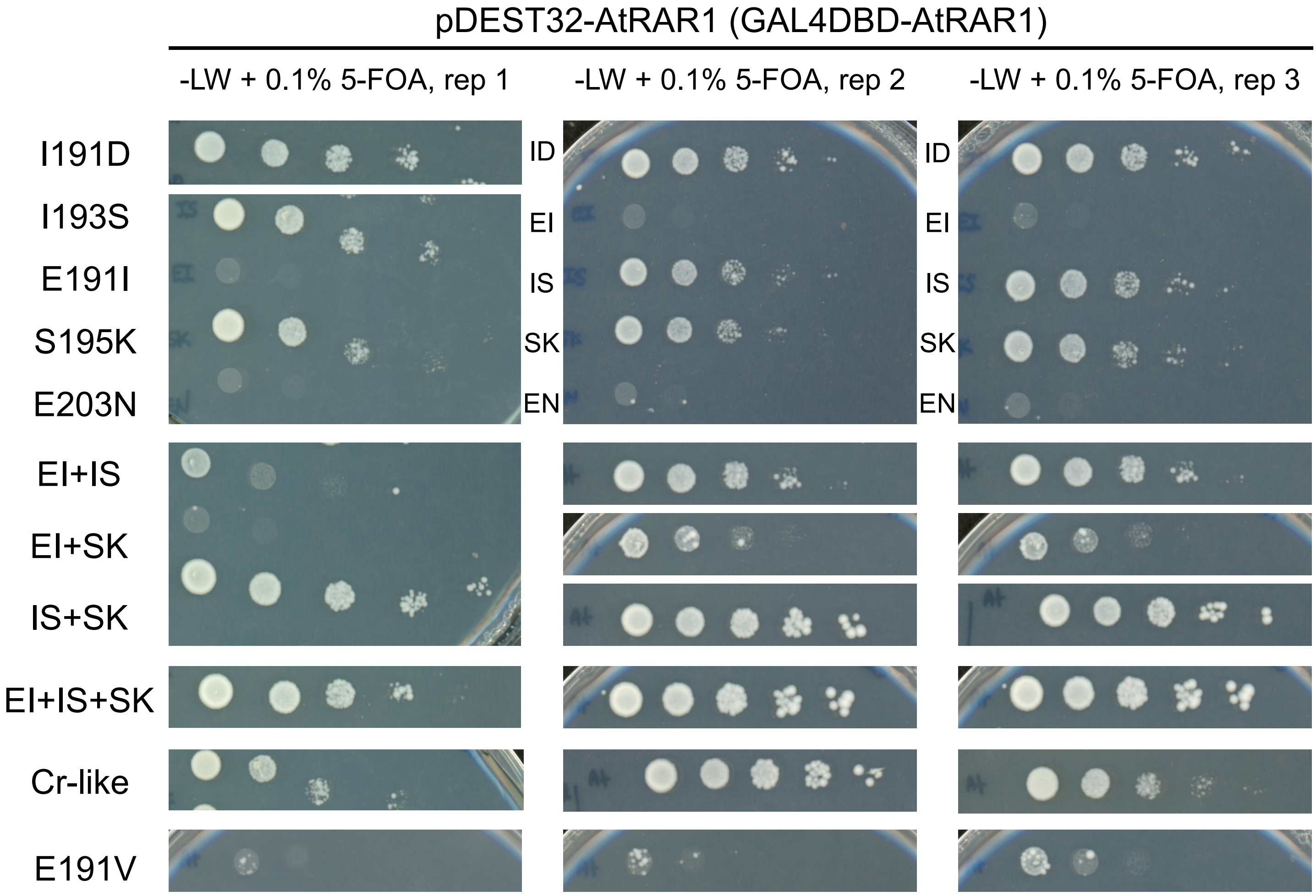


I191D; AtSGT1b$^{ID}$-pDEST22
I193S; AtSGT1b$^{IS}$-pDEST22
E191I; AtSGT1b$^{EI}$ -pDEST22
S195K; AtSGT1b$^{SK}$-pDEST22
E203N; AtSGT1b$^{EN}$-pDEST22
EI+IS; AtSGT1b$^{EI+IS}$-pDEST22
EI+SK; AtSGT1b$^{EI+SK}$-pDEST22
IS+SK; AtSGT1b$^{IS+SK}$-pDEST22
EI+IS+SK; AtSGT1b$^{EI+IS+SK}$-pDEST22
Cr-like; AtSGT1b$^{Cr\text{-}like}$-pDEST22
E191V; AtSGT1b$^{EV}$-pDEST22

(every SGT1 homolog and variant was cloned into pDEST22 (GAL4AD))

(serial dilution was started to from $OD_{600}$ = 1.0 and finished at $OD_{600}$ = 1.0 × $10^{-4}$)

(every plate was scanned three days after inoculation)

#### Data S1 – Raw data and extended yeast two-hybrid analyses

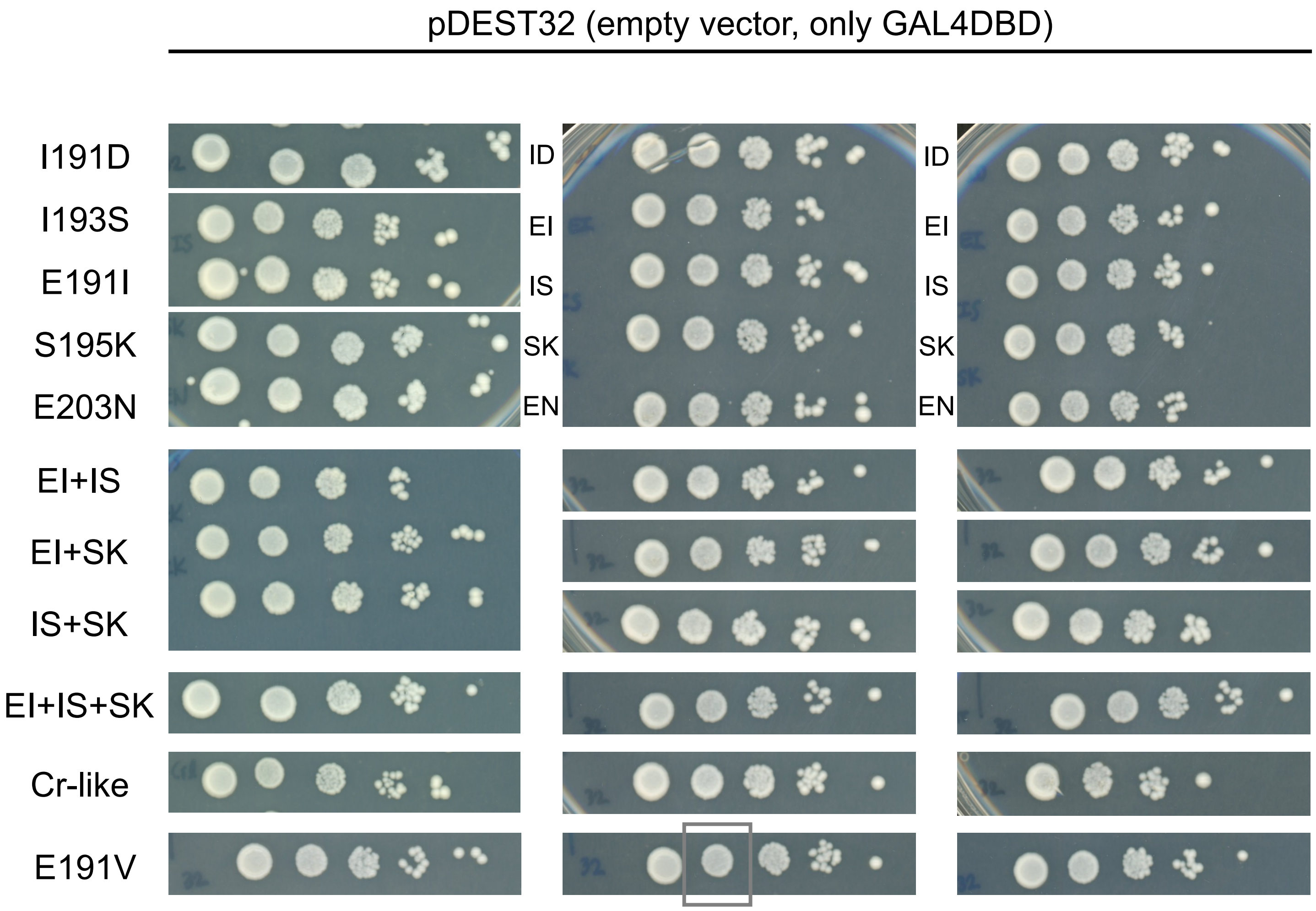


Corresponds to Figure S5

I191D; AtSGT1b$^{ID}$-pDEST22
I193S; AtSGT1b$^{IS}$-pDEST22
E191I; AtSGT1b$^{EI}$ -pDEST22
S195K; AtSGT1b$^{SK}$-pDEST22
E203N; AtSGT1b$^{EN}$-pDEST22
EI+IS; AtSGT1b$^{EI+IS}$-pDEST22
EI+SK; AtSGT1b$^{EI+SK}$-pDEST22
IS+SK; AtSGT1b$^{IS+SK}$-pDEST22
EI+IS+SK; AtSGT1b$^{EI+IS+SK}$-pDEST22
Cr-like; AtSGT1b$^{Cr\text{-}like}$-pDEST22
E191V; AtSGT1b$^{EV}$-pDEST22

(every SGT1 homolog and variant was cloned into pDEST22 (GAL4AD))

(serial dilution was started to from $OD_{600}$ = 1.0 and finished at $OD_{600}$ = 1.0 × $10^{-4}$)

(every plate was scanned three days after inoculation)

### Data S1 – Raw data and extended yeast two-hybrid analyses

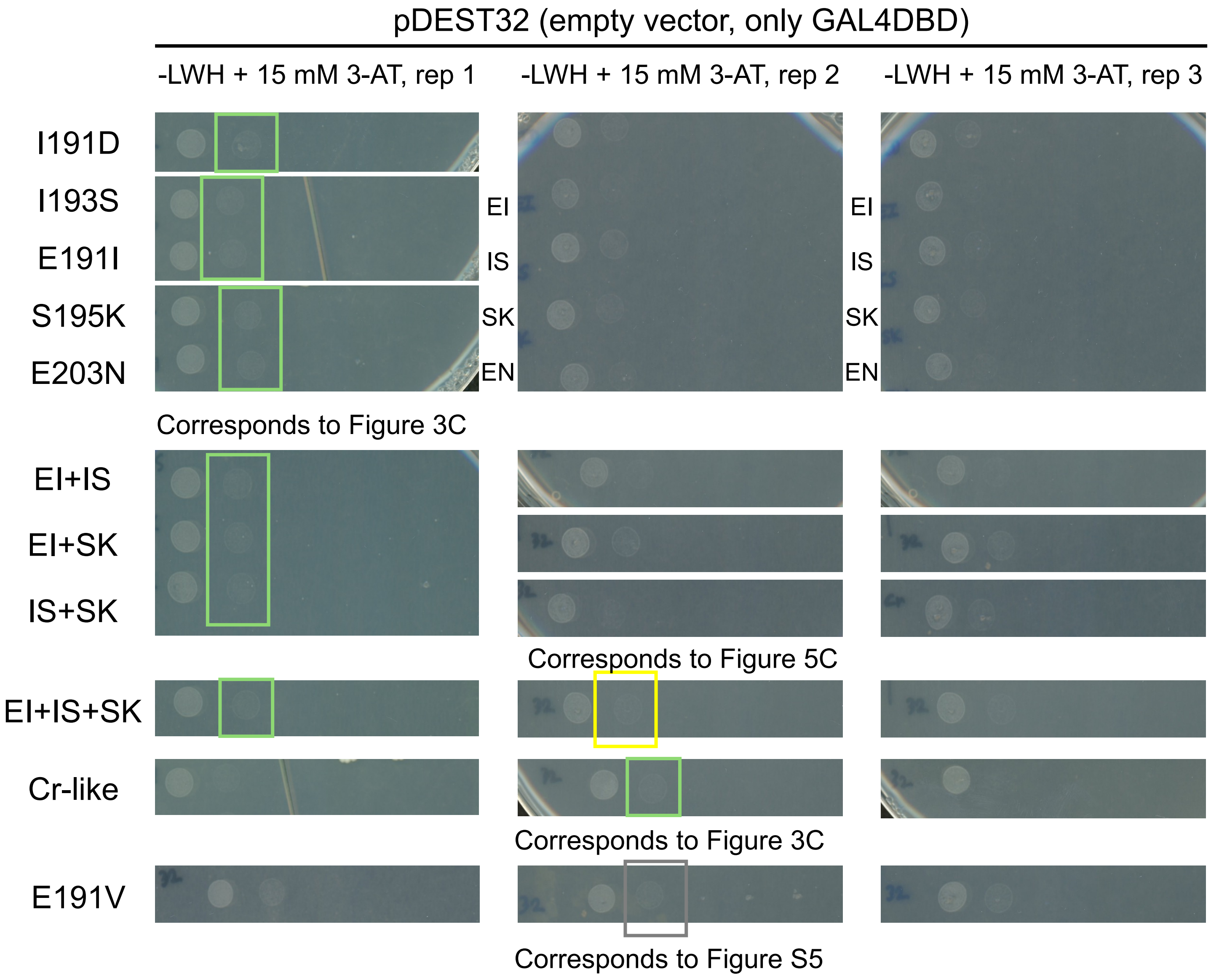


I191D; AtSGT1b$^{ID}$-pDEST22
I193S; AtSGT1b$^{IS}$-pDEST22
E191I; AtSGT1b$^{EI}$ -pDEST22
S195K; AtSGT1b$^{SK}$-pDEST22
E203N; AtSGT1b$^{EN}$-pDEST22
EI+IS; AtSGT1b$^{EI+IS}$-pDEST22
EI+SK; AtSGT1b$^{EI+SK}$-pDEST22
IS+SK; AtSGT1b$^{IS+SK}$-pDEST22
EI+IS+SK; AtSGT1b$^{EI+IS+SK}$-pDEST22
Cr-like; AtSGT1b$^{Cr\text{-}like}$-pDEST22
E191V; AtSGT1b$^{EV}$-pDEST22

(every SGT1 homolog and variant was cloned into pDEST22 (GAL4AD))

(serial dilution was started to from $OD_{600}$ = 1.0 and finished at $OD_{600}$ = 1.0 × $10^{-4}$)

(every plate was scanned three days after inoculation)

#### Data S1 – Raw data and extended yeast two-hybrid analyses

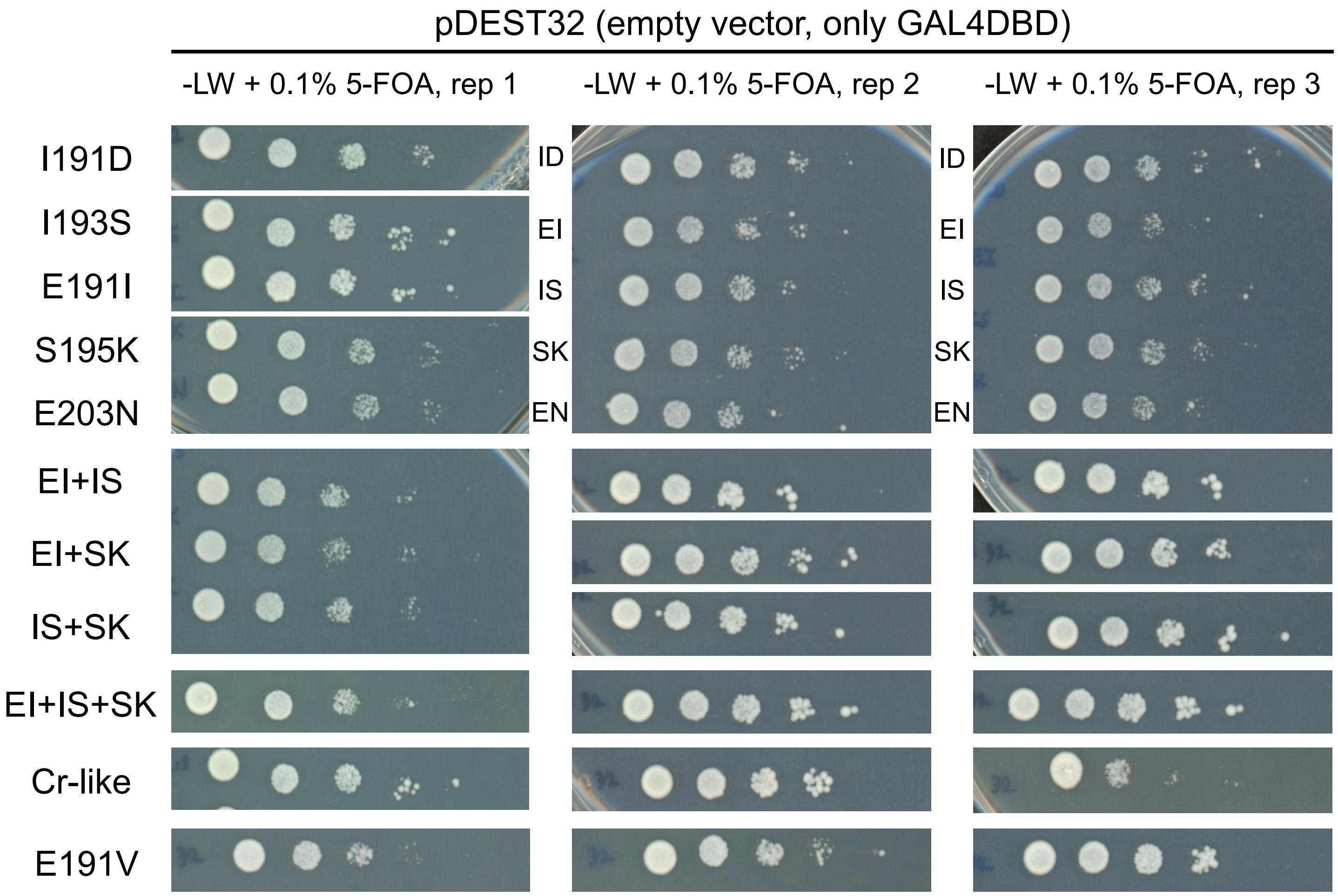


I191D; AtSGT1b$^{ID}$-pDEST22
I193S; AtSGT1b$^{IS}$-pDEST22
E191I; AtSGT1b$^{EI}$ -pDEST22
S195K; AtSGT1b$^{SK}$-pDEST22
E203N; AtSGT1b$^{EN}$-pDEST22
EI+IS; AtSGT1b$^{EI+IS}$-pDEST22
EI+SK; AtSGT1b$^{EI+SK}$-pDEST22
IS+SK; AtSGT1b$^{IS+SK}$-pDEST22
EI+IS+SK; AtSGT1b$^{EI+IS+SK}$-pDEST22
Cr-like; AtSGT1b$^{Cr\text{-}like}$-pDEST22
E191V; AtSGT1b$^{EV}$-pDEST22

(every SGT1 homolog and variant was cloned into pDEST22 (GAL4AD))

(serial dilution was started to from $OD_{600}$ = 1.0 and finished at $OD_{600}$ = 1.0 × $10^{-4}$)

(every plate was scanned three days after inoculation)

#### Data S1 – Raw data and extended yeast two-hybrid analyses

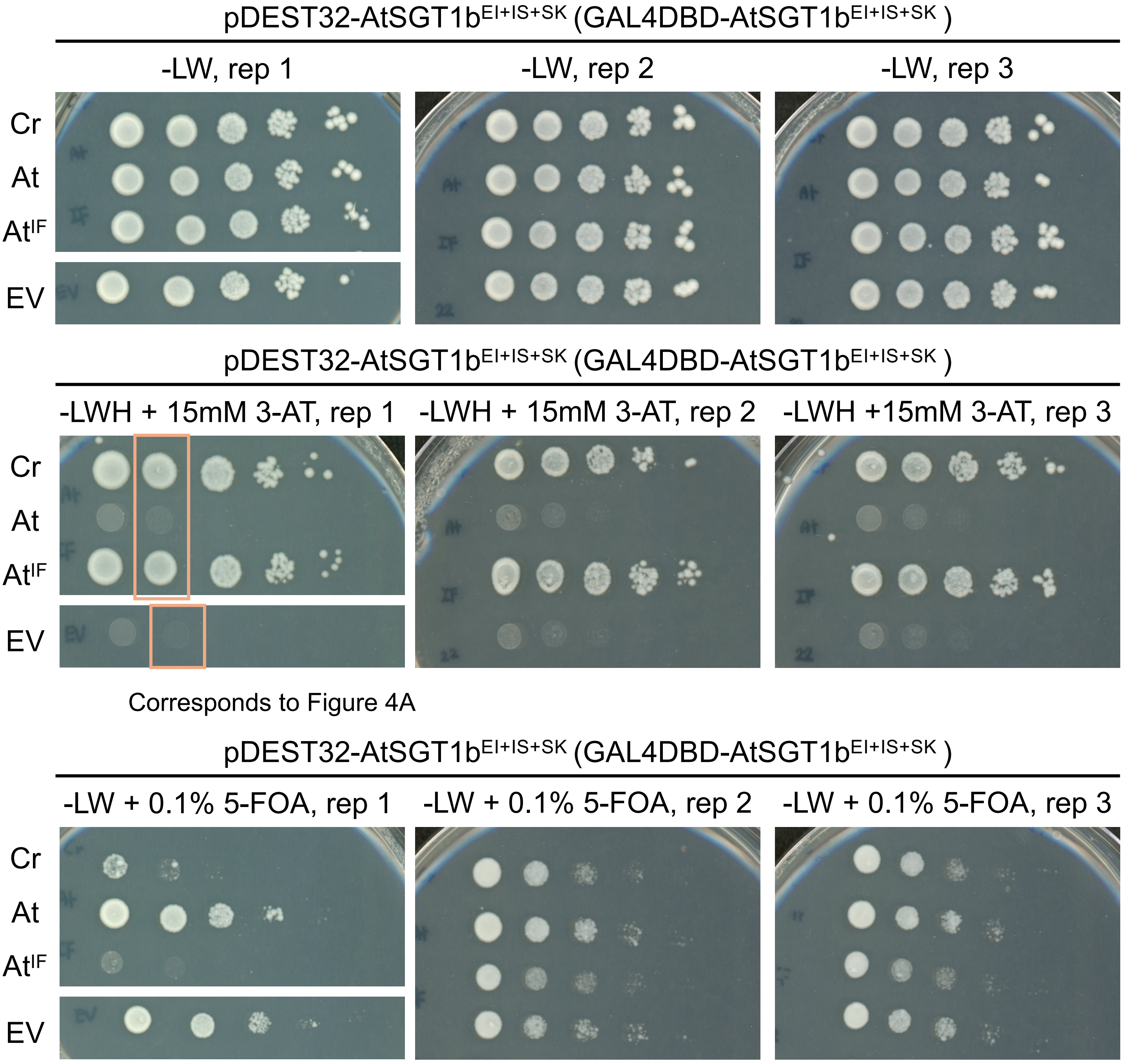


Cr; CrRAR1-pDEST22
At; AtRAR1-pDEST22
At$^{IF}$; AtRAR1$^{I153F}$-pDEST22
EV; empty pDEST22

(every RAR1 homolog and variant was cloned into pDEST22 (GAL4AD))

(serial dilution was started to from $OD_{600}$ = 1.0 and finished at $OD_{600}$ = 1.0 × $10^{-4}$)

(every plate was scanned three days after inoculation)

#### Data S1 – Raw data and extended yeast two-hybrid analyses

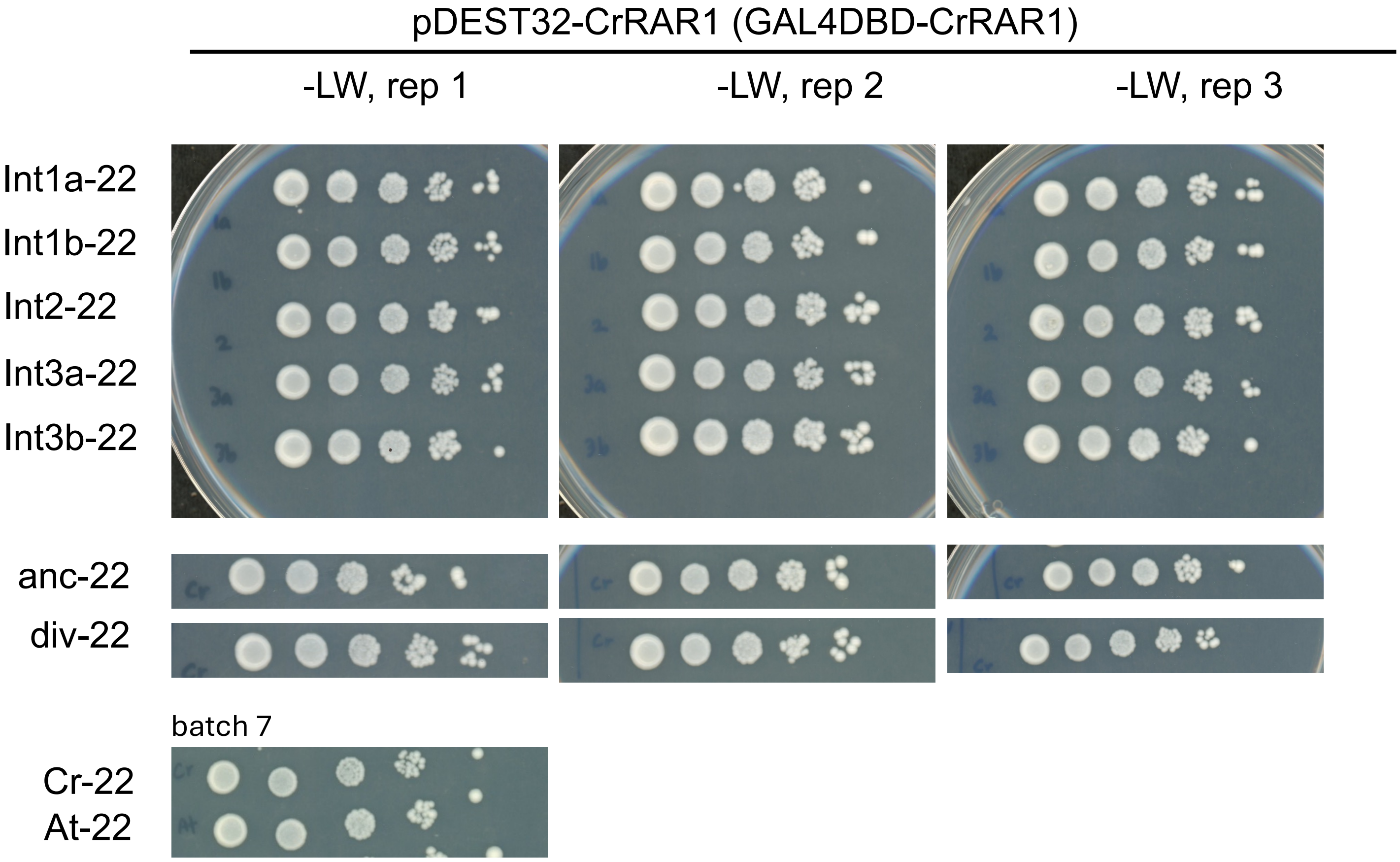


Int1a-22; SGT1-Int1a-pDEST22
Int1b-22; SGT1-Int1b-pDEST22
Int2-22; SGT1-Int2-pDEST22
Int3a-22; SGT1-Int3a-pDEST22
Int3b-22; SGT1-Int3b-pDEST22
anc-22; SGT1-anc-pDEST22
div-22; SGT1-div-pDEST22
Cr-22; CrSGT1-pDEST22
At-22; AtSGT1b-pDEST22

(every SGT1 homolog was cloned into pDEST22 (GAL4AD), except AtSGT1b (pDEST-AD-AtSGT1b))

(serial dilution was started to from $OD_{600}$ = 1.0 and finished at $OD_{600}$ = 1.0 × $10^{-4}$)

(every plate was scanned three days after inoculation)

#### Data S1 – Raw data and extended yeast two-hybrid analyses

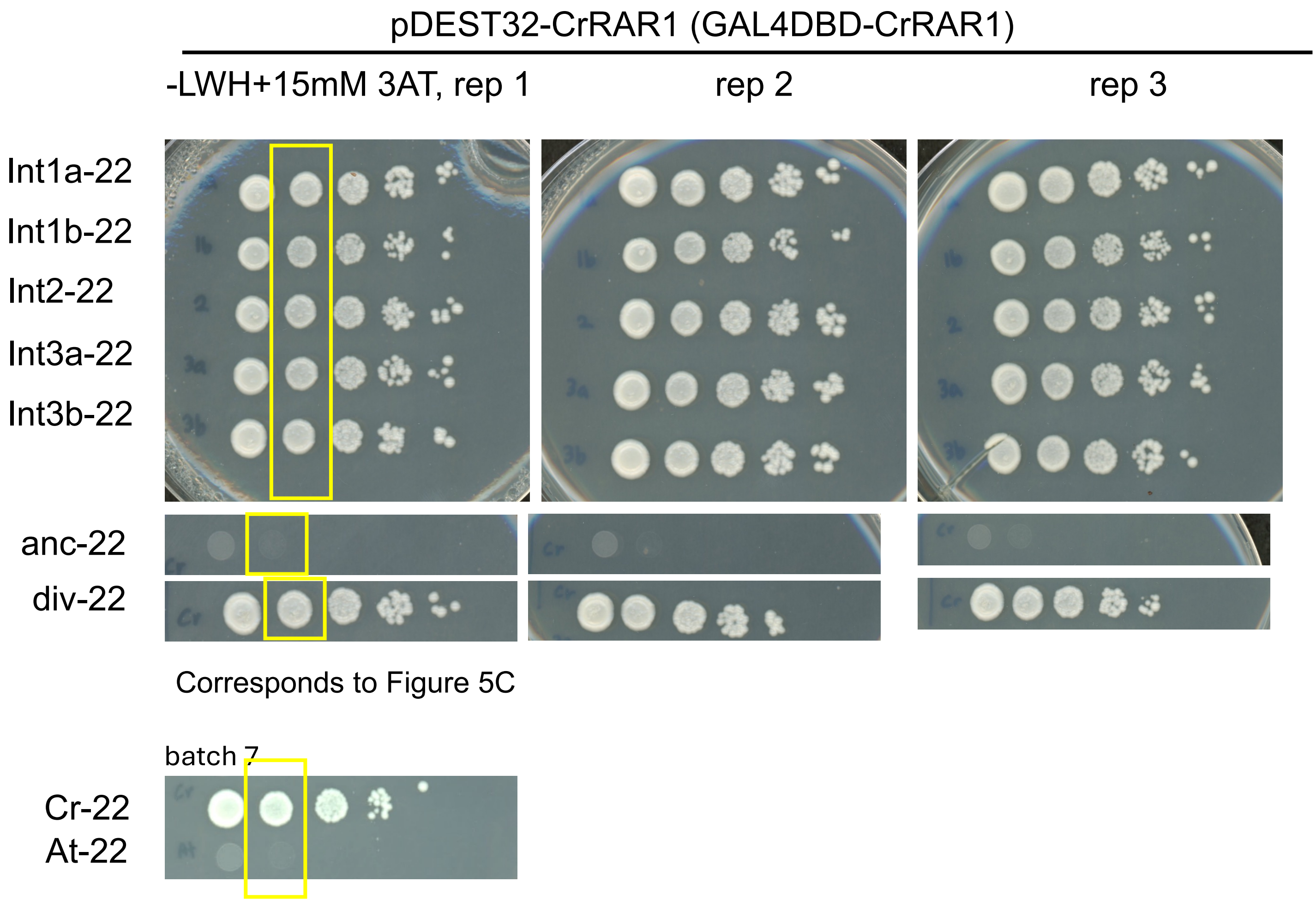


Int1a-22; SGT1-Int1a-pDEST22
Int1b-22; SGT1-Int1b-pDEST22
Int2-22; SGT1-Int2-pDEST22
Int3a-22; SGT1-Int3a-pDEST22
Int3b-22; SGT1-Int3b-pDEST22
anc-22; SGT1-anc-pDEST22
div-22; SGT1-div-pDEST22
Cr-22; CrSGT1-pDEST22
At-22; AtSGT1b-pDEST22

(every SGT1 homolog was cloned into pDEST22 (GAL4AD), except AtSGT1b (pDEST-AD-AtSGT1b))

(serial dilution was started to from $OD_{600}$ = 1.0 and finished at $OD_{600}$ = 1.0 × $10^{-4}$)

(every plate was scanned three days after inoculation)

### Data S1 – Raw data and extended yeast two-hybrid analyses

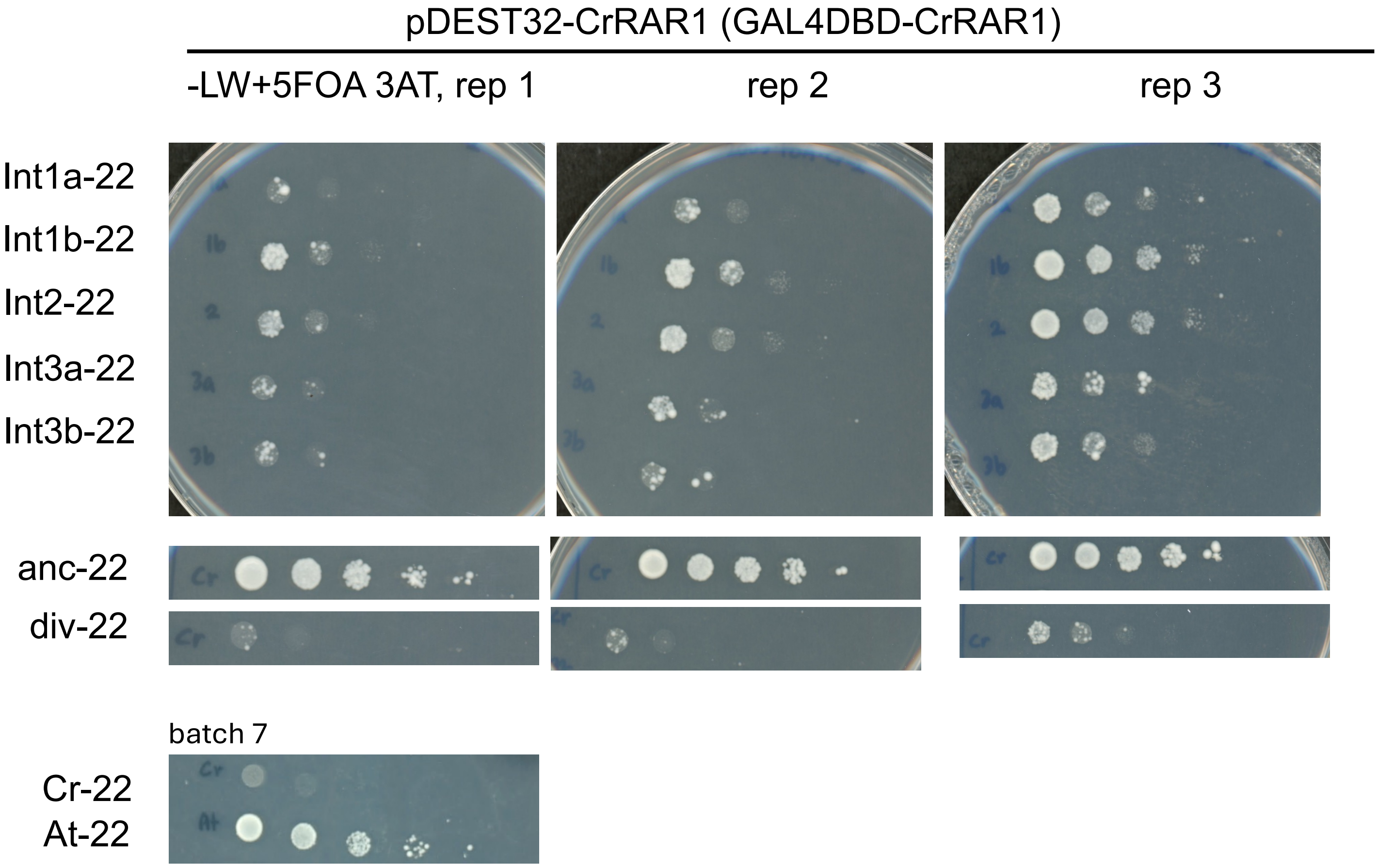


Int1a-22; SGT1-Int1a-pDEST22
Int1b-22; SGT1-Int1b-pDEST22
Int2-22; SGT1-Int2-pDEST22
Int3a-22; SGT1-Int3a-pDEST22
Int3b-22; SGT1-Int3b-pDEST22
anc-22; SGT1-anc-pDEST22
div-22; SGT1-div-pDEST22
Cr-22; CrSGT1-pDEST22
At-22; AtSGT1b-pDEST22

(every SGT1 homolog was cloned into pDEST22 (GAL4AD), except AtSGT1b (pDEST-AD-AtSGT1b))

(serial dilution was started to from $OD_{600}$ = 1.0 and finished at $OD_{600}$ = 1.0 × $10^{-4}$)

(every plate was scanned three days after inoculation)

#### Data S1 – Raw data and extended yeast two-hybrid analyses

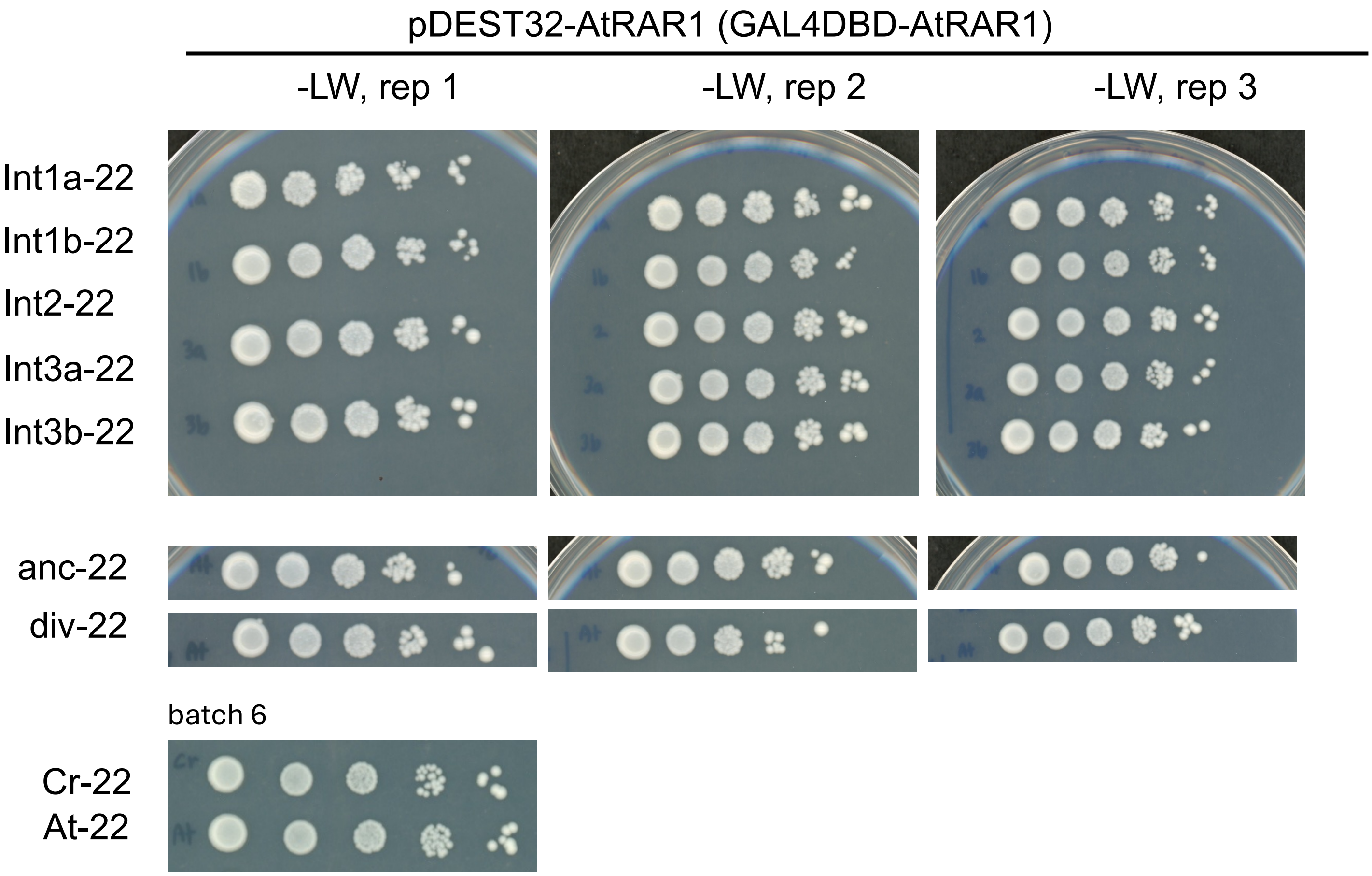


Int1a-22; SGT1-Int1a-pDEST22
Int1b-22; SGT1-Int1b-pDEST22
Int2-22; SGT1-Int2-pDEST22
Int3a-22; SGT1-Int3a-pDEST22
Int3b-22; SGT1-Int3b-pDEST22
anc-22; SGT1-anc-pDEST22
div-22; SGT1-div-pDEST22
Cr-22; CrSGT1-pDEST22
At-22; AtSGT1b-pDEST22

(every SGT1 homolog was cloned into pDEST22 (GAL4AD), except AtSGT1b (pDEST-AD-AtSGT1b))

(serial dilution was started to from $OD_{600}$ = 1.0 and finished at $OD_{600}$ = 1.0 × $10^{-4}$)

(every plate was scanned three days after inoculation)

#### Data S1 – Raw data and extended yeast two-hybrid analyses

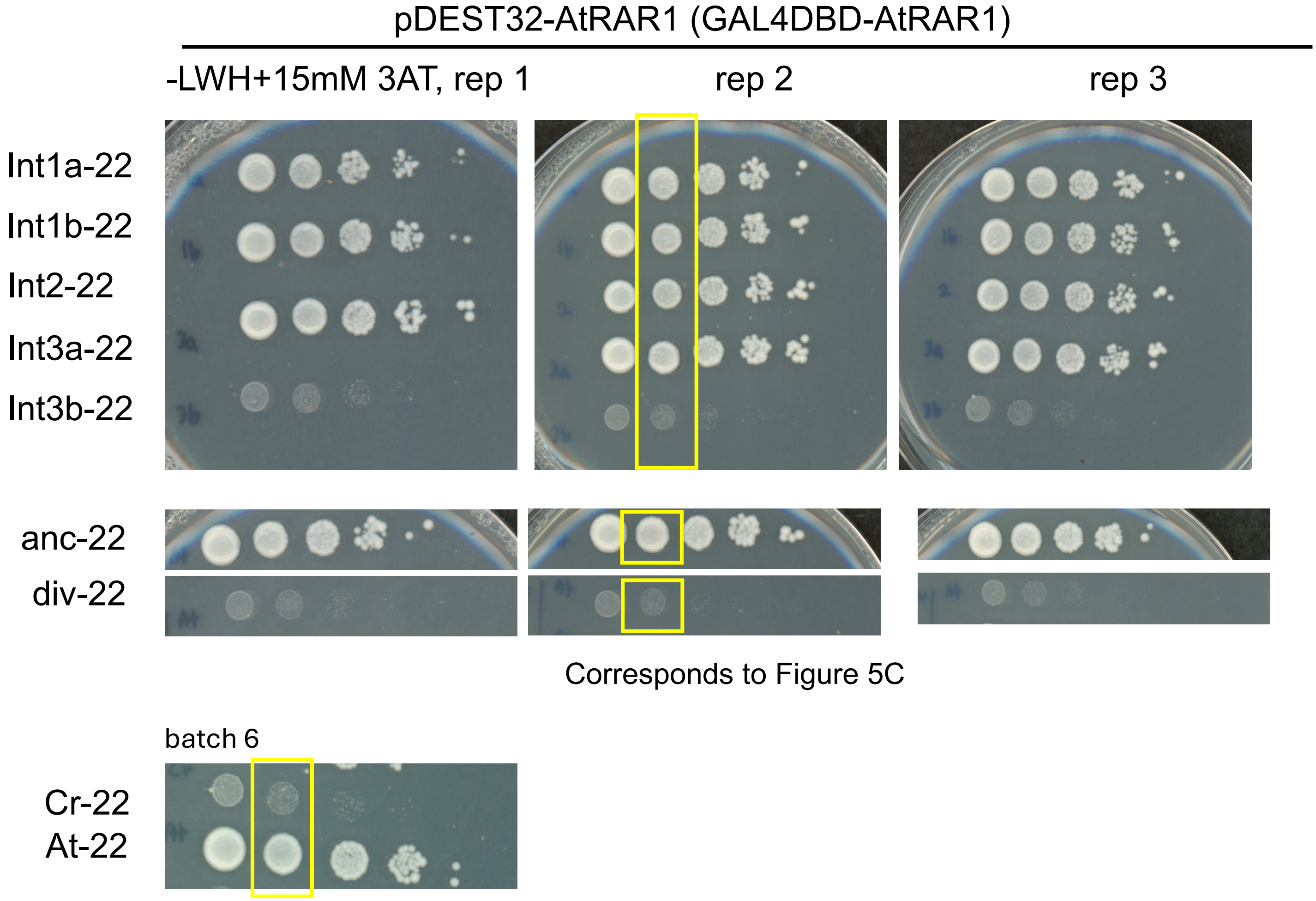


Int1a-22; SGT1-Int1a-pDEST22
Int1b-22; SGT1-Int1b-pDEST22
Int2-22; SGT1-Int2-pDEST22
Int3a-22; SGT1-Int3a-pDEST22
Int3b-22; SGT1-Int3b-pDEST22
anc-22; SGT1-anc-pDEST22
div-22; SGT1-div-pDEST22
Cr-22; CrSGT1-pDEST22
At-22; AtSGT1b-pDEST22

(every SGT1 homolog was cloned into pDEST22 (GAL4AD), except AtSGT1b (pDEST-AD-AtSGT1b))

(serial dilution was started to from $OD_{600}$ = 1.0 and finished at $OD_{600}$ = 1.0 × $10^{-4}$)

(every plate was scanned three days after inoculation)

#### Data S1 – Raw data and extended yeast two-hybrid analyses

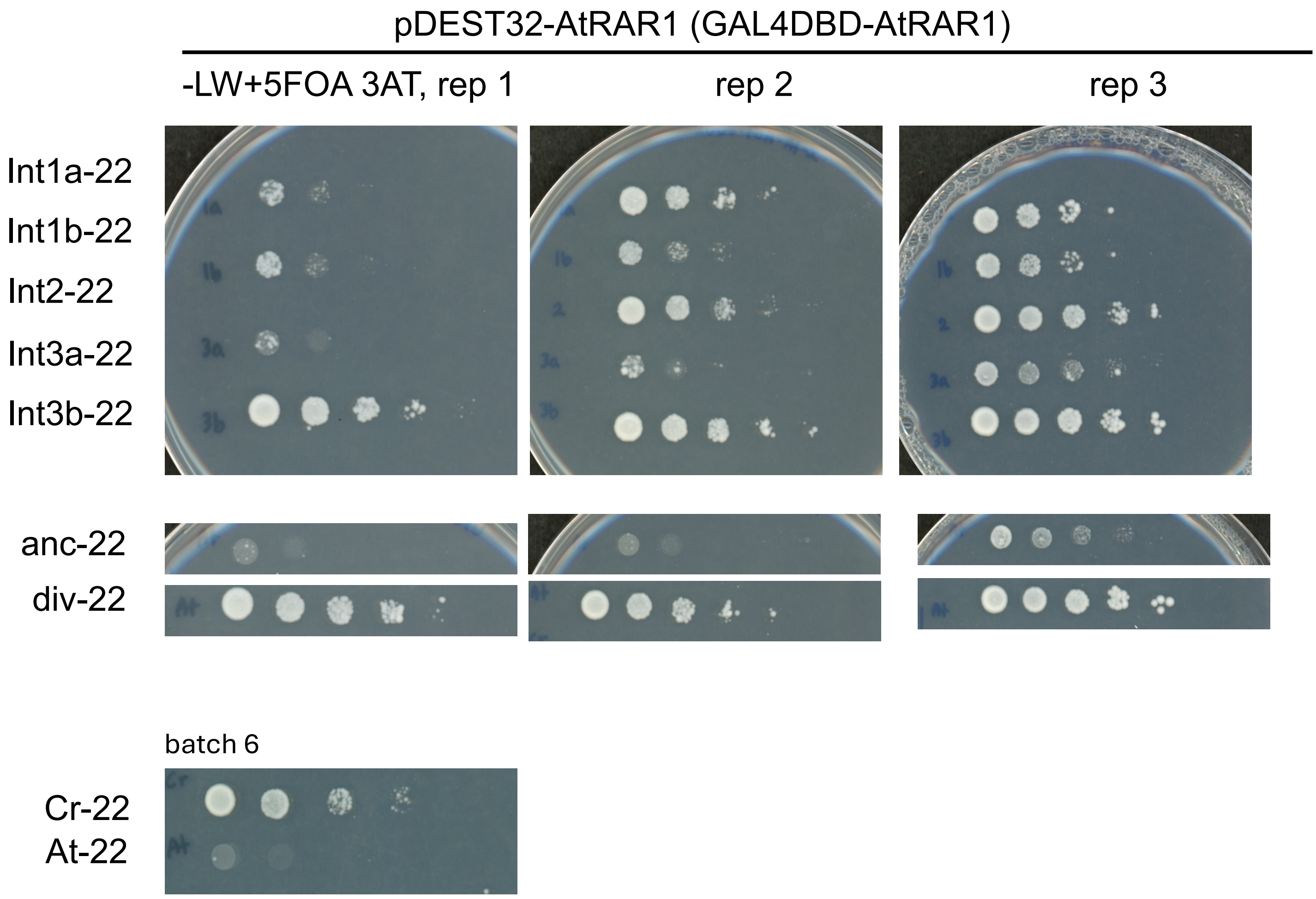


Int1a-22; SGT1-Int1a-pDEST22
Int1b-22; SGT1-Int1b-pDEST22
Int2-22; SGT1-Int2-pDEST22
Int3a-22; SGT1-Int3a-pDEST22
Int3b-22; SGT1-Int3b-pDEST22
anc-22; SGT1-anc-pDEST22
div-22; SGT1-div-pDEST22
Cr-22; CrSGT1-pDEST22
At-22; AtSGT1b-pDEST22

(every SGT1 homolog was cloned into pDEST22 (GAL4AD), except AtSGT1b (pDEST-AD-AtSGT1b))

(serial dilution was started to from $OD_{600}$ = 1.0 and finished at $OD_{600}$ = 1.0 × $10^{-4}$)

(every plate was scanned three days after inoculation)

### Data S1 – Raw data and extended yeast two-hybrid analyses

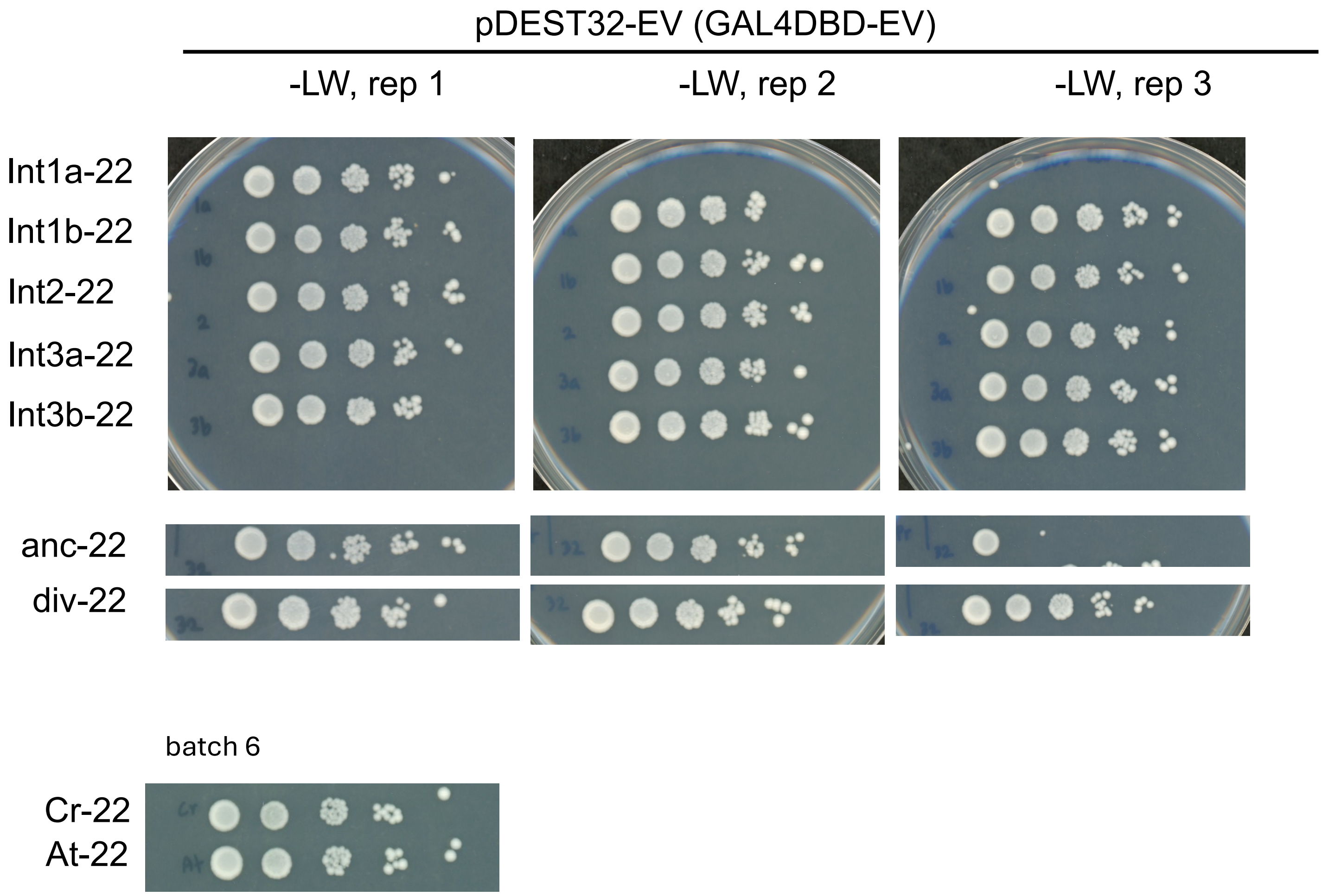


Int1a-22; SGT1-Int1a-pDEST22
Int1b-22; SGT1-Int1b-pDEST22
Int2-22; SGT1-Int2-pDEST22
Int3a-22; SGT1-Int3a-pDEST22
Int3b-22; SGT1-Int3b-pDEST22
anc-22; SGT1-anc-pDEST22
div-22; SGT1-div-pDEST22
Cr-22; CrSGT1-pDEST22
At-22; AtSGT1b-pDEST22

(every SGT1 homolog was cloned into pDEST22 (GAL4AD), except AtSGT1b (pDEST-AD-AtSGT1b))

(serial dilution was started to from $OD_{600}$ = 1.0 and finished at $OD_{600}$ = 1.0 × $10^{-4}$)

(every plate was scanned three days after inoculation)

### Data S1 – Raw data and extended yeast two-hybrid analyses

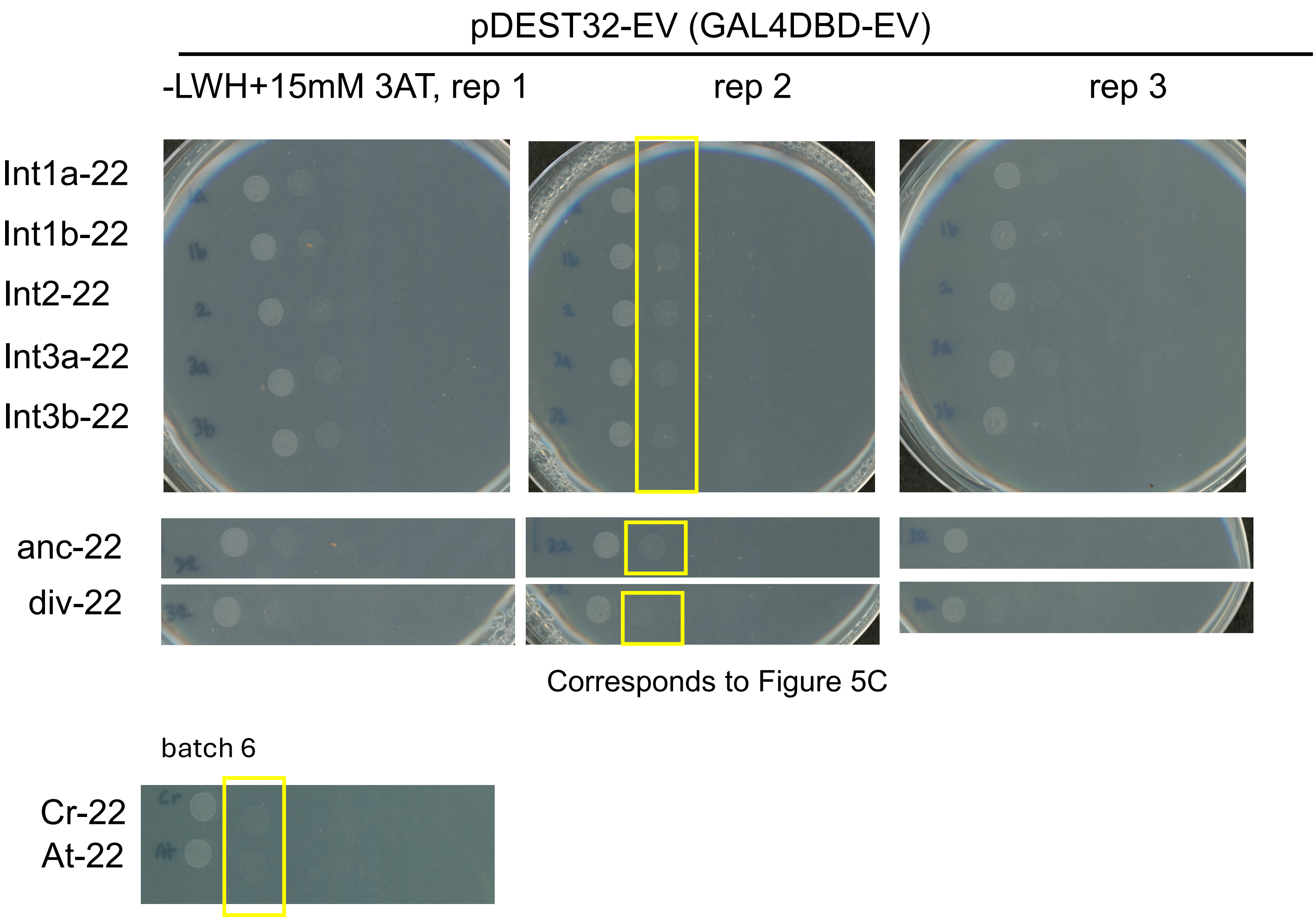


Int1a-22; SGT1-Int1a-pDEST22
Int1b-22; SGT1-Int1b-pDEST22
Int2-22; SGT1-Int2-pDEST22
Int3a-22; SGT1-Int3a-pDEST22
Int3b-22; SGT1-Int3b-pDEST22
anc-22; SGT1-anc-pDEST22
div-22; SGT1-div-pDEST22
Cr-22; CrSGT1-pDEST22
At-22; AtSGT1b-pDEST22

(every SGT1 homolog was cloned into pDEST22 (GAL4AD), except AtSGT1b (pDEST-AD-AtSGT1b))

(serial dilution was started to from $OD_{600}$ = 1.0 and finished at $OD_{600}$ = 1.0 × $10^{-4}$)

(every plate was scanned three days after inoculation)

#### Data S1 – Raw data and extended yeast two-hybrid analyses

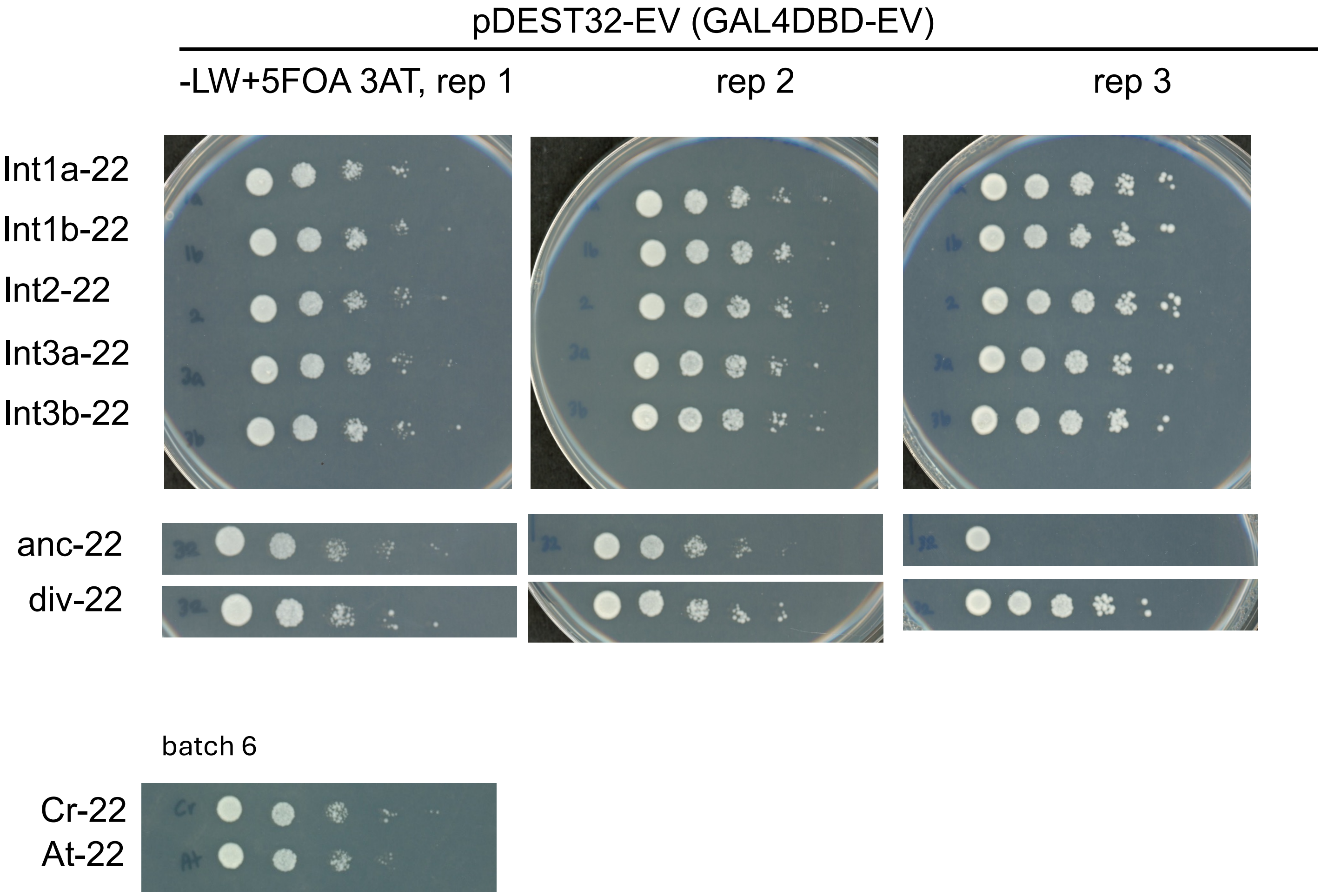


Int1a-22; SGT1-Int1a-pDEST22
Int1b-22; SGT1-Int1b-pDEST22
Int2-22; SGT1-Int2-pDEST22
Int3a-22; SGT1-Int3a-pDEST22
Int3b-22; SGT1-Int3b-pDEST22
anc-22; SGT1-anc-pDEST22
div-22; SGT1-div-pDEST22
Cr-22; CrSGT1-pDEST22
At-22; AtSGT1b-pDEST22

(every SGT1 homolog was cloned into pDEST22 (GAL4AD), except AtSGT1b (pDEST-AD-AtSGT1b))

(serial dilution was started to from $OD_{600}$ = 1.0 and finished at $OD_{600}$ = 1.0 × $10^{-4}$)

(every plate was scanned three days after inoculation)

### Data S1 – Raw data and extended yeast two-hybrid analyses

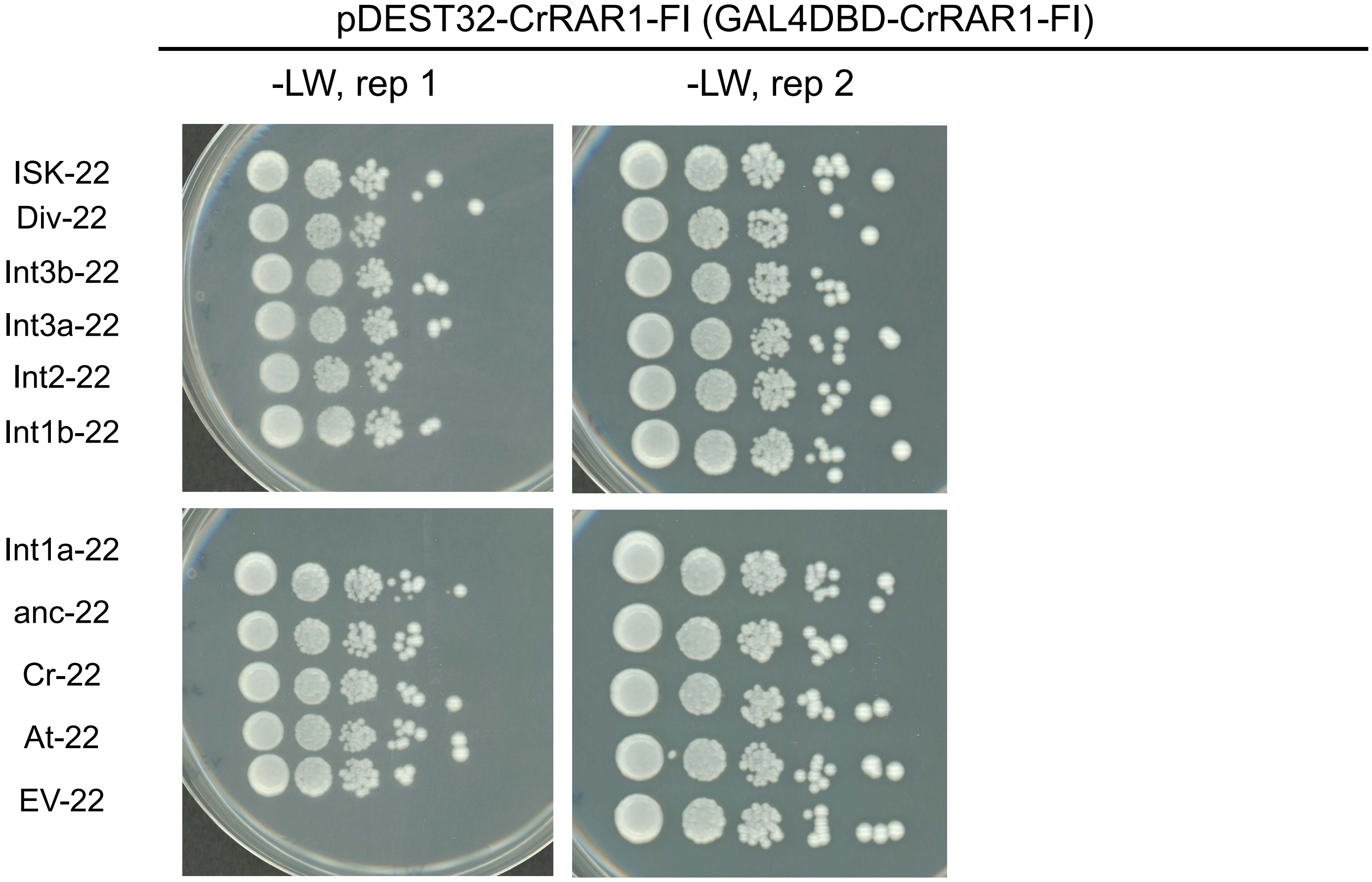


ISK-22; AtSGT1$^{ISK}$-pDEST22;
Int1a-22; SGT1-Int1a-pDEST22
Int1b-22; SGT1-Int1b-pDEST22
Int2-22; SGT1-Int2-pDEST22
Int3a-22; SGT1-Int3a-pDEST22
Int3b-22; SGT1-Int3b-pDEST22
anc-22; SGT1-anc-pDEST22
div-22; SGT1-div-pDEST22
Cr-22; CrSGT1-pDEST22
At-22; AtSGT1b-pDEST22
EV-22; empty pDEST22

(every SGT1 homolog was cloned into pDEST22 (GAL4AD), except AtSGT1b (pDEST-AD-AtSGT1b))

(serial dilution was started to from $OD_{600}$ = 1.0 and finished at $OD_{600}$ = 1.0 × $10^{-4}$)

(every plate was scanned three days after inoculation)

### Data S1 – Raw data and extended yeast two-hybrid analyses

pDEST32-CrRAR1-FI (GAL4DBD-CrRAR1-FI)

-LWH+15mM 3AT, rep 1 rep 2

ISK-22
Div-22
Int3b-22
Int3a-22
Int2-22
Int1b-22

Int1a-22
anc-22
Cr-22
At-22
EV-22

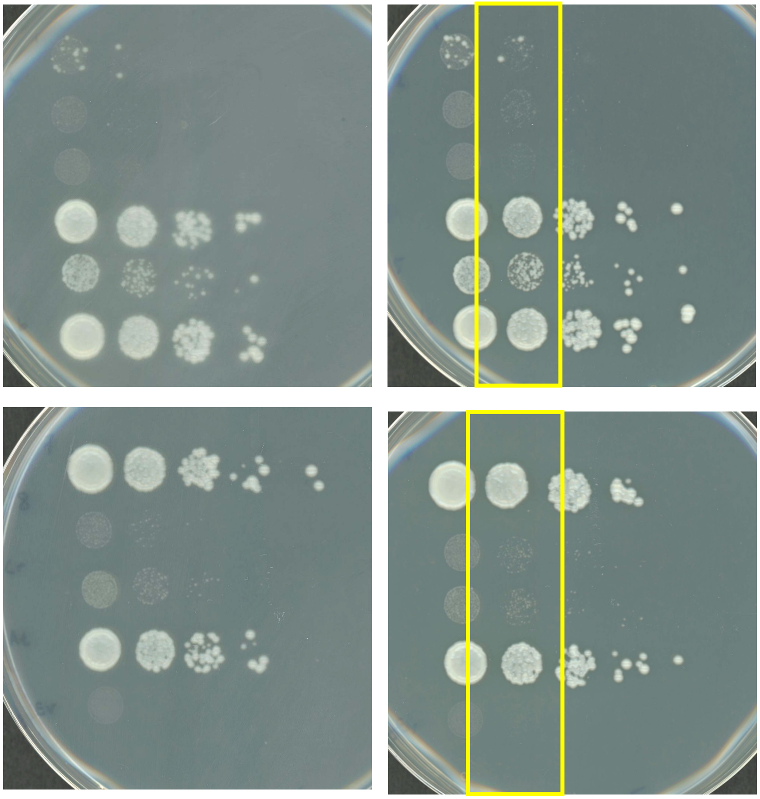

Corresponds to Figure 5C

ISK-22; AtSGT1$^{ISK}$-pDEST22;
Int1a-22; SGT1-Int1a-pDEST22
Int1b-22; SGT1-Int1b-pDEST22
Int2-22; SGT1-Int2-pDEST22
Int3a-22; SGT1-Int3a-pDEST22
Int3b-22; SGT1-Int3b-pDEST22
anc-22; SGT1-anc-pDEST22
div-22; SGT1-div-pDEST22
Cr-22; CrSGT1-pDEST22
At-22; AtSGT1b-pDEST22
EV-22; empty pDEST22

(every SGT1 homolog was cloned into pDEST22 (GAL4AD), except AtSGT1b (pDEST-AD-AtSGT1b))

(serial dilution was started to from $OD_{600}$ = 1.0 and finished at $OD_{600}$ = 1.0 × $10^{-4}$)

(every plate was scanned three days after inoculation)

#### Data S1 – Raw data and extended yeast two-hybrid analyses

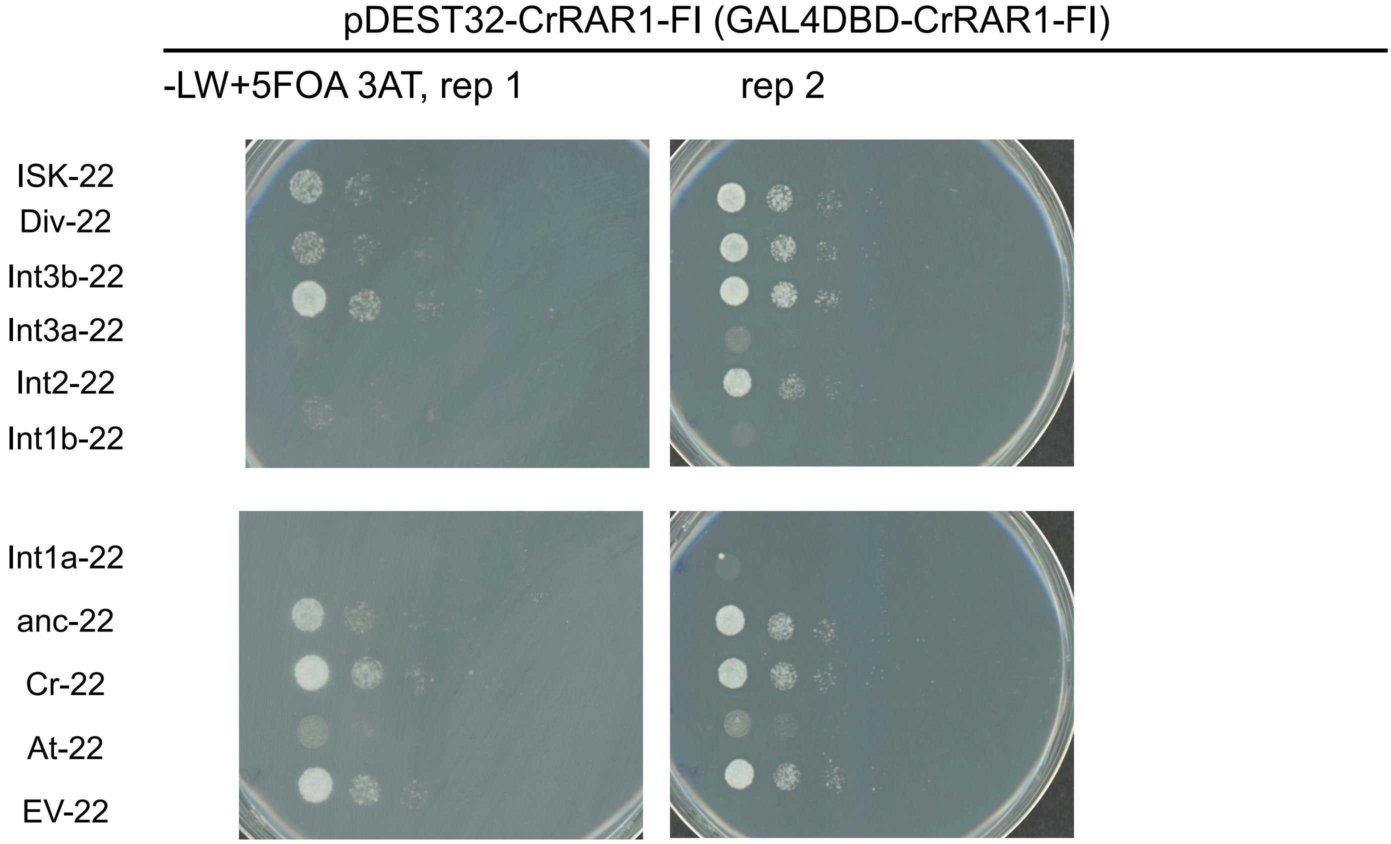


ISK-22; AtSGT1^ISK-pDEST22;
Int1a-22; SGT1-Int1a-pDEST22
Int1b-22; SGT1-Int1b-pDEST22
Int2-22; SGT1-Int2-pDEST22
Int3a-22; SGT1-Int3a-pDEST22
Int3b-22; SGT1-Int3b-pDEST22
anc-22; SGT1-anc-pDEST22
div-22; SGT1-div-pDEST22
Cr-22; CrSGT1-pDEST22
At-22; AtSGT1b-pDEST22
EV-22; empty pDEST22

(every SGT1 homolog was cloned into pDEST22 (GAL4AD), except AtSGT1b (pDEST-AD-AtSGT1b))

(serial dilution was started to from $OD_{600}$ = 1.0 and finished at $OD_{600} = 1.0 \times 10^{-4}$)

(every plate was scanned three days after inoculation)

#### Data S1 – Raw data and extended yeast two-hybrid analyses

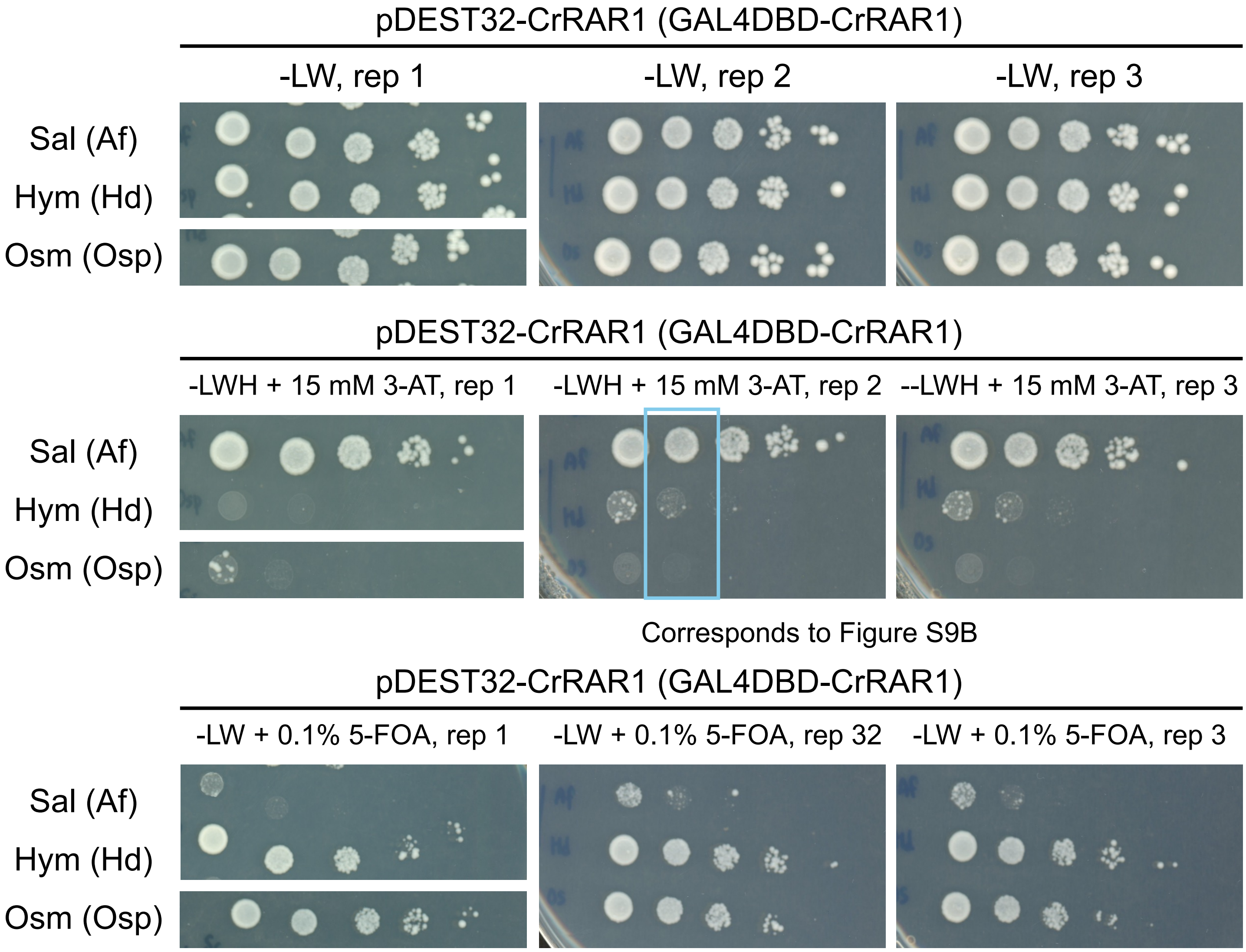


Sal (Af); AfSGT1-pDEST22
Hym (Hd); HdSGT1-pDEST22
Osm (Osp); OspSGT1-pDEST22

(every SGT1 homolog was cloned into pDEST22 (GAL4AD), except AtSGT1b (pDEST-AD-AtSGT1b))

(serial dilution was started to from $OD_{600}$ = 1.0 and finished at $OD_{600}$ = 1.0 × $10^{-4}$)

(every plate was scanned three days after inoculation)

#### Data S1 – Raw data and extended yeast two-hybrid analyses

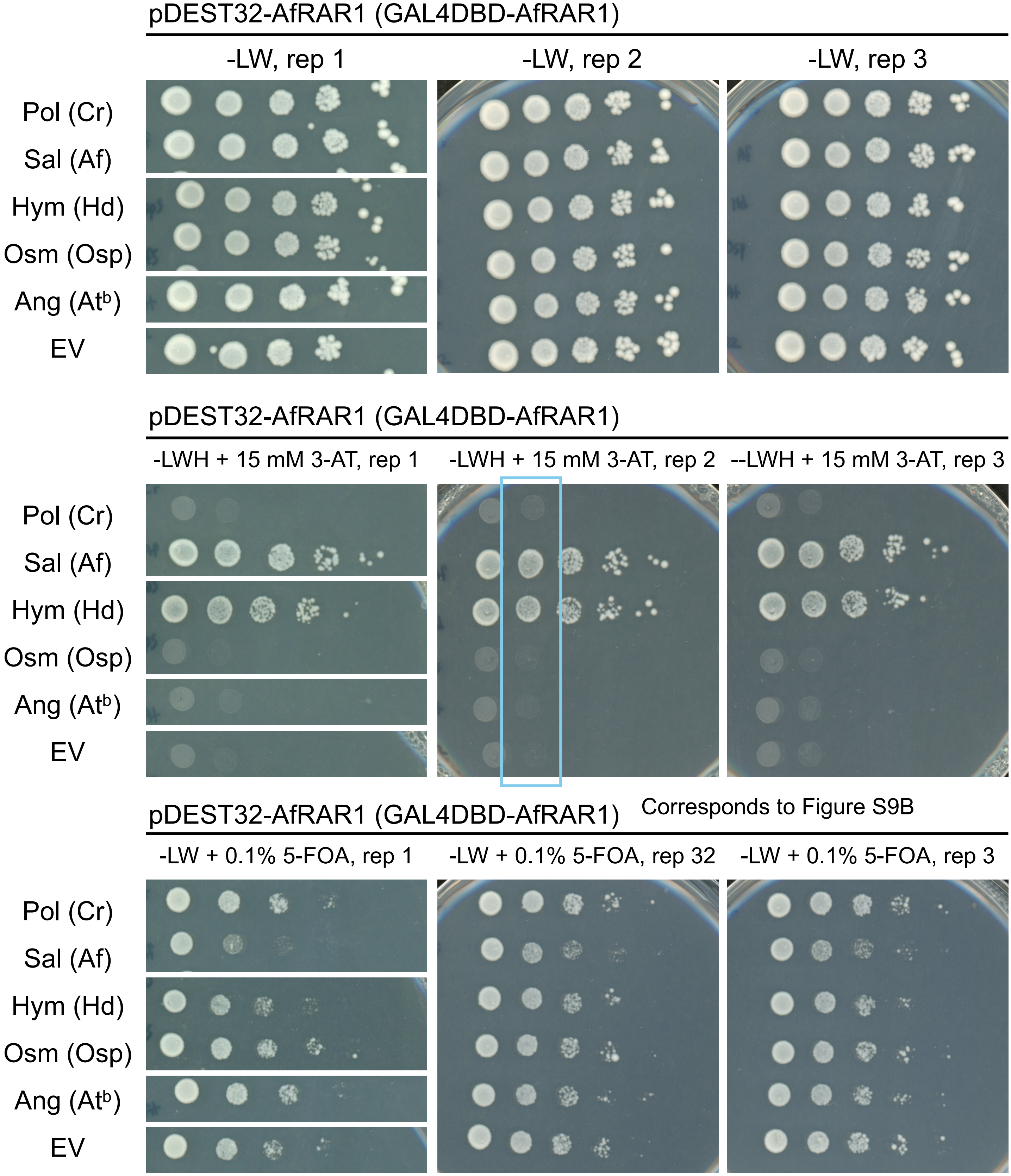


Sal (Af); AfSGT1-pDEST22
Hym (Hd); HdSGT1-pDEST22
Osm (Osp); OspSGT1-pDEST22

Pol (Cr); CrSGT1-pDEST22
Ang ($At^b$); pDEST-AD-AtSGT1b
EV; empty pDEST22

(serial dilution was started to from $OD_{600}$ = 1.0 and finished at $OD_{600}$ = 1.0 × $10^{-4}$)

(every plate was scanned three days after inoculation)

#### Data S1 – Raw data and extended yeast two-hybrid analyses

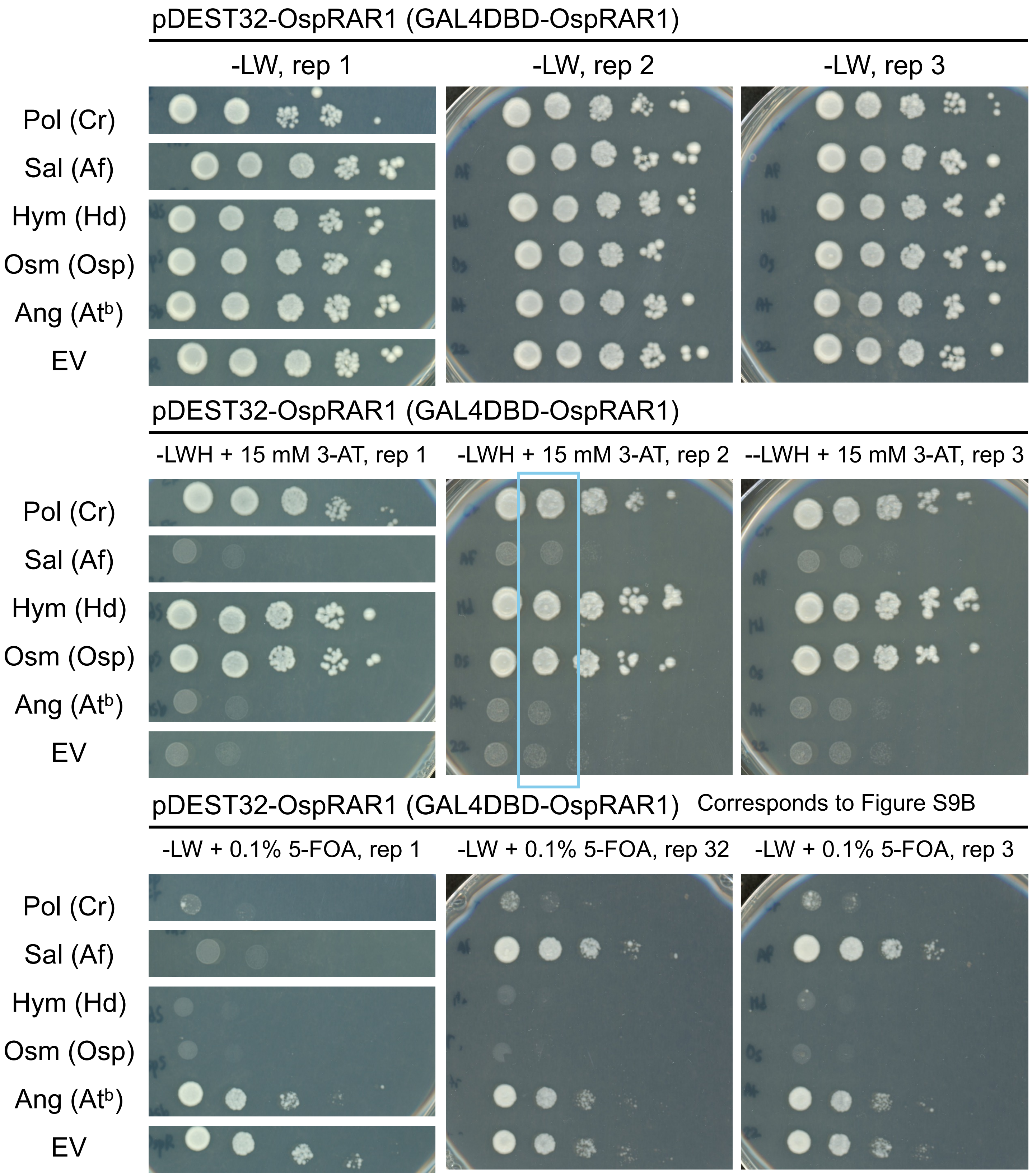


Sal (Af); AfSGT1-pDEST22
Hym (Hd); HdSGT1-pDEST22
Osm (Osp); OspSGT1-pDEST22

Pol (Cr); CrSGT1-pDEST22
Ang (At$^b$); pDEST-AD-AtSGT1b
EV; empty pDEST22

(serial dilution was started to from $OD_{600}$ = 1.0 and finished at $OD_{600}$ = 1.0 × $10^{-4}$)

(every plate was scanned three days after inoculation)

#### Data S1 – Raw data and extended yeast two-hybrid analyses

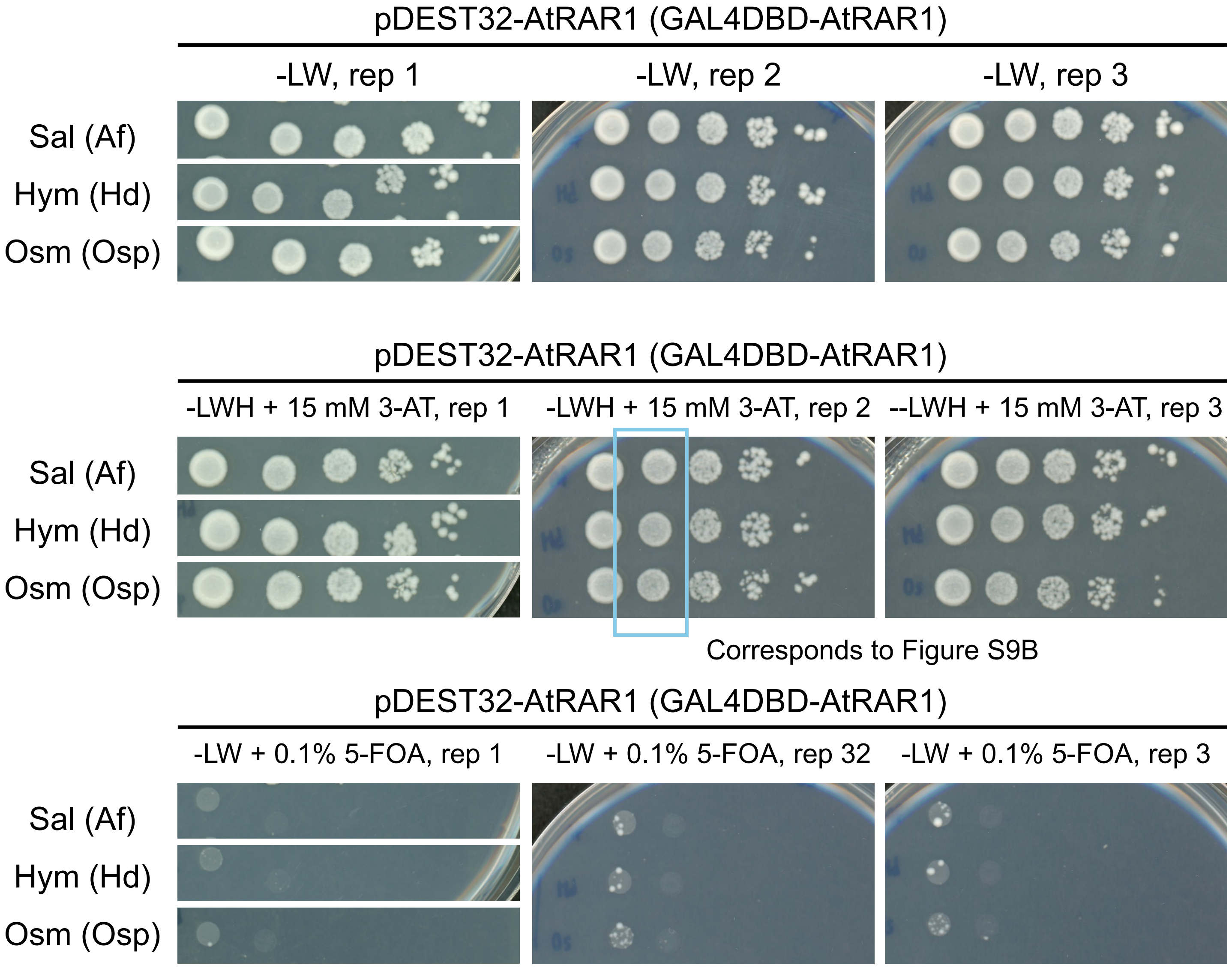


Sal (Af); AfSGT1-pDEST22
Hym (Hd); HdSGT1-pDEST22
Osm (Osp); OspSGT1-pDEST22

(serial dilution was started to from $OD_{600}$ = 1.0 and finished at $OD_{600}$ = 1.0 × $10^{-4}$)

(every plate was scanned three days after inoculation)

#### Data S1 – Raw data and extended yeast two-hybrid analyses

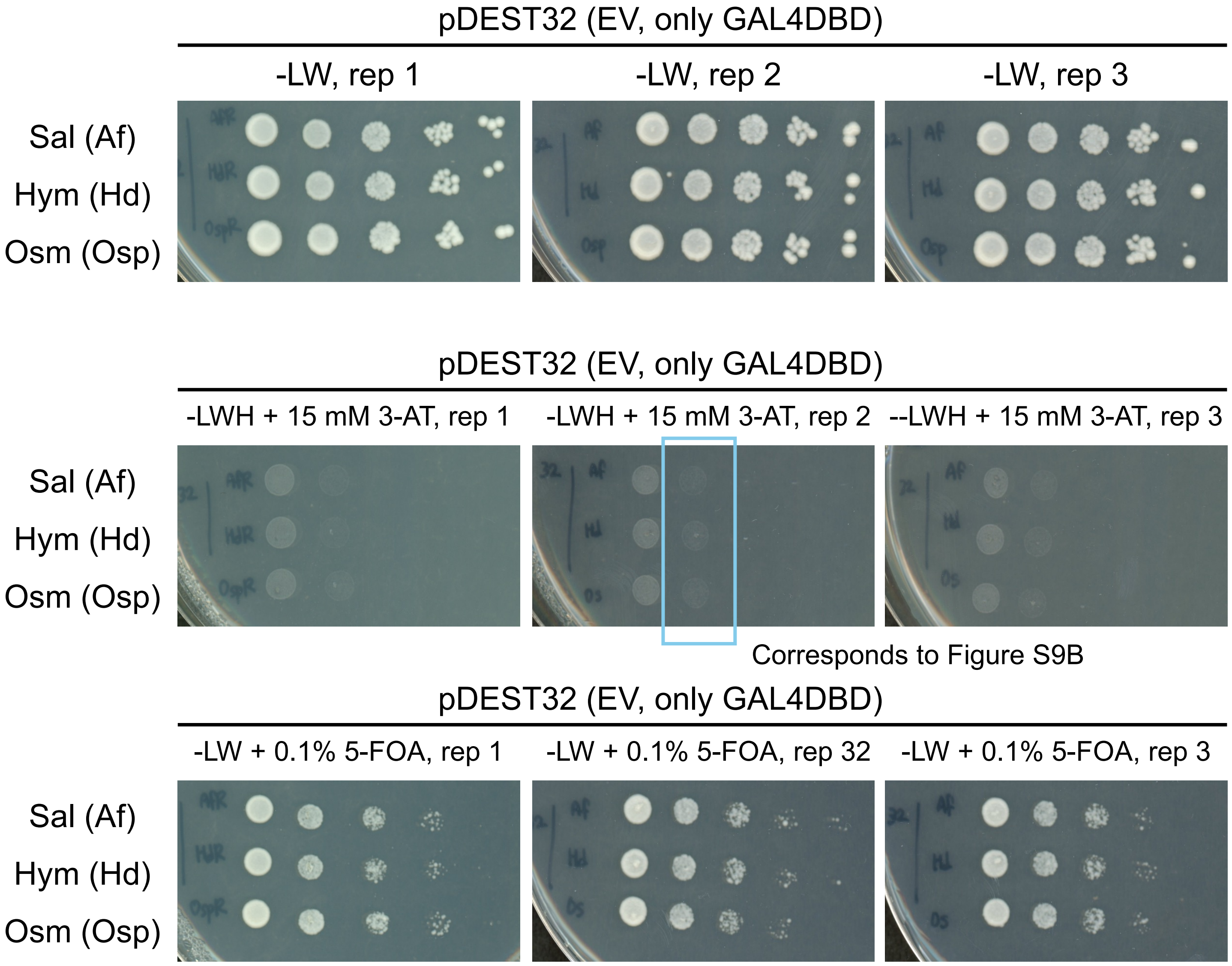


Sal (Af); AfSGT1-pDEST22
Hym (Hd); HdSGT1-pDEST22
Osm (Osp); OspSGT1-pDEST22

(serial dilution was started to from $OD_{600}$ = 1.0 and finished at $OD_{600}$ = 1.0 × $10^{-4}$)

(every plate was scanned three days after inoculation)

### Data S1 – Raw data and extended yeast two-hybrid analyses

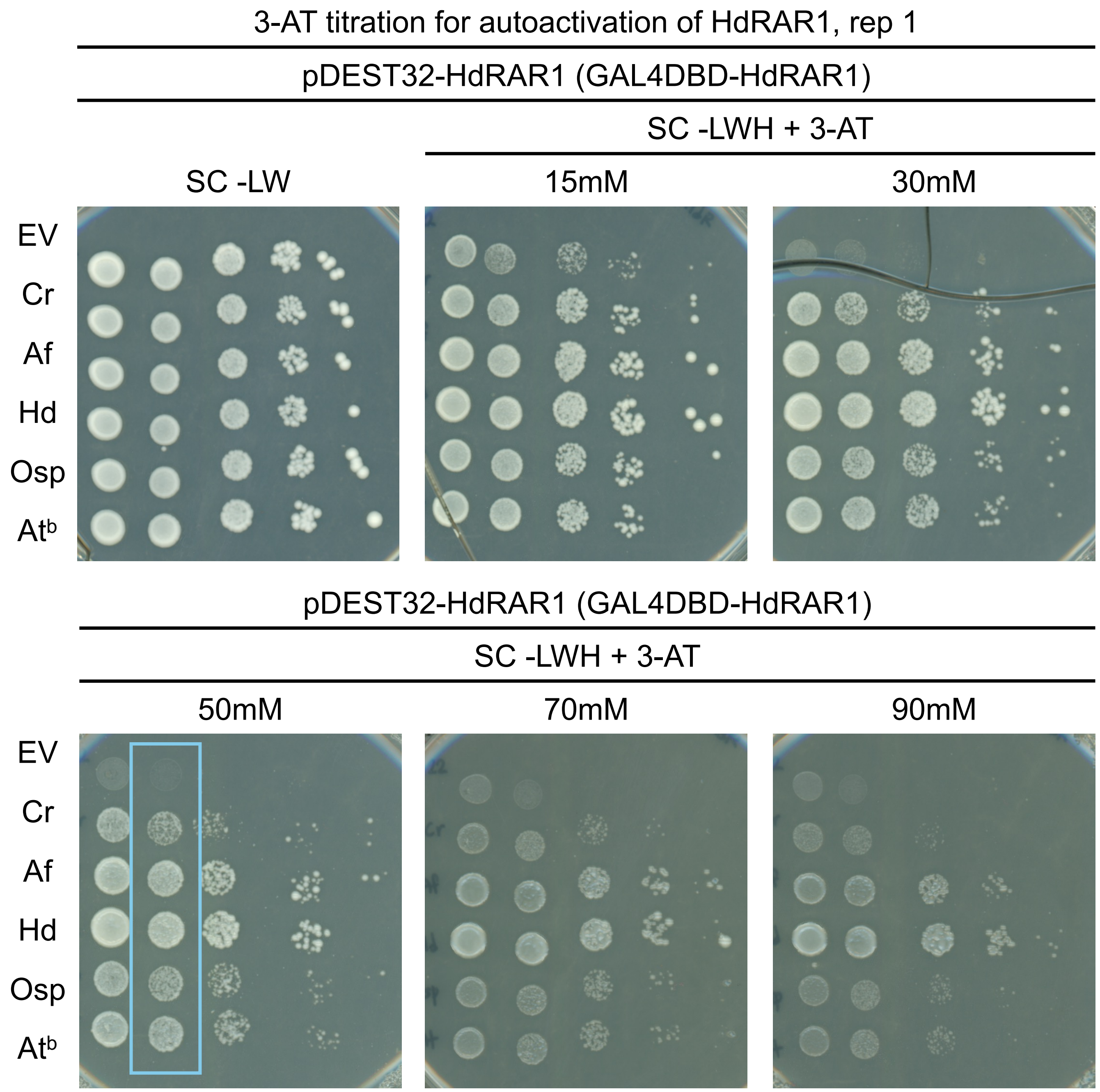


Corresponds to Figure S9B

EV; empty pDEST22
Cr; CrSGT1-pDEST22
Af; AfSGT1-pDEST22
Hd; HdSGT1-pDEST22
Osp; OspSGT1-pDEST22
At[b]; AtSGT1b-pDEST22

(serial dilution was started to from $OD_{600}$ = 1.0 and finished at $OD_{600}$ = 1.0 × $10^{-4}$)

(every plate was scanned three days after inoculation)

#### Data S1 – Raw data and extended yeast two-hybrid analyses

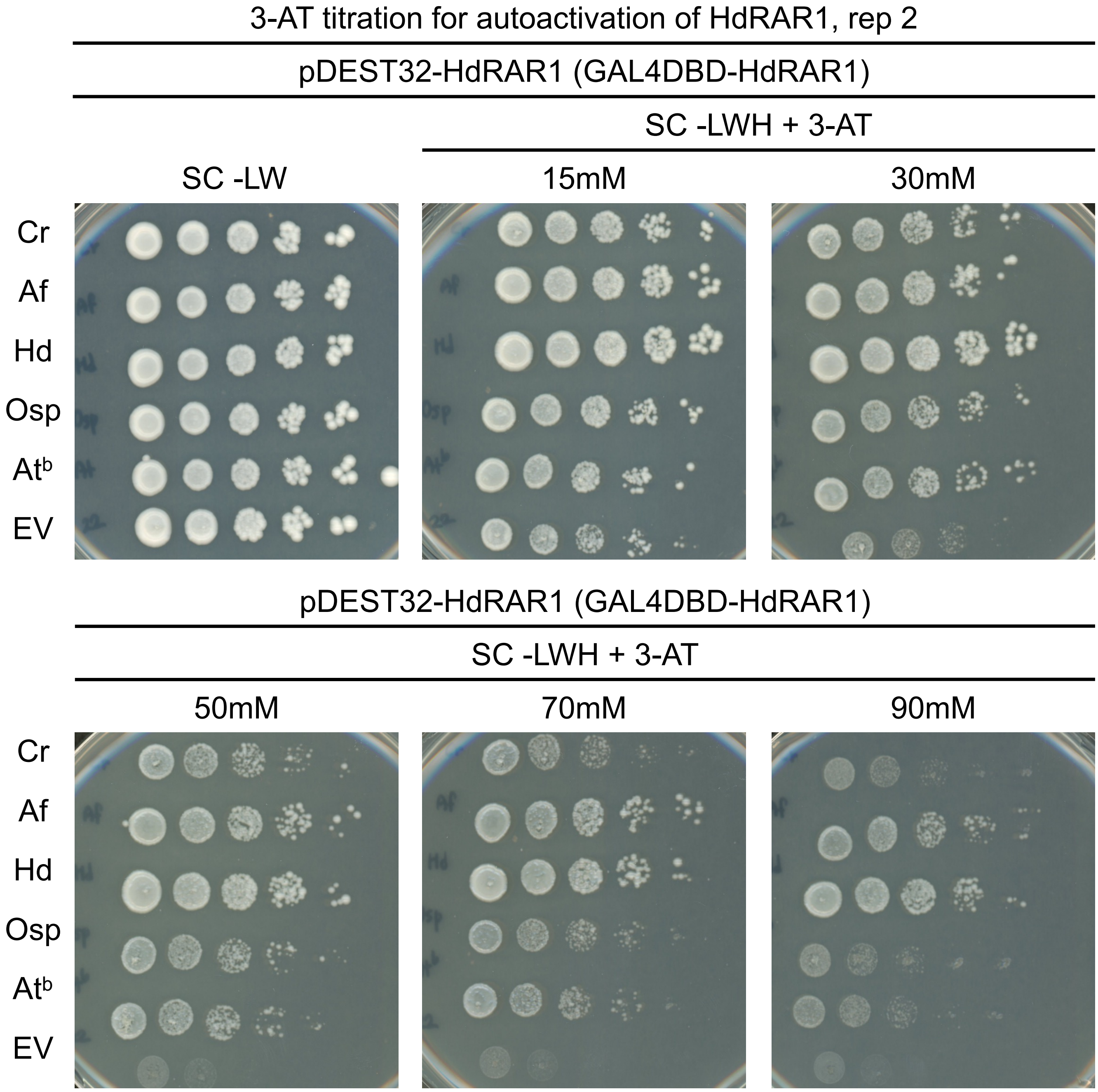


EV; empty pDEST22
Cr; CrSGT1-pDEST22
Af; AfSGT1-pDEST22
Hd; HdSGT1-pDEST22
Osp; OspSGT1-pDEST22
At$^{b}$; AtSGT1b-pDEST22

(serial dilution was started to from $OD_{600}$ = 1.0 and finished at $OD_{600}$ = 1.0 × $10^{-4}$)

(every plate was scanned three days after inoculation)

#### Data S1 – Raw data and extended yeast two-hybrid analyses

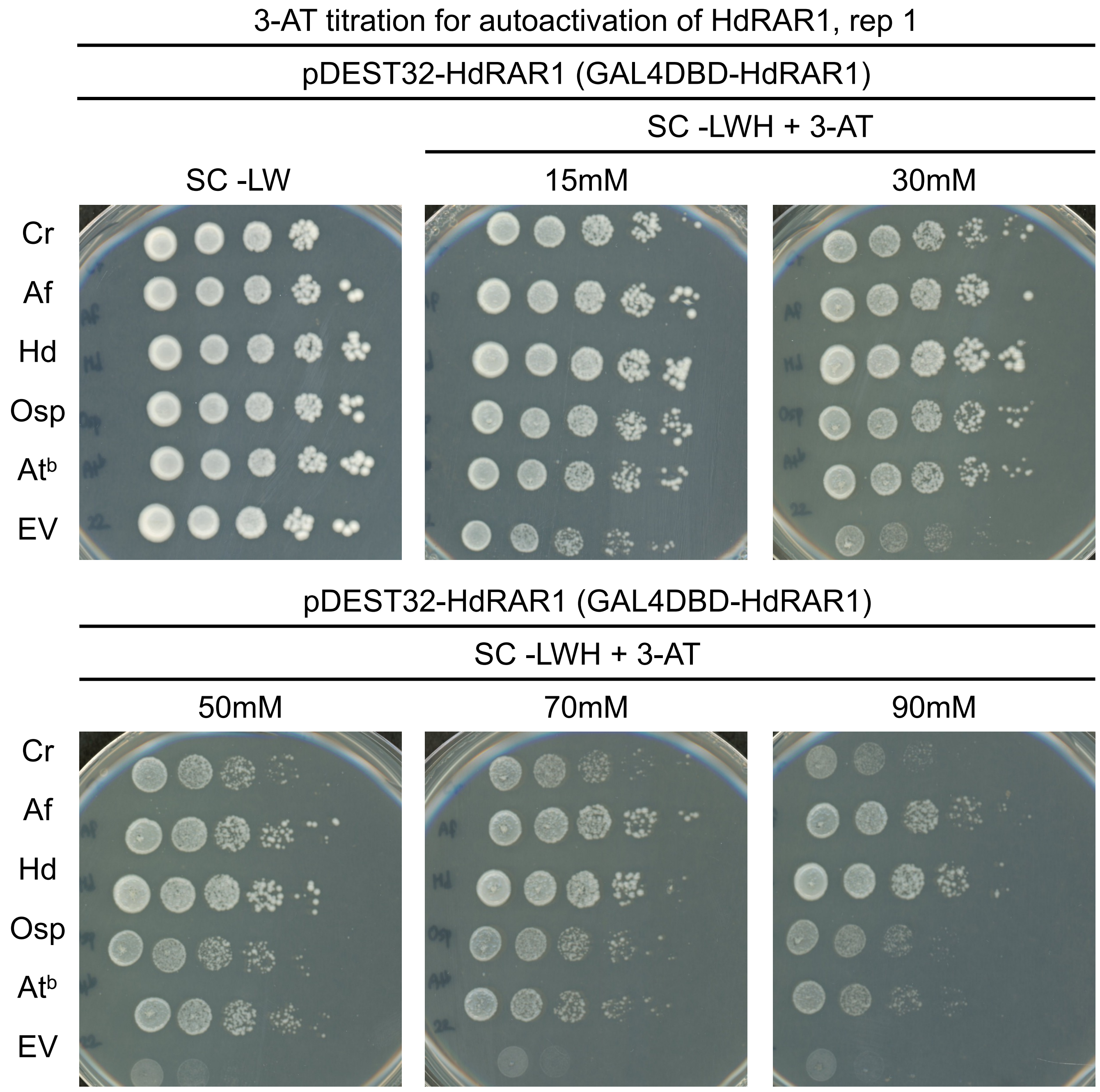


EV; empty pDEST22
Cr; CrSGT1-pDEST22
Af; AfSGT1-pDEST22
Hd; HdSGT1-pDEST22
Osp; OspSGT1-pDEST22
$At^b$; AtSGT1b-pDEST22

(serial dilution was started to from $OD_{600}$ = 1.0 and finished at $OD_{600}$ = 1.0 × $10^{-4}$)

(every plate was scanned three days after inoculation)

#### Data S1 – Raw data and extended yeast two-hybrid analyses

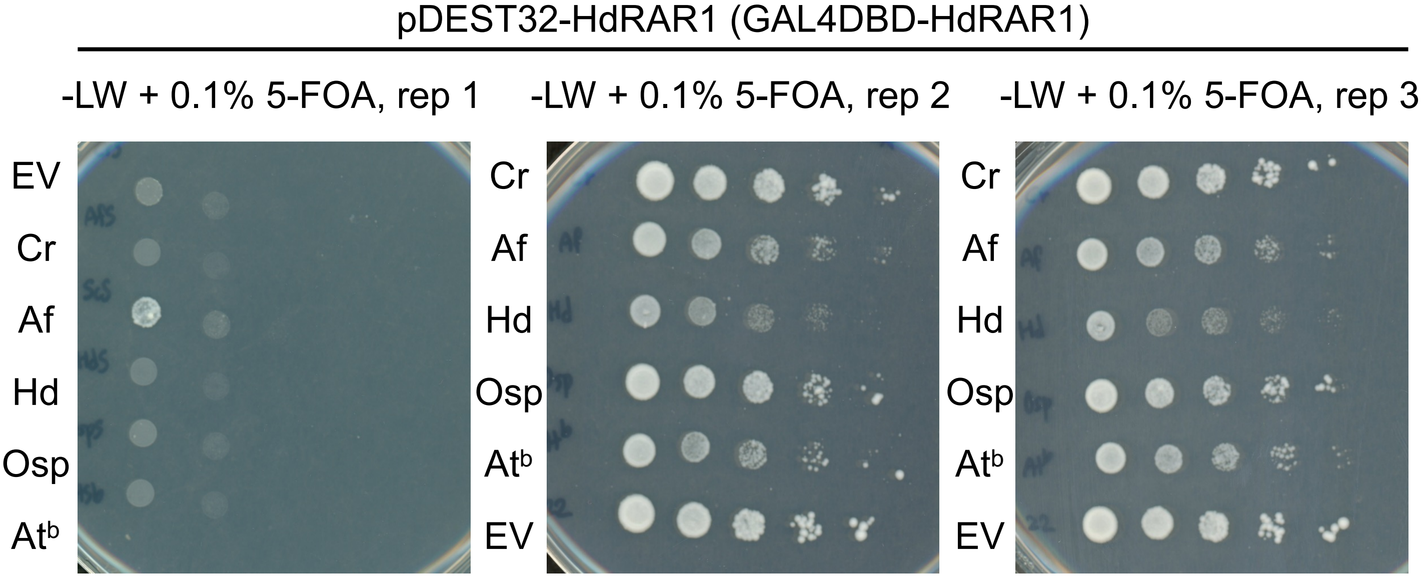


EV; empty pDEST22
Cr; CrSGT1-pDEST22
Af; AfSGT1-pDEST22
Hd; HdSGT1-pDEST22
Osp; OspSGT1-pDEST22
At[b]; AtSGT1b-pDEST22

(serial dilution was started to from $OD_{600}$ = 1.0 and finished at $OD_{600} = 1.0 \times 10^{-4}$)

(every plate was scanned three days after inoculation)
